# Hormonal gatekeeping via the blood brain barrier governs behavior

**DOI:** 10.1101/2022.12.01.518733

**Authors:** Linyang Ju, Karl M. Glastad, Lihong Sheng, Janko Gospocic, Callum J. Kingwell, Shawn M. Davidson, Sarah D. Kocher, Roberto Bonasio, Shelley L. Berger

## Abstract

Here we reveal an unanticipated role of the blood-brain-barrier (BBB) in regulating complex social behavior in ants. Using scRNA-seq we find localization in the BBB of a key hormone-degrading enzyme called Juvenile hormone esterase (Jhe), and we show that this localization governs the level of Juvenile Hormone (JH3) entering the brain. Manipulation of the Jhe level reprograms the brain transcriptome between ant castes. While ant Jhe is retained and functions intracellularly within the BBB, we show that *Drosophila* Jhe is naturally extracellular. Heterologous expression of ant Jhe into the *Drosophila* BBB alters behavior in fly to mimic what is seen in ant. Most strikingly, manipulation of Jhe levels in ant reprograms complex behavior between worker castes. Our study thus uncovers a novel, potentially conserved role of the BBB serving as a molecular gatekeeper for a neurohormonal pathway that regulates social behavior.

## Introduction

Adaptive behavioral response to environmental variation is a hallmark of animal life. Eusocial insects are exemplars for investigation of mechanisms driving behavioral plasticity, due to segregation of tasks within the colony between behaviorally, and sometimes morphologically distinct individuals that arise from a common genome (*1–3*). For example, in most eusocial insect colonies, reproductive tasks are performed by one or a limited subset of colony members, resulting in differentiation of individuals into sterile (worker) and reproductive (queen) physiological castes (*1*). Along with this classical reproductive division of labor, in many eusocial insect species there is further division of labor resulting in the allocation of distinct colony roles among worker groups (*4–6*). Multiple mechanisms underlie this phenotypic plasticity including epigenetic regulation (*7–15*), and differential regulation of hormone levels (*8, 16–20*).

Juvenile Hormone (JH3) and 20-hydroxyecdysone (20E) are the two most well-understood insect hormones. During insect development, 20E stimulates ecdysis/molting (*21–23*), while JH3 has an antagonistic role to retard ecdysis (*24, 25*). In the context of eusocial insect adult behavior these hormones are poorly understood. Associations between these hormones and eusocial insect phenotypic or behavioral castes include correlations between organismal JH3 levels and foraging or reproductive status (*8, 16, 20, 26-30*). The Florida carpenter ant, *Camponotus floridanus,* has two distinct worker castes: Major and Minor workers. Along with possessing distinct morphologies, Major and Minor workers exhibit distinct behavioral repertories. While the smaller Minors perform foraging and nursing of brood, the larger Major workers defend the nest as soldiers and only rarely forage (*7*). An important factor determining behavioral division of labor between Major and Minor workers are the levels of JH3 early in adulthood. Specifically, an epigenetic switch occurs early in adult life leading to differential regulation of several enzymes that degrade JH3, whereby JH3 degrading enzymes are repressed in Minors, leading to elevated JH3 levels and increased levels of foraging (*8*). However, it remains unknown how JH3 degradation results in neurodevelopmental outcomes that translate into distinct worker behaviors, how this is accomplished at the molecular and cellular level, and whether differential regulation of JH3 levels early in life is sufficient to accomplish lifelong Major-Minor division of labor in *C. floridanus*.

Here, using scRNA-seq, we compared single-cell transcriptomes of Major and Minor worker brains in the pupal stage and immediately after eclosion. We discovered that a JH degrading enzyme implicated in caste-specific regulation of JH3 levels (*8, 31*), JH esterase (hereafter: *cf*Jhe), displayed caste-biased expression in brain, has lost secretion from cells, and strikingly, was confined in expression to perineurial glial cells, the insect equivalent of the blood-brain barrier (BBB)(*32, 33*). Reduction of *cf*Jhe or injection of JH3 into Majors resulted in a Minor-like molecular phenotype. Transgenic expression of *cf*Jhe in *D. melanogaster* reduced foraging as in Majors, despite more than 300 million years of divergence from ants. Our results show that the BBB acts as a hormonal gatekeeper to modulate brain hormone levels leading to distinct behaviors, and that this mechanism may have mammalian analogs to attenuate brain hormone levels.

## Results

### scRNA-seq reveals neuronal and glial heterogeneity in the ant brain

Early adulthood is a key time in behavioral differentiation, including between Major and Minor workers in the ant *C. floridanus* (*7, 8*), however the underlying molecular control is not well understood. Further, behavior can be driven by a limited subset of cells within the brain (*34–36*). Hence, to dissect the brain heterogeneity between ant castes that is masked in bulk profiling methods, we carried out scRNA-seq on the central brains (*37*) of newly-eclosed (d0) Major and Minor ants (Fig. 1A), with each replicate pair taken from a distinct colony background (see methods). Filtering, integration, and clustering were performed by Seurat (*38*), and cell-types were identified using cluster-specific marker genes and visualized via UMAP-based dimension reduction (Figs. 1B, S1; Table S1), observing no gross differences in cluster representation between castes (Fig, S2A). While rigorous elucidation of cell types is best served by functional validation, we sought to contextualize our results using findings from other, better characterized systems such as *D. melanogaster*. We identified conserved neuronal (*fne+*) and glial (*repo+*) cell types, by taking advantage of single-cell RNA-seq expression markers identified in *D. melanogaster* (*39, 40*) and in the ant *Harpegnathos saltator* (*41*) (Figs. 1B-D, S1, Table S2). We performed cross-species analyses by integrating our dataset with an *H. saltator* worker/reproductive brain scRNA dataset (re-clustered using the same approach as for *C. floridanus* data, see Methods); 73% of *C. floridanus* and 67% of *H. saltator* cells could be assigned to a corresponding cluster in the alternate species, and showed strong correspondence of expression level (Fig. 1E).

**Fig 1.**
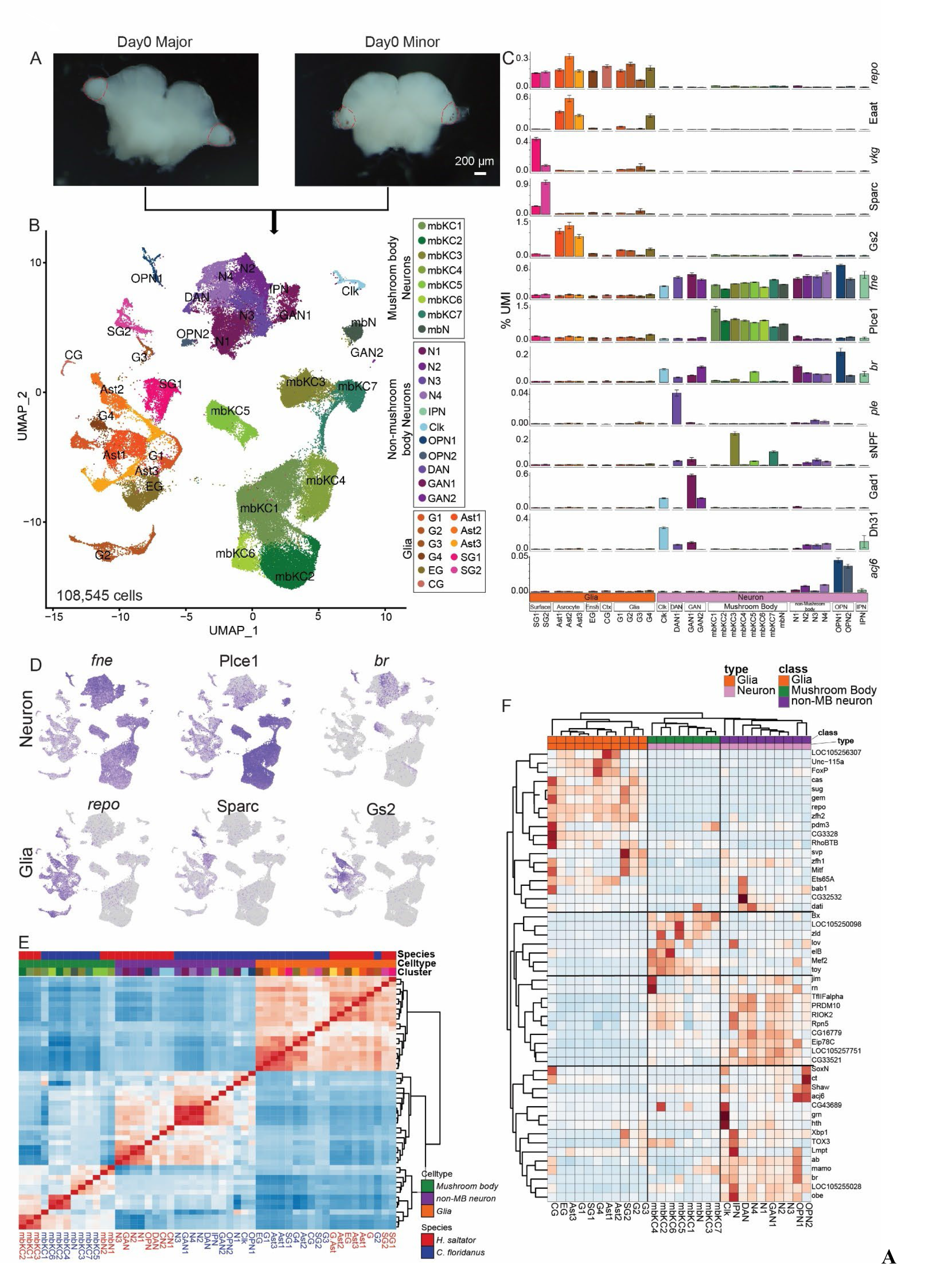
**Single-cell transcriptome profiling of adult *Camponotus floridanus* Major and Minor worker brains**. A) Widefield photo of Day0 Major and Minor non-visual brain (red dashed lines indicate optic lobe which was removed). B) Annotated UMAP visualization of integrated Major (n = 5) and Minor (n = 5) scRNA-seq data. Clusters were manually curated for known marker genes(N: Neuron, G: Glia, mbN: non-Kenyon cell Mushroom body neuron, mbKC: Kenyon cells, GAN: Gabaergic neuron, DAN: Dopaminergic neuron, OPN: Olfactory projection neuron, CN: Clock neuron, IPN: Insulin producing neuron, Ast: Astrocyte-like glia, EG: Ensheathing glia, SG: Surface glia, CG: Cortex glia). Major celltype classes are colored similarly (Mushroom body neurons: Greens; non-mushroom body neurons: purples; glia: brown, pink, orange). C) Selected marker genes for cluster annotation presented as average % total per-cell UMIs per cluster; top 5 illustrating example glia and glial-subtype marker genes, bottom eight illustrating example neuron and neuronal-subtype markers. D) Featureplot of marker genes showing per-cell expression levels of key markers for major cell groups. Neurons (fne, top left), mushroom body neurons (plce1, top middle), non MB neurons (br, top right), glia (repo, bottom left), surface glia (Sparc, bottom middle), and astrocytes (Gs2, bottom right). E) Correlation heatmap between *C. floridanus* (blue labels) and *H. saltator* (red labels) cluster expression, illustrating strong correspondence in per-cluster expression level for scRNA-seq data between species. Pearson’s correlations of mean normalized expression level per-cluster (as in Figure S1C) using all orthologous genes shared between species. F) Heatmap of all DNA binding domain containing genes showing cluster-specificity (SoupX TF-IDF > 0.75 and focal cluster/all clusters’ expression > 1.5 fold, >50% of cells in focal cluster expressing TF, adjusted p-value < 0.05; 50 genes) illustrates that expression of TFs alone is able to organize clusters into glia, Mushroom body neurons, and non-Mushroom body neurons.

Transcription factors (TFs) have crucial roles in cell differentiation during neurodevelopment and in maintaining identity of established cells after differentiation (*40, 42–44*). In support of this, when clustering our data using genes showing moderate cluster-specificity and possessing a predicted DNA-binding domain, the clustering of the major cell types was recapitulated, as were the distinct branches of mushroom body neurons (MBN), non-MB neurons, and glia (Fig. 1F).

MBNs and glial populations comprise a larger fraction of total ant brain cell types compared with solitary *melanogaster*, as previously noted in the ant *H. saltator* (*41*). Among neuronal cell types, MBNs [Phospholipase C Epsilon 1 (Plce1)+ (*45*), ∼43 % of total recovered cells) showed unique functional associations (based on Gene Ontology of marker genes) compared with non-MBNs (Fig. S2B), and top markers genes contributing to enriched terms support these association (S2B, right). For example, “synaptic transmission, cholinergic”, and “phospholipase C-activating GPCR signaling” were enriched in MBN, whereas “ionotropic glutamate receptor signaling pathway”, “sodium ion transmembrane transport” and “positive regulation of circadian sleep/wake cycle” were enriched in non-MBN, which also displayed differential extra-cellular and intra- cellular signaling functional enrichment compared to MBN (Fig. S2B). Further dissection of neuronal populations led to identification of 19 subclusters (Fig. 1B), according to the expression of genes involved in neurotransmitter production and detection (cholinergic: VAChT, dopaminergic: *ple*, glutaminergic: VGluT, GABAergic: Gad1, peptidergic: sNPF, Allatostatin A) (Figs. 1C and S2E), which were also found in *D. melanogaster* and in mammalian brain datasets (*40, 46*). Using established markers we identified olfactory projection neurons (OPN1, OPN2, *acj6+*)(*44*), insulin producing neurons (IPC, Ilp2+)(*47*), and clock neurons (Clk, Dh31+) ((*48*); Fig. 1C). In MBN we found Kenyon cells, the main component of the mushroom body, and these displayed a diverse clustering pattern (Fig. 1B, green clusters), as found previously (*49, 50*) and overlapped well with data from *H. saltator* (*41*)(Figs. 1E, S3A and S3B). Among marker genes in MBN clusters, we identified a series of conserved TFs, such as *dati*, *rn, jim* and *lov*, which showed consistent cell type distribution between species (Fig. S3B), as did multiple other neuronal and glial cell type markers (Fig. S3C). Interestingly, one Kenyon cell subtype, mbKC5, was not present in the existing *H. saltator* clustered data; cluster-specific TFs *Kruppel* (*Kr*)(*51*) and *Zelda* (*52, 53*) (Fig. S3D) illustrate that mbKC5 may be a population of newly-born or actively differentiating neurons unique in early adult (*C. floridanus*, day 0) but not in mature adult stage (*H. saltator*, day 30)(*41*).

In contrast with neurons, glial cell types showed overall better separation as illustrated by our UMAP plot (Fig.1B) due to distinct transcriptional profiles, which likely correspond to distinct functions. Based on genetic markers and reciprocal cross-species analyses, we annotated astrocyte-like (Eaat, Gs2), ensheathing (Idgf4), cortex (*wrapper*), and surface glia (*vkg*, SPARC), which are the four major glial cell-types previously identified in the insect central brain (*40, 41, 54*) (Fig. 1B). Similar with features commonly found in insect and mammalian glial single-cell datasets indicating specific subfunctions (*40, 41, 55, 56*), we observed clear heterogeneity within individual glial cell types, such as Astrocyte-like cells (Ast1, Ast2, Ast3), Surface Glia (SG1, SG2) and Ensheathing Glia (EG). Clustering based entirely upon enrichment among GO terms identified using cluster maker genes recapitulated stratification of the major cellular types (MB, non-MB and glia) (Fig. S4). In addition to the clear separation and different GO terms in MBN, non-MBN, and glial cells, we identified potential distinct functions in each subcluster of the same cell type. For example, among three astrocyte-like glia clusters (Ast1-3), “drug transport”, “neurotransmitter reuptake” and “amino acid import” are enriched, respectively (Fig. S4, full list in Table S5).

We also performed gene regulatory analysis using SCENIC (*57, 58*). We did not find that TF regulons (regulatory units) well-clustered our data (e.g. (*42*)), likely because due to lack of homology between *D. melanogaster* DNA binding motifs used to annotate *C. floridanus* gene promoters (see Methods) across 300 million years of divergence. However, we identified several putative regulatory networks associated with key marker genes for each major cell group (Fig. S5, Table S3, see Methods), including the TF *twin of eyeless* (*toy*) for Mushroom body neurons (Fig S5), which is consistent with reports in *Drosophila* where *toy* is a regulator of embryonic and post-embryonic mushroom body development and patterning (*59*). For glia, the top TF-linked regulon was regulated by the well-known glial marker *reverse polarity* (*repo*; Fig S5), and *broad* (*br*) for non-Mushroom body neurons (Fig S5). Each of these was linked in co-expression to multiple downstream TFs that showed more specific expression within these major cell groups (Fig. S5).

We validated the relevance of the scRNA dataset in assessing caste-specific patterns of gene expression by comparing to previous bulk RNA-seq data (*8*). Notably, caste-biased differentially expressed genes (DEGs) in the bulk RNA-seq demonstrated significant overlap with per-cluster caste scDEGs between Major and Minor for many scRNAseq clusters (Fig. S6A). Among these were genes previously associated with social insect caste including the Minor-biased Ftz-f1 (*60*), ari-1 (*61*), and ttk (*8*), and Major-biased Jheh (*8*), AstA-R1 (*62*) and *ciboulot* (*cib*)(*63*) (Table S4). Interestingly, Major-biased expression of caste-specific genes showed stronger enrichment among glial cell types, while Minor-biased genes were more strongly enriched among MBNs (Figs. S6A and S6B). This suggests that glia in Major may be particularly impactful in shaping the caste-specific gene expression profile observed in the brain of Major workers.

### Juvenile hormone esterase (*cf*Jhe) is specifically expressed by surface glia cells in Majors

In addition to the canonical role of Juvenile hormone (JH3) in regulating insect development, and as an emerging regulator of social insect caste development(*64*), JH3 is also a key hormonal signal that triggers foraging behavior in *C. floridanus* Minor workers (*8*), as well as in other social insects (*26, 27, 65-67*). In *C. floridanus*, JH3 levels are differentially regulated by expression of a JH3 esterase (LOC105248438, hereafter referred to as *cf*Jhe) and a JH3 esterase hydrolase (Jheh2), both of which degrade JH3 and are preferentially expressed in Major worker brains (*8*).

Evaluation of *cf*Jhe expression in our scRNA-seq data showed exclusive expression in SG2 (Figs. 2A–C), one of two subtypes of surface glia detected in our data. Surface glia constitute the insect structure functionally homologous to the capillary endothelial and glial cells of the mammalian brain-blood barrier (BBB) and prevent direct contact between the brain and circulating hemolymph (*32*). In *D. melanogaster,* the BBB is comprised of two types of surface glia, each possessing distinct morphological and physiological features, termed perineurial glia and subperineurial glia (*68*). In *C. floridanus*, we observed multiple genes differentially expressed between the two surface glia subsets (identified using the shared surface glia markers *vkg* and SPARC (*33, 40, 41*) [Fig. 2C]), including Prip, Ndg, TkR99D, and *swim,* biased to SG1; and Tret1, *svp*, and *cf*Jhe biased to SG2 (Figs. 2C and S6C). The SG2-biased genes Tret1 (*69, 70*) and *svp* (*71*) are established markers for perineurial glia, which is the peripheral layer of surface glia in fly (*69, 71*), leading us to conclude that SG2 cells, which exclusively express *cf*Jhe, correspond to perineural glia. Genes expressed by SG1 and SG2 shared similar functional terms indicative of the BBB, including “InR/TOR signal pathway”, which is required for proliferation of surface glia (*72, 73*) (Table S5). We further observed multiple functional terms distinguishing SG1 and SG2 from one another including “transmembrane transport” and “septate junction assembly” in SG1, the latter is a sub-perineurial glia feature in fly (*33*), and “serine family amino acid metabolic process” and “positive regulation of GTPase activity” in SG2 (Fig. S6D, Table S6). Comparison of both surface glia clusters to published fly scRNA-seq data (*40*) also revealed that fly perineurial glia was the most similar cluster with SG2, and SG1 showed the most significant overlap with fly subperineurial glia (Fig. S6E). Interestingly, more than 20 genes annotated as “ROS-like RTK (tyrosine kinase receptor)” and without obvious orthologues in *D. melanogaster*, were highly specific to SG2 (Table S1). These novel predicted membrane-bound cell surface receptors are composed of a diverse extracellular ligand binding domain and an intracellular tyrosine kinase domain (*74*), suggesting that there is active and unique membrane signaling transduction in the ant BBB; active membrane signaling was recently observed to regulate ecdysone signaling in fly fat body cells through the JH-RTK-USP pathway (*75*).

**Fig 2.**
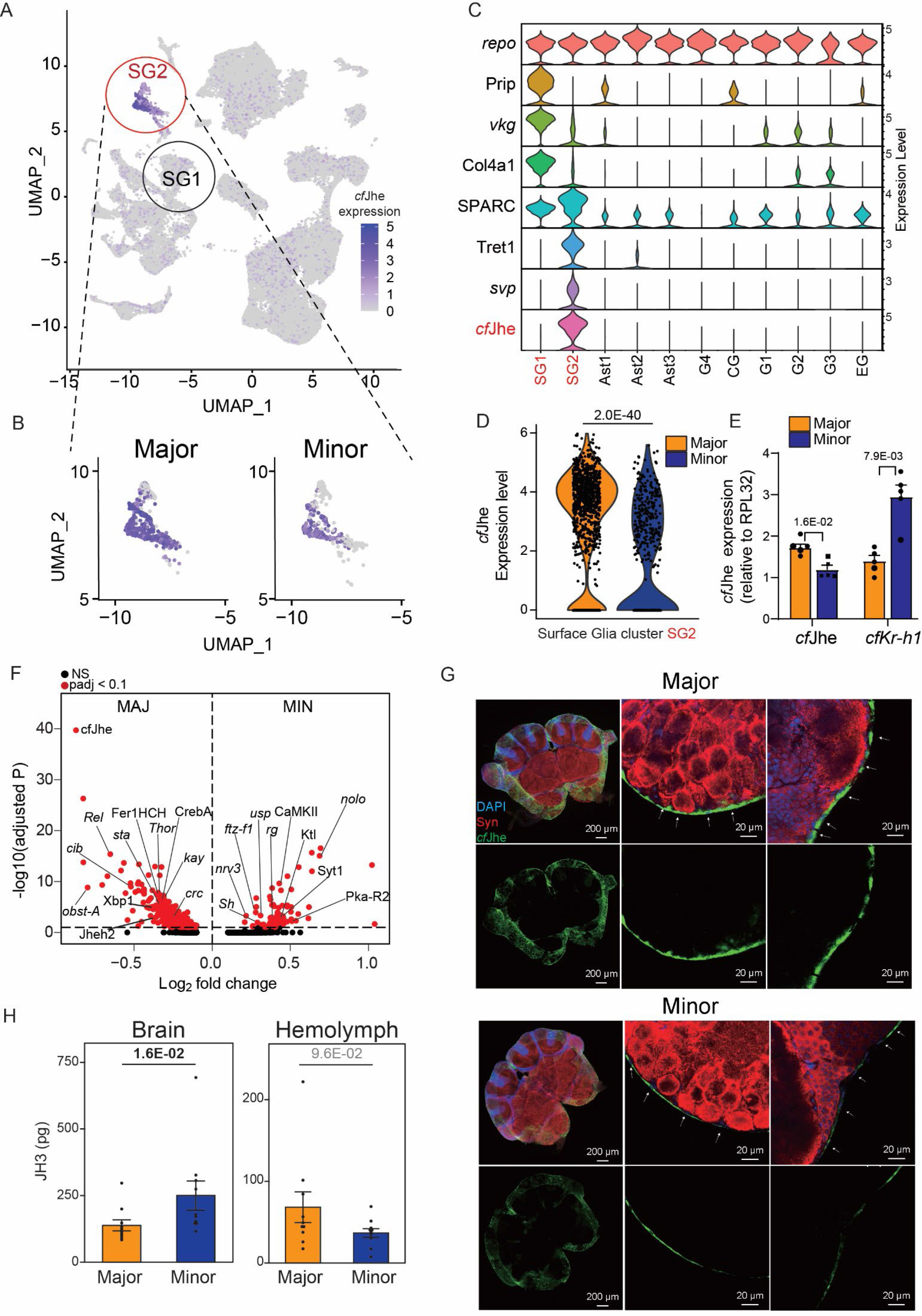
BBB-specific and caste-biased expression of Juvenile hormone esterase, the gatekeeper of JH level in brain. (A) UMAP plot of d0 Major and Minor scRNA-seq data, with per-cell expression of *cf*Jhe plotted. *cf*Jhe demonstrates highly-specific expression in surface glia cluster 2 (SG2). B) UMAP of zoomed *cf*Jhe- containing SG cluster with Major and Minor expression per-cell plotted separately, illustrating caste-specific differences in per-cell expression. C) Violin plot of focal marker genes shared between (top plot) and distinguishing (remaining plots) the two surface glia types here. *cf*Jhe is highlighted and strongly demarcates SG2. D) Violin plot of Major and Minor *cf*Jhe expression level from (C), showing significant Major-bias via differential expression testing between SG2 Major and Minor cells’ expression levels. P-values represent adjusted p-values from Seurat’s FindMarkers function comparing expression from Major SG2 cells to that of Minor SG2 cells. E) RT-qPCR validation of *cf*Jhe and downstream JH3 effector Kr-h1 levels in d0 Major and Minor brain, illustrating negative relationship between *cf*Jhe and expression of Kr-h, a known downstream JH3-responsive gene in other species. n = 5, P-values generated via Mann-Whitney U test (two-sided). F) Volcano plot of DEGs (adjusted p-value < 0.1) between Major and Minor in SG2 (*cf*Jhe+). G) Immunoflouresence of *cf*Jhe contrasting Major and Minor brains (counterstained with Neuronal structure marker Syn and DAPI) First columns show Z-stack of a 100um brain section, second and third columns show the periphery localization of JHE in single slice. H) LC-MS quantification of JH3 levels in the brains (left) and hemolymph (right) of Major and Minor workers, showing significantly elevated brain JH3 in Minor workers but no difference in Hemolymph JH3. n = 10, P-values generated via Mann-Whitney U test (two- sided) after blocking for colony background.

Thus, expression of *cf*Jhe, which has important roles in caste determination in *C. floridanus* (*8*), was highly specific to perineurial glia cells. In addition, we found that *cf*Jhe expression exhibited a notable caste bias, being expressed higher in perineurial glia from Major brain compared to Minor brain (Figs. 2B and 2D), and the population of cfJhe expressing cells in Major SG2 was also more abundant than Minor (80.8% vs 53.5%, Fig. S6F). We validated caste-specificity via RT-qPCR of *cf*Jhe and of *Kruppel homolog 1* (*Kr-h1*), a gene induced by JH3 (*16, 20, 76*), and confirmed Major-biased expression of *cf*Jhe, and a reciprocal pattern of *Kr-h1* expression, further supporting higher JH3 levels in Minors (*8*) (Fig. 2E).

While we focus on *cf*Jhe, we note other key genes specifically expressed in perineurial glia that also displayed a caste-biased pattern of expression, including conserved metabolic enzymes (Major-biased Jheh)(*77*), and hormone-regulated transcription factors (Major-biased *Rel* (*78*), *crc* (*79*) and Minor-biased EcR/*ultraspiracle* (*usp*) (*80*); Figs. 2F and S6G). Along with genes related to hormone regulation, *nolo* (no longer nerve cord), a key component of the neuronal lamina extra cellular matrix that restricts excessive expansion of the CNS (*81*), was among the most Minor-biased genes in perineurial glia (Fig. 2F), potentially functioning to shape CNS development to fit a caste-specific body plan. In neuronal clusters, putative JH3-induced genes Ftz-f1 (*60, 82, 83*) and Hr38 (*84*) showed global Minor-biased expression. Conversely, genes linked to Ecdysone function, including Eip75B (*85*), Hr3 (*86, 87*) and Blimp-1 (*88*), showed Major bias and were restricted in expression to several neuronal clusters (Fig. S6H, Table S4). Increased gene expression of ecdysone response pathways in Major may reflect increased *cf*Jhe and reduced JH3 levels in the developing Major brain, resulting in a shift from JH3 towards ecdysone.

We performed immunofluorescence (IF) against *cf*Jhe protein expression (see Methods for custom antibody production; Fig. S7A) and, consistent with our findings at the single-cell level, *cf*Jhe was specifically expressed within cells at the surface layer of the brain but not within neuronal cells (Syn+) inside; localization and morphology of these cells are consistent with previous surface glia characterization in *D. melanogaster* (*89, 90*)(Figs. 2G and S7B). In addition, a comparison of the *cf*Jhe IF signal from brains of Major and Minor workers showed higher levels in Majors, consistent with our single-cell RNA-seq observations Fig. 2G).

To determine the functional impact of this difference in *cf*Jhe, we used LC-MS (*91*) to measure JH3 levels and determined that brains of Minor workers contain significantly more JH3 compared to Major workers (p=1.6E- 02, Mann-Whitney U test). As a control, JH3 levels in the hemolymph were comparable between castes (p=9.6E- 02; Fig 2H). Hence, caste-specific differences in the level of brain JH3 (independent of the systemic JH3 level) correspond with differences in the level of *cf*Jhe between castes.

### Caste-specific pupal brain maturation is associated with Jhe in BBB

We evaluated whether *cf*Jhe expression is associated with the onset of adulthood just after eclosion, a timeframe that we previously connected as decisive to behavioral division of labor in *C. floridanus* (*8*). We assessed *cf*Jhe expression in surface glia across pupal development (Fig. 3A) using RT-qPCR. *Cf*Jhe expression increased exponentially during the last week of pupal development, starting at pupal day 15, and by day 20 was trending but not yet significantly different (p=5.5E-02, Mann-Whitney U test) between Major and Minor (Fig. 3B), suggesting the mid-to-late pupal stage is critical in behavioral caste determination. Thus, we chose day 18 in pupa (p18) as a critical timepoint in pupal *cf*Jhe expression change and as a comparative reference to adult. To validate this timepoint, we examined *cf*Jhe protein levels in the brain across pupal stages and confirmed that p18 accurately captured increasing expression of *cf*Jhe without differences between castes, which emerge within 2 days following this timepoint (Fig. 3C).

**Fig 3.**
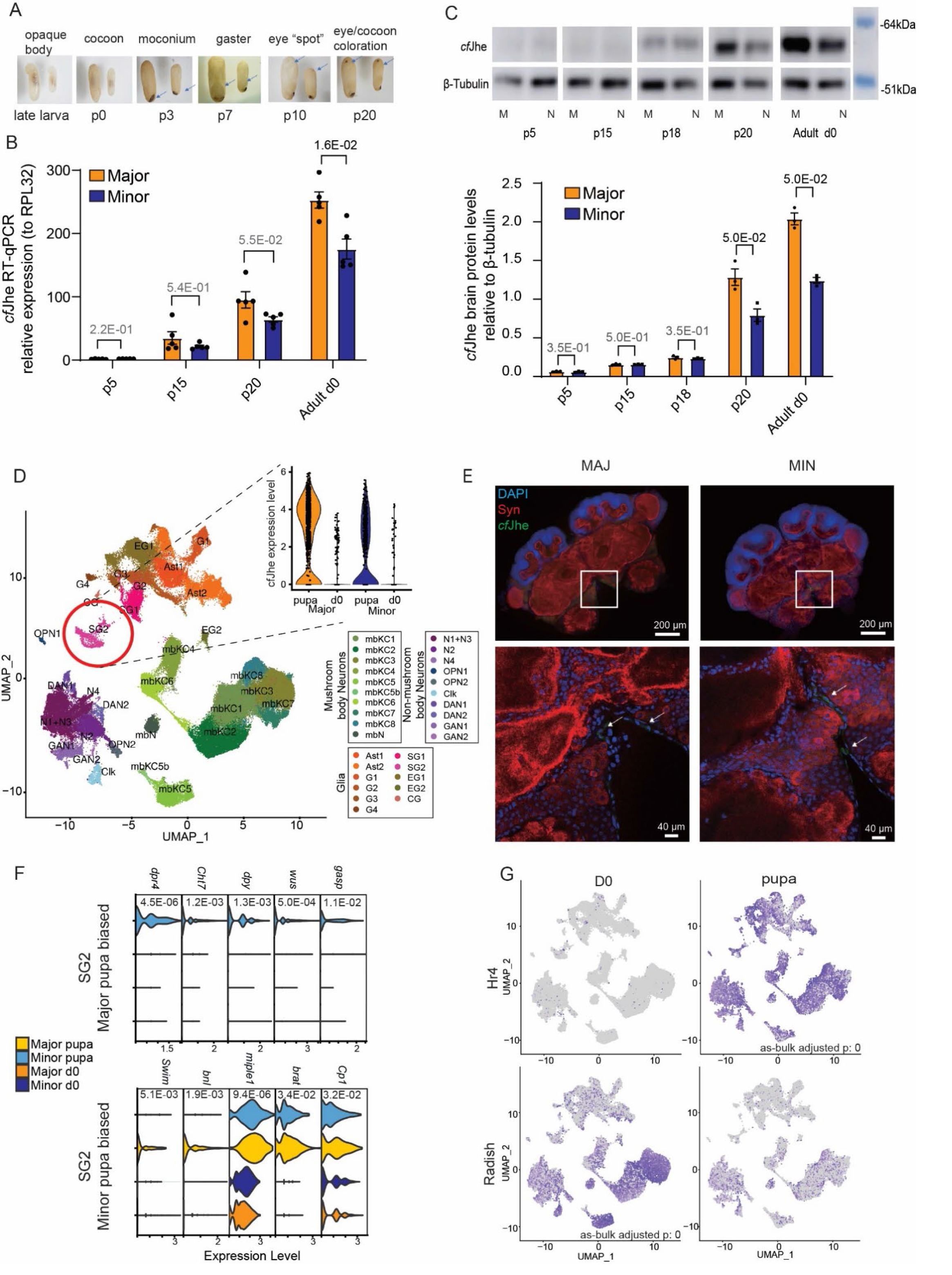
Mid-to-late pupa brain maturation and emergence of *cf*Jhe caste bias in developing *C. floridanus* Major and Minor pupae. A) Morphological progression of Major and Minor pupal development, illustrating major phenotypic benchmarks used to identify key timepoints. B) RT-qPCR (n=5) and C) western blot (n=3) of *cf*Jhe expression levels at key pupal timepoints illustrating p18 as the timepoint where *cf*Jhe is just beginning to exhibit differences between Major and Minor workers. P-values generated via Mann-Whitney U test (two-sided for RT-qPCR and one-sided for WB as a validation). D) UMAP of integrated d0 and pupa (p18) scRNA-seq data, with cell type identities taken from d0 data, illustrating that *cf*Jhe+ cells are significantly under-represented in pupa (violin plot inset). E) Immunofluorescence of *cf*Jhe in Major and Minor pupa illustrating few cells expressing *cf*Jhe at p18. F) Violin plots of pupa-biased | caste-biased TFs in SG2, for (top) Major-biased genes showing proliferation or differentiation-related function, and (bottom) Minor-biased genes showing remodeling-related function. All presented genes are significant (adjusted p-value < 0.0001) via differential expression testing in Seurat (Wilcoxon test). G) Selected stage-specific signaling-related genes exhibiting striking d0 (Radish) or pupa (Hr4) bias.

We performed scRNA-seq on p18 Major and Minor brains using the same protocol as d0 above, and integrated the datasets. Most neuronal and glial cell identities were established at the earlier p18 (Fig. 3D, Table S2 and S8), which is similar with findings in fly brain at equivalent mid-to-late pupal stage (*92*). For both d0 and combined p18+d0 clustering, we assessed representation differences of clusters between castes or developmental stage. There was only one significantly differing cluster between d0 Majors and Minors (SG2; Fig S7A and B) but we identified one cluster showing significant bias to d0 (compared to p18) – mbKC3, a dopaminergic neuronal cluster (Figs S7C-E). In SG2, *cf*Jhe expression was significantly lower in p18 and not yet differentially expressed between castes (Fig. 3D, inset). Consistently, IF staining only detected very limited *cf*Jhe in only a limited number of cells in the p18 brain in both castes (Fig. 3E). Similar to *cf*Jhe, most of the hormone regulation-related TFs that were biased between castes in SG2 at d0 (e.g. Ktl (*93*), Ftz-f1 (*60, 82, 83*), *usp* (*80*) and *Rel* (*78*)) were not significantly differentially expressed in p18 (Fig. S9A). Nonetheless, GO terms enriched among Major or Minor- biased genes in SG2 cells in p18 revealed distinct functions between castes (Fig. S9B, Table S6). Minor SG2 cells were characterized by remodelers involved in extracellular matrix organization, such as multiple chitinases (Cht5, Cht6, Cht7)transmembrane traffickers [e.g. *wurst* (*wus)* (*94*), *dumpy (dpy)* (*95*), *Gasp* (*96*)], and synapse formation [dpr4 (*97*)] (Fig. 3F, Table S9). In contrast, Major SG2 cells at p18 displayed active regulation of both proliferation and differentiation, which is likely establishing the enhanced *cf*Jhe+ cell identity in Majors upon eclosion (Figs. 3F and S9B). Major biased self-renewal and differentiation regulator genes include miple1 (*98, 99*), *brain tumor* (*brat*)(*100*), *branchless* (*bnl*)(*101, 102*), Mi-2 (*103*) and wg/Wnt signaling pathway regulators, such as Swim (*104, 105*), *crooked legs (crol*)(*106*) and Cp1 (*107*). (Fig. 3F, Table S9).

Together these data suggest that prior to caste-biased up-regulation of *cf*Jhe protein, caste-biased gene expression is already established among the *cf*Jhe+ BBB cell types. Hence, comparing p18 to d0, we observed multiple key genes globally regulated in a stage-specific manner, such as Hr4 (*108*) which is expressed only in pupa, and Radish (*109*) which is d0-specific (Fig. 3G, Table S10). Further, several of these genes showed significant differential expression between castes (Fig. S9C, Table S9), showing that differences in caste neurodevelopment are in place prior to emergence of the caste-specific *cf*Jhe+ BBB cells. Nevertheless, the emerging *cf*Jhe+BBB cells likely trigger downstream global gene differences between pupa and adult, by altering the JH3 levels in the brain.

We found a small tail of cells in the Kenyon cell cluster (mbKC5b) in p18 data that were also observed within our d0 data; these cells specifically expressed cell markers characteristic of newborn neurons (e.g. *Nerfin- 2* (*110*) and *Hey* (*111*); Figs. S9D and S7E) in addition to previously identified *Zelda* (*52, 53*), and these genes were significantly higher in Major pupa (Fig. S7F). Together with Major-biased self-renewal markers identified in pupa SG2 (Fig.3F), these data indicate that the Major brain is undergoing more active proliferation or differentiation at p18.

### *cf*Jhe has lost the property of secretion into the hemolymph

The BBB controls entry and exit of solutes from the brain through tight or septate junctions that prevent free diffusion, and through facilitated transport via channels, importers, or transporters (*33*). However, enzymatic degradation of hormones at the BBB has, to our knowledge, not been previously reported. In solitary genera in *Insecta*, such as *Drosophila* (*112*)*, Aedes* (*113*) and *Bombyx* (*114*), Jhe degradation enzymes are largely secreted from fat body cells for circulation within the hemolymph (*77, 115, 116*). Hence, gain of a JH degrading system that is brain-localized and BBB-specific is remarkable, and in the context of eusociality may have a unique role. We compared the Jhe-like gene family in *D. melanogaster* and *C. floridanus* (Fig. 4A) and found strong homology between the catalytic domains of *cf*Jhe and *dm*Jhe (Fig. 4B), in the context of an expanded family of Jhe enzymes in ants (*31*). Interestingly, *cf*Jhe is distinct from *dm*Jhe and from three other *C. floridanus* esterases (Est15, Est18B, Est17) in the same clade: *cf*Jhe lacks a complete N-terminal secretion signal peptide and cleavage site (Fig. 4C, left) and is the only esterase showing brain expression among the four *C. floridanus dm*Jhe homologs (Fig. 4C, right), or among any of the esterases examined.

**Fig 4.**
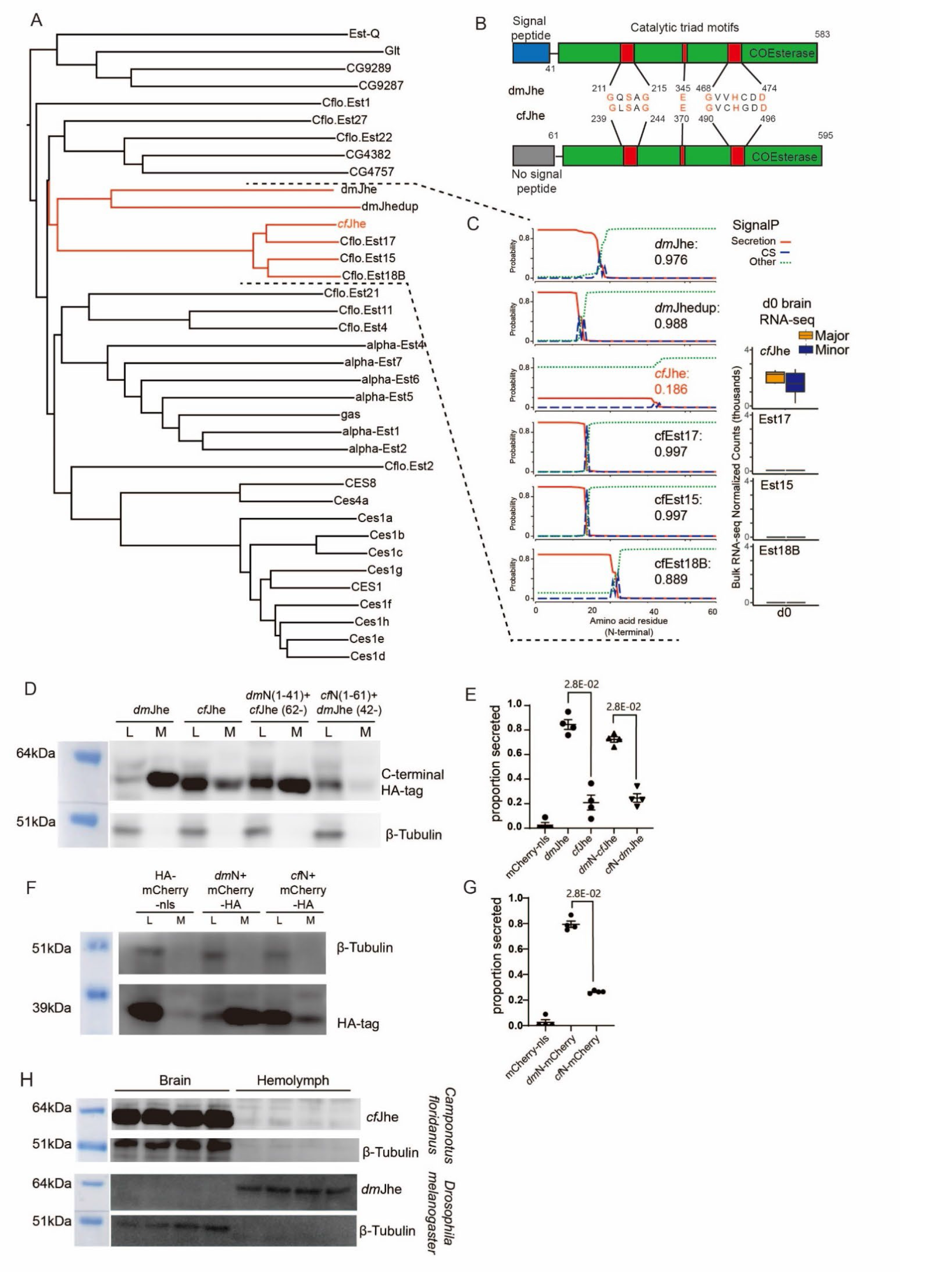
Evolutionary context and loss of secretion in *cf*Jhe. A) Phylogenetic tree of Jhe family in *C. floridanus*, *D. melanogaster* and *M. musculus*. B) Conserved catalytic triad domain comparison between *cf*Jhe and putative parent copy of Jhe from *D. melanogaster* (*dm*Jhe). C) Loss and gain of function of *cf*Jhe. Left: SignalP secretion prediction of *cf*Jhe and closest other esterases in *D. melanogaster* and *C. floridanus* illustrating *cf*Jhe’s unique loss of secretion signal, .Right: brain bulk RNA-seq expression levels for d0 Major and Minor workers illustrating only *cf*Jhe is expressed in the brain. D) Western blot of cell lysate (L) vs media supernatant (M) of *cf*Jhe (cytoplasmic), *dm*Jhe (predominantly secreted), and chimeric *dm*N-*cf*Jhe and *cf*N-*dm*Jhe by switching N-terminus of *dm*Jhe (1-41 a.a.) and *cf*Jhe (1-61 a.a.) upon overexpression in S2 cells, illustrating predominant retention of *cf*Jhe and *cf*N-*dm*Jhe but predominant secretion of *dm*Jhe and *dm*N-*cf*Jhe into culture media. E) Secretion ratio, % of media/(lysate+media), comparison for western blot data from (D) and (F). n = 4, P-values generated via Mann-Whitney U test (two-sided). F) Western blot of cell lysate vs media supernatant of mCherry-nls (nuclear), chimeric *dm*N-mCherry and *cf*N-mCherry by conjugating N-terminus of *dm*Jhe (1-41 a.a.) and *cf*Jhe (1-61 a.a.) to mCherry upon overexpression in S2 cells, illustrating predominant retention of *cf*N-mCherry but predominant secretion of *dm*N-mCherry into culture media. G) Secretion ratio, % of media/(lysate+media), comparison for western blot data from (F). n = 4, P-values generated via Mann-Whitney U test (two-sided). H) Western blot of *cf*Jhe and *dm*Jhe in corresponding brain cell lysate vs hemolymph supernatant. n = 4.

To test whether the different N-terminal sequence affects secretion of *cf*Jhe, we expressed *cf*Jhe-HA and *dm*Jhe-HA in S2 cells in culture, and compared protein levels within cells and secreted into the media (Fig. 4D). Unlike the highly secreted *dm*Jhe (84% secreted), *cf*Jhe was largely retained within cells (21% secreted); 2% of control nuclear-retained HA-tagged mCherry fused to a nuclear localization signal was detected in the media (Fig. 4D, 4E, left lanes). To investigate the sufficiency of the N-terminal sequences of *cf*Jhe and *dm*Jhe for regulating secretion levels, we swapped the non-homologous N-terminal sequences (1-41 a.a. of *dm*Jhe and 1-61 a.a. of *cf*Jhe, Fig. 4B) on the two proteins (*dm*N-*cf*Jhe and *cf*N-*dm*Jhe) and repeated our secretion assay as before. We found that the N terminus from *Drosophila* Jhe was sufficient to promote secretion of cfJhe, whereas fusion of the N terminus from the ant protein inhibited secretion of *Drosophila* Jhe (Fig. 4D, 4E, right lanes). To test sufficiency of the N-termini for secretion or retention, we fused each N-terminal sequence to mCherry, which recapitulated the striking difference in secretion between *cf*Jhe and *dm*Jhe (*dm*N-mCherry: 79%, *cf*N-mCherry: 26%; Fig. 4F. 4G).

To examine brain specificity of *cf*Jhe and secretion status of Jhe in both species *in vivo*, we developed a custom antibody targeting *dm*Jhe and assayed brain and hemolymph Jhe protein levels in both ant and fly. Consistent with our *in vitro* results (Fig. 4D, 4E), *cf*Jhe was highly expressed in brain but not secreted into hemolymph (Fig. 4H). In contrast, *dm*Jhe was secreted and present in fly hemolymph, but not in the brain (Fig. 4H), consistent with previous findings (*40, 117*).

Hence, *cf*Jhe exhibits novel cell retention supporting a role as an intracellular JH3 degrader to regulate JH3 levels in a cell-type or tissue-specific manner; this mechanism is markedly different from Drosophila Jhe, which is secreted and broadly diffused in hemolymph (*77, 112–116*).

### Knockdown of *cf*Jhe and JH3 injections promote a Minor-like transcriptome in Majors

We tested whether *cf*Jhe attenuates JH3 entry into the brain using two approaches: injection of double stranded siRNA targeting *cf*Jhe for knockdown (KD) into the brain of Majors (Fig. 5A; validation Fig. S10A), and injection of excess JH3 or control DMSO into the gaster of Majors (validation Fig. S10B), followed by RNA- seq. For both experiments replicates were taken from at least 5 distinct colony backgrounds, matched between conditions, and this information was incorporated into analyses to control for effects of colony background (see methods). We predicted that either treatment would shift transcription towards a Minor-like transcriptional profile due to a higher level of JH3 crossing brain blood barrier and entering the brain.

**Fig 5.**
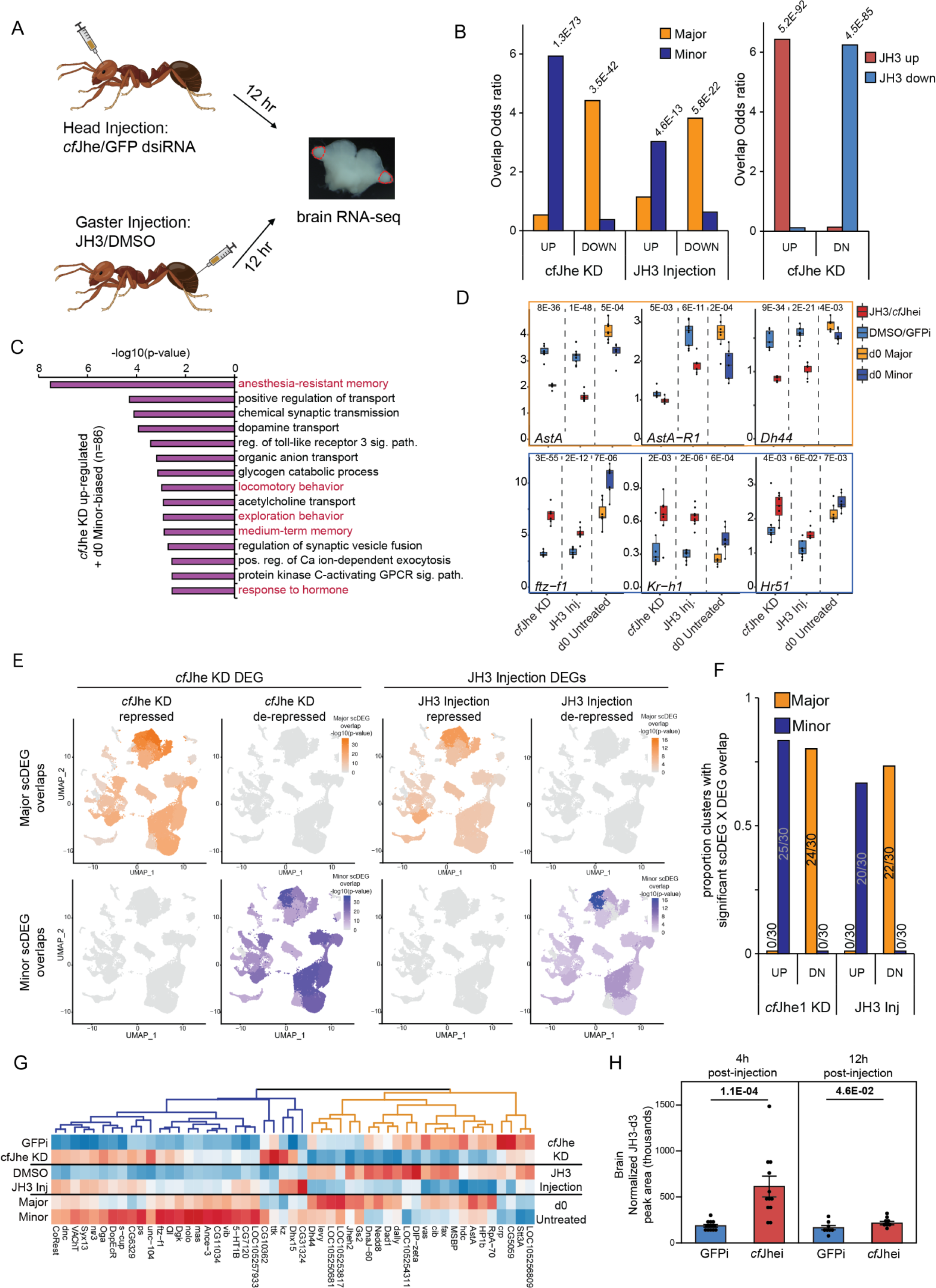
***cf*Jhe KD or JH3 injection both reprogram Major brain transcriptome to a Minor-like state.** A) Model of injection. JH3/DMSO n=7, Jhei/GFPi n=9 B) Odds ratios for overlaps (Fisher’s exact test) comparing bulk RNA-seq DEGs from *cf*Jhe KD (left) and JH3 gaster injections (middle) to d0 Major and Minor-biased DEGs, as well as (right) comparing *cf*Jhe KD DEGs to JH3 gaster injection DEGs. Fisher’s exact test p-values given above bars for all significant overlaps. C) GO terms associated with *cf*Jhe KD up- regulated DEGs also showing d0 Minor-worker bias (left) illustrating strong presence of foraging and neuronal terms in JH3-upregualted genes that are also biased to Minor workers. Significant DEG’s defined as genes with a DESeq2 adjusted p-value <0.1 in both datasets. D) Boxplots of expression (DESeq2 normalized counts) of single-gene expression for bulk RNA-seq from *cf*Jhe KD, JH3 injection, and d0 untreated castes illustrating multiple concordant genes across all three experiments. Plots boxed in blue show significant Minor-biased expression in untreated castes as well as significant bias to JH3 injection and/or *cf*Jhe KD, while those boxed in orange show the opposite (Major-biased expression and repression upon JH3 injection and/or *cf*Jhe KD). P-values represent adjusted p-value from DESeq2. E) UMAP plot visualizing the overlap of *cf*Jhe KD (left) and JH3 injection (right) repressed/derepressed genes with each clusters’ caste-biased scDEGs (determined by comparing Major to Minor cells’ expression for the given cluster via Seurat), showing that *cf*Jhe KD (or JH3 injection) represses Major-biased genes and de-represses Minor-biased genes, but not *vice versa*. For each cluster all genes significantly differing between Major and Minor for that cluster were compared to bulk *cf*Jhe KD (or JH3 injection) DEGs, and –log10(p-value) of overlap (fisher’s exact test) was generated for that cluster. F) Bar graphs showing proportions of clusters with significant overlap of bulk *cf*Jhe KD (left) and JH3 injection (right) DEGs with scDEGs biased to Major or Minor. G) Heatmap of top genes (padj < 0.001 in all three comparisons. n=51, full list presented in Fig S11) showing consistent *cf*Jhe KD significant differential expression, JH3 injection significant differential expression, and d0 Major vs Minor significant differential expression. H) LC-MS quantification of JH3-d3 levels (normalized peak area) of gaster-injected JH3-d3 present in the brain (left) after *cf*Jhe and GFP KD from 4h (left) and 12h (right) following JH3-d3 injection. n=12 for 4h and n = 8 for 12h, P-values from a Mann-Whitney U test (two-sided) after blocking for colony background.

At the transcriptional level, consistent with our hypothesis, both *cf*Jhe KD and JH3 gaster injection led to up-regulation of Minor-biased genes and down-regulation of Major-biased genes (Fig. 5B, left). The two datasets showed similar effects with significant overlap between *cf*Jhe KD and JH3 injection up-regulated genes, and *cf*Jhe KD and JH3 injection down-regulated genes (Fig. 5B, right), and a significant positive correlation between log2 fold-changes (Fig. S10C). Functional analysis of gene classes up-regulated in *cf*Jhe KD and with Minor bias in d0 bulk RNA-seq show enrichment for terms aligning with JH3 in modulating behavior, and, in particular, foraging behavior; top terms included “exploration behavior”, with other terms related to behavior and cognition (Fig. 5C). In addition, “response to hormone” was among the top terms, which includes genes of hormone response in the fly (e.g. Ftz-f1 (*82, 83*), DopEcR (*118*) and *Kr-h1* (*76*), and *usp* (*80*)) (Fig. S10D, left).

Upregulated DEGs in *cf*Jhe KD-induced/Minor-biased dataset included *Kr-h1* (*16, 76*), *nolo* (*81*), *usp* (*80*), and Hr51 (*119, 120*), whereas *cf*Jhe KD-repressed/Major-biased genes included Allatostatin-A and its receptor AstA-R1 (*121*), nAChRalpha1 (*122*), Dh44 (*123*) and HP1b (*124*) (Figs. 5D and S10D, Table S11). These DEGs were previously characterized as regulators of JH synthesis or signaling, neuronal signal transmission, CNS development, or chromatin regulation, and several have been directly associated with caste differentiation (*16, 76, 81, 119-124*). Notably, the ortholog of fly Jheh, another caste-specific JH3-degrading enzyme (*8*), was repressed upon *cf*Jhe KD, suggesting that Jheh regulation is downstream of *cf*Jhe. We identified neurotransmitter transporter/receptors (e.g. DopEcR(*118*)), nACRalpha1 (*122*), GABA-B-R1 (*125*)) in the altered caste-biased DEGs after *cf*Jhe KD (Fig. S10D, bottom blue). Further, CoREST and ttk—previously linked to regulation of worker caste specific behavior via *cf*Jhe repression (*8, 126*)—were upregulated upon both *cf*Jhe KD and JH3 injection (Fig. S9D). This upregulation of CoREST and ttk suggests a positive feedback loop where JH3 induces CoREST and ttk, which in turn bind to the *cf*Jhe promoter for long-term repression during Minor development. Overall, the intersection among the DEGs from the three bulk RNAseq datasets—*cf*Jhe KD, JH3 injection, and d0 untreated castes—reveals key genes driving caste plasticity via *cf*Jhe-mediated JH3-induced pathways.

We compared the bulk RNAseq DEGs from *cf*Jhe KD and JH3 injection to single cell RNAseq caste DEGs (scDEGs). For d0 scDEGs by cluster, there was significant overlap between Minor-biased scDEGs with *cf*Jhe KD/JH3 injection up-regulated genes, whereas Major-biased scDEGs overlapped with *cf*Jhe KD/JH3 injection down-regulated genes (Figs. 5E-F, S10E and S10F; Table S12). For pupa p18 scDEGs, only *cf*Jhe KD/JH3 injection down-regulated genes but not up-regulated genes, showed similar strong caste-biased overlap (Fig. S10G). This suggests that at p18, when *cf*Jhe is first beginning to be differentially regulated between castes (Figs. 3B and C), the initial neuronal transcriptomic response to lowering JH3 levels is to activate normally JH3- repressed genes. Interestingly, the *cf*Jhe+ surface glia cluster (SG2) showed strong overlap between Minor-biased scDEGs and *cf*Jhe KD repressed DEGs, but less so with JH3-repressed DEGs (Fig S10E and F). This possibly suggests that *cf*Jhe expression responds quickly to alterations in hormone levels in an attempt to restore Major worker brain homeostasis, despite persistence of effects downstream of JH3 in other cell types show slower response to altered JH3 levels. Consistent with this, we observed upregulation of *cf*Jhe at 12h following JH3 administration (Fig. S10H); while not reaching significant levels, this suggests that *cf*Jhe is starting to respond to increased JH3 levels in an attempt to restore Major worker brain internal homeostasis.

Overall, 176 genes showed consistent differential expression across the three conditions: untreated d0 castes, JH3-injection, and *cf*Jhe KD (Top 51 in Fig. 5G, all in Fig. S11). Many of these genes were more strongly induced or repressed with JH3 injection compared to *cf*Jhe KD, possibly due to higher levels/pulse of injected JH3 vs incomplete/transient KD of *cf*Jhe.

To test our model, we used a recently developed JH3 assay (*91*) to examine entry of JH3 into the brain and JH3 degradation via *cf*Jhe in the BBB. We performed *cf*Jhe KD in Major brain, and followed this (6h later) by gaster injection of deuterated JH3 (JH3-d3), utilizing multiple colony backgrounds as for RNA-seq data. We performed LC-MS 4h and 12h post injection of JH3-d3, to compare the level of JH3-d3 entering the brain between *cf*Jhe KD Majors and control injected Majors. We found that *cf*Jhe KD increased the amount of JH3-d3 entering the Major brain 4h following JH3-d3 injection (p=1.1E-04, Mann-Whitney U test; Fig. 5H left), demonstrating that *cf*Jhe reduces JH3 entry into the brain and that KD of *cf*Jhe is sufficient to increase brain JH3. At 12h following JH3-d3 injection, *cf*Jhe KD still showed increased brain JH3-d3 but the magnitude of difference was considerably reduced by this time (p=4.6E-02, Mann-Whitney U test; Fig 5H, right). We noted that the JH3-d3 level in hemolymph of the same individual ants did not show significant difference in either timepoint (p=4.5E- 01 and 1.1E-01, Mann-Whitney U test; Fig. S10I), supporting the specificity of cfJhe KD, and importance of cfJhe for modulating JH3 levels in brain specifically.

### *Cf*Jhe recapitulates Major-like reduced foraging behavior in fly, and knockdown Minor-like enhanced foraging in ants

Given that JH3 stimulates foraging in social insects (*26, 27, 65-67*), we investigated whether manipulating its level via expression of *cf*Jhe would be sufficient to influence foraging behavior in *D. melanogaster*, a solitary species which lacks the social complexity of *C. floridanus*. We expressed *cf*Jhe in all glial cells in *D. melanogaster* using *repo*-Gal4, to evaluate the impact of pan-glial degradation of JH3 on fly behavior (*127*) and then evaluated the sufficiency of surface glia specific expression of *cf*Jhe as a key gatekeeper regulating brain JH activity by using well-established Gal4 drivers including 9-137-Gal4 (Fig. 6A) (*73, 89, 128, 129*). Foraging activity was assayed in adults, by recording exploratory locomotion and food interest within a foraging arena (Figs. 6A and 6B; methods; (*130–132*). Gal4 drivers and UAS-*cf*Jhe lines were crossed to identical wildtype flies as control groups.

**Fig 6.**
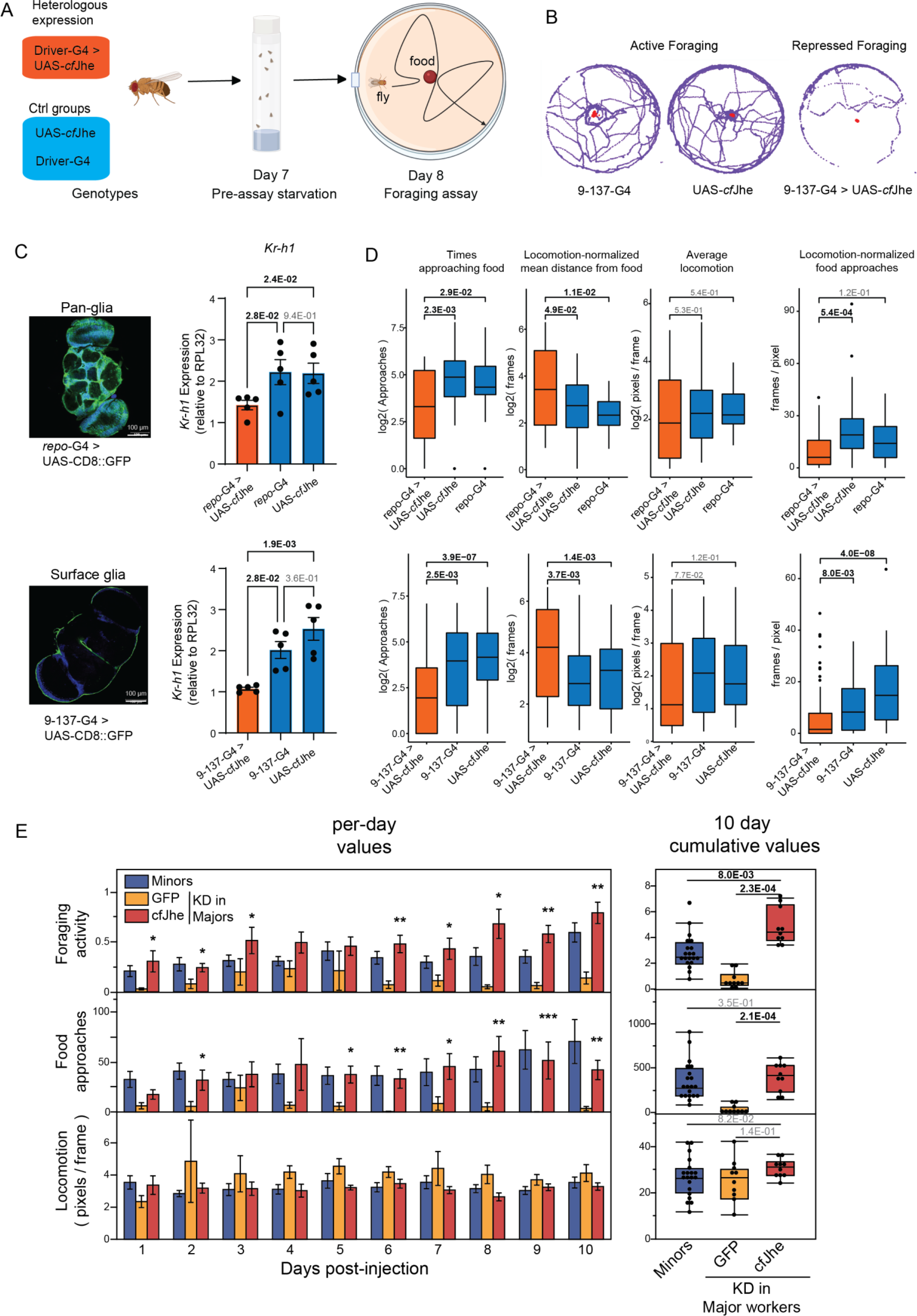
**Trans-organismal foraging behavior alternation by gain and loss of function of *cf*Jhe**. A) Model of *cf*Jhe overexpression experiment (left) and foraging arena (right). B) Representative 10-min foraging trajectory as analyzed by DeepLabCut. C) Pan glia (upper) and surface glia (lower) over-expression of *cf*Jhe illustrating (left) specificity of expression of each driver, (right) RT-qPCR of Kr-h1 illustrating impact of *cf*Jhe on Kr-h1 levels when *cf*Jhe is overexpressed in given glial subtypes. n =5, P-values generated via Kruskal-Wallis tests followed by Dunn’s test (two-sided). D) *cf*Jhe overexpression is sufficient to decrease *D. melanogaster* foraging when expressed exclusively in surface glia. Row order as in panel C, illustrating *cf*Jhe overexpression in pan-glia (top row) and surface glia (bottom row) both increase fly number of times approaching food (left), mean distance from food (left center) and non-significantly impacts locomotion (right center), as evidenced by average locomotion-normalized number of times approaching food (right). n >20, P-values generated via two-sided Kruskal-Wallis tests followed by Dunn’s test. E) RNAi against *cf*Jhe in *C. floridanus* Major workers leads to a marked increase in foraging across 10 days of video tracking, reaching levels not statistically different from (or higher than) those seen in Minor workers from the same assay (right), but does not dramatically impact locomotion (bottom). Majors n=10 per condition, Minors n=20, P-values generated via a two-sided Kruskal-Wallis tests followed by Dunn’s test after blocking for colony background. Asterisks (left plot) represent results of a Mann-Whitney U test comparing a given day’s *cf*Jhe KD values to that of GFP KD control after blocking for colony background.

**Figure 7.**
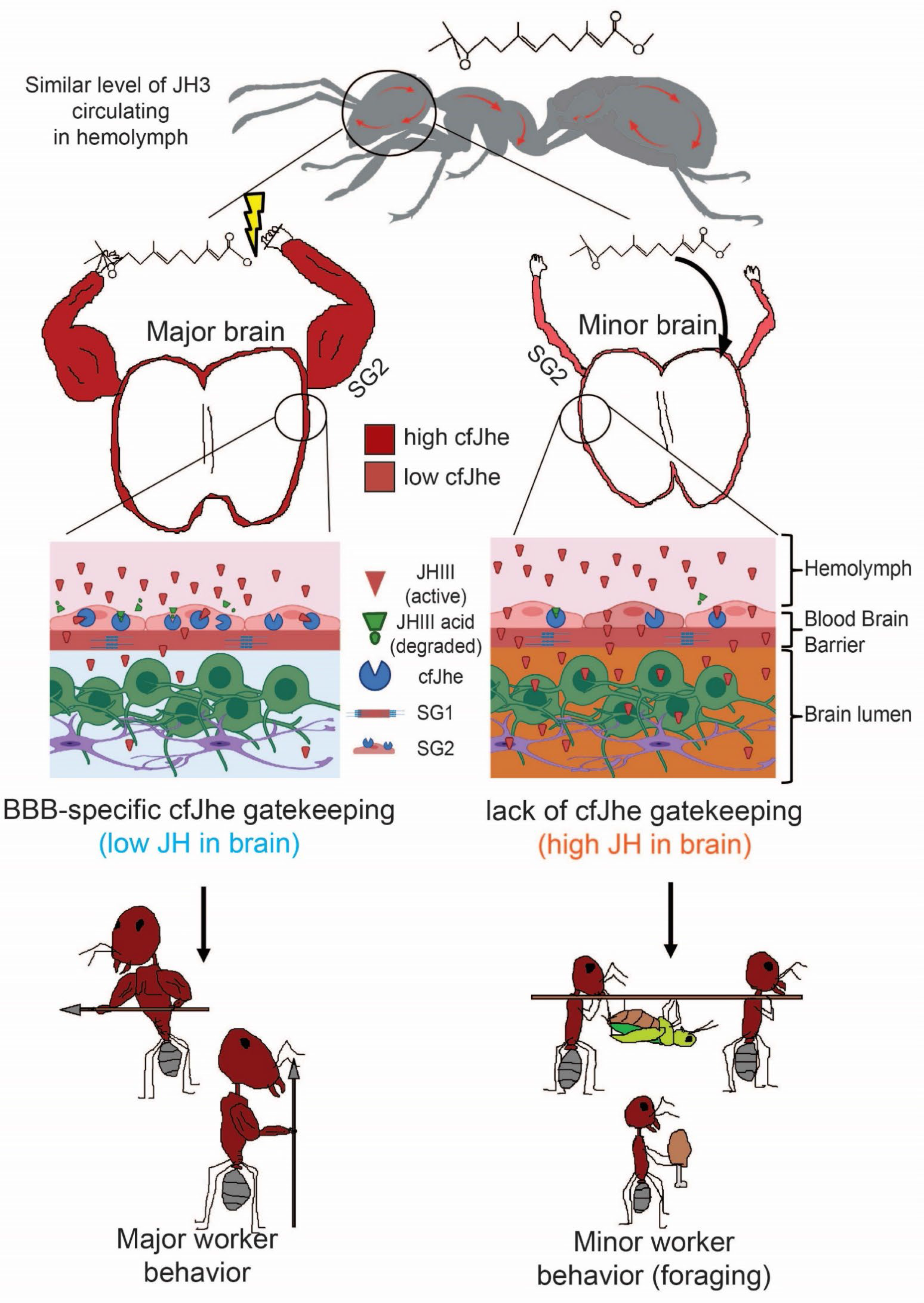
Model of *cf*Jhe-mediated BBB hormonal gatekeeping. JH3 levels circulating in hemolymph are subject to differential filtering in perineural glia between Major (left) and Minor (right) workers, where Major workers expression much higher levels of *cf*Jhe at the blood brain barrier, resulting in decreased JH3 entry into the brain, effecting caste-specific reduction in Major foraging.

We first evaluated *cf*Jhe heterologous expression efficiency and altered JH3 activity by measuring *cf*Jhe and *Kr-h1* mRNA levels (downstream of JH3) in the brain of corresponding genotypes (Figs. 6C and S12A). We also validated that protein levels of *cf*Jhe were increased (Fig. S12B). Pan glial heterologous expression of *cf*Jhe in *D. melanogaster* was sufficient to downregulate *Kr-h1* in the brain, consistent with a decreased level of JH3 in the brain (Fig. 6C upper). Importantly, similar to *C. floridanus* (Fig. 2E), heterologous *cf*Jhe expression in *D. melanogaster* surface glia (Fig. 6C, upper) was sufficient to downregulate *Kr-h1* to similar levels as seen with pan-glial expression (Fig. 6C lower).

To test foraging behavior as an effect of *cf*Jhe expression in surface glia, we recorded individual fly locomotion trajectory for quantitative analysis (Fig. 6B). Overall, adapting from similar fly foraging assays (*130–132*), we divided the foraging pattern into two quantifiable behavioral outputs: locomotion and interest in food, which are behaviors that are either directly affected by metabolism of JH3 (*133–135*), or correlated with signaling of hormones interacting with JH3, such as ecdysone (*136, 137*) and insulin (*135, 138, 139*). Consistent with our prediction, heterologous expression of *cf*Jhe in surface glia and in pan-glia resulted in significant decrease in food interest (fewer times approaching food and larger normalized distance from food); we also used total locomotion as an overall proxy for foraging activity and found similar effects (Fig. 6D).

To evaluate potential bias of specific drivers or genetic background, we repeated the foraging assay using another widely used surface glia driver, NP6293-Gal4 (*89, 128, 129*), to heterologously express *cf*Jhe or crossing driver and UAS-*cf*Jhe to a secondary wildtype, Oregon R, as control groups. Both additional methods of testing consistently exhibited repressed foraging behavior (Fig. S12C).

Since worker ants are females, we further analyzed female behaviors of genotypes in Fig. 6D. Consistent with results in male flies, we observed decreased foraging-like behavior in female exogenous expression lines (Fig. S12D). Furthermore, to evaluate whether there was a potential global behavior defect in exogenous expression lines which affected foraging outcomes, we assayed fly baseline behavior at day 4 and day 7 using a climbing assay (*140–142*), finding no major impact of *cf*Jhe expression on climbing behavior (Fig. S12E). In addition, normalization of number of times visiting food by locomotion did not impact our overall results (right column of Fig 6D, S12C and S12D), indicating that reduced foraging was not due to gross behavioral or locomotor defects.

Importantly, we evaluated the reciprocal effect in ants, testing whether *cf*Jhe reduction was sufficient to increase foraging activity of Major workers. We injected dsiRNA targeting *cf*Jhe or GFP (control, as used in Fig 5 results) and assayed foraging activity over 10 days following *cf*Jhe KD, using automated tracking of ant foraging. For each behavioral replicate we utilized distinct colony backgrounds matched between conditions, and this was accounted for in statistical testing (see methods). Strikingly, *cf*Jhe KD led to increased foraging behavior across 10 days, approaching or exceeding foraging levels exhibited by Minor workers in the same assay (Fig. 6E; Fig. S12F cumulative values). To rule out potential systemic effects of *cf*Jhe KD, we measured *cf*Jhe and Kr-h1 levels in fat body, a widely distributed and metabolically active cell type. Neither *cf*Jhe or Kr-h1 differed between KD and control groups in fat body (Fig. S12G), indicating that the effects of *cf*Jhe KD are due to the resulting changes in JH3 levels in the brain.

Taken together these results suggest that *cf*Jhe has adopted a novel function as a non-secreted version of Jhe, localized specifically within surface glia in *C. floridanus,* and that expression of *cf*Jhe in *D. melanogaster* surface glia and knockdown of *cf*Jhe in *C. floridanus* Major workers is sufficient to respectively reduce or increase foraging activity — and in the KD in Major, foraging approaches levels detected in Minor workers.

## Discussion

In this study, we uncovered a remarkably simple mechanism driving ant caste-specific behavior. Using single cell brain transcriptomics, molecular structure/function dissection, and cross-species functional studies, we addressed the basis of profound asymmetries in foraging behavior between Major and Minor workers in the ant*floridanus* and their development during late pupal stages. One key finding is that *cf*Jhe, which degrades Juvenile Hormone (JH3) and is critical to caste-specific behavioral differentiation (*8, 116*), shows restriction in expression to one cell type in the brain—the perineurial glia—which forms a structure akin to the vertebrate blood-brain barrier (BBB) (Fig. 2). Further, we discovered that in the Major brain each perineural glial cell expresses more *cf*Jhe than their counterparts in the Minor brain (Figs. 2B and 2E). Recent work has contributed to an emerging view that the BBB is an underappreciated regulatory component of brain development and homeostasis: in the fly, active transport of steroids and a circadian rhythm of BBB permeability regulate larval behavior and adult sleep (*89, 128, 129, 143*), and in mouse, xenobiotic efflux and circadian clocks are highly dependent on the function of the BBB (*143, 144*). Here we show that a caste difference in the hormonal degradation taking place within the BBB via *cf*Jhe is integral to the division of labor underpinning the social structure of ant colonies.

*cf*Jhe is a member of a family of esterases that has expanded in ants (*31*), and our second key finding is that, relative to the ancestral form shared with fly, the Jhe paralogue expressed in the perineural glia has lost its secreted nature, and instead is intracellular in cells of the ant BBB (Fig. 4). Thirdly, we demonstrate that either genetic reduction of *cf*Jhe expression in Major brain, or elevation of systemic levels of the JH3 hormone, lead to a dramatic shift to a Minor-like transcriptome and foraging behavior (Fig. 5-6); these manipulations thus reveal the central importance and sufficiency of *cf*Jhe to differentiate the brain of Major and Minor workers. Finally, despite >300my of divergence, we found that expression of *cf*Jhe in perineurial glial cells in the *Drosophila* brain leads to reduced foraging, consistent with absence of foraging of the Major worker in *C. floridanus* and high *cf*Jhe expression in glial cells (Fig. 6). Taken together, our findings establish that one of several homologs of fly Jhe has adopted a unique role in the brain of *C. floridanus*, to function as a hormonal gatekeeper, restricting entry of JH3 into the developing brain to tune hormone levels, leading to precise caste determination. This unusual *cf*Jhe expression strategy leading to caste specification simultaneously avoids altering systemic levels of a physiologically important hormone.

### Conserved and unique neuronal and glia cell types in *C. floridanus* brain

Here we carry out scRNAseq of *C. floridanus* brain, and identify conserved groups of neuronal and glial cell types compared to other insects, including *D. melanogaster*. Similar to previous observations in social insects (*145–148*), the mushroom body of *C. floridanus* comprises ∼39.8% of brain cells (60.6% of neurons) and comprises subtypes of intrinsic Kenyon cells, whereas glial cells comprise 29.5%. The relative abundance of mushroom body neurons and glial cells is markedly higher than in *D. melanogaster* (mushroom body: 5-10% of neurons, glia: 2-10% of brain cells) (*40, 54, 149*), which is consistent with the observation in *H. saltator* (*41*), suggesting a potential global expansion of mushroom body and glial populations in Hymenoptera.

In neuronal populations, we discovered mbKC6, a subset of mushroom body Kenyon cells (Fig. 1E) notable for expressing neuronal genes associated with newborn neurons. This mbKC6 cluster suggests that neurodevelopment is still actively occurring from late pupa to d0 in *C. floridanus*, as *Kr*, *zelda*, *nerfin-2* and Hey (Fig. S6E) are implicated in neuroblast proliferation or neuronal differentiation (*51, 52, 110, 111*). In glial populations in both ant species and in *D. melanogaster* (*39–41*), we observed two surface glia clusters (*vkg*+, SPARC+) mapping to SG1 and SG2, respectively. However, while there was conservation of core marker expression in SG cell types between the two species, we document unique hormonal metabolic activity in SG2 for *C. floridanus*.

### Novel caste-biased hormonal gatekeeping function of *C. floridanus* blood-brain barrier

Our key observation in *C. floridanus* is that surface glia, the glial populations corresponding to the insect blood-brain barrier, show unique Major-biased features. In particular, one of the SG sub-clusters shows gene expression indicative of hormone degradation (Fig. 2). Strikingly, we found that one paralog of Juvenile hormone esterase (hereafter *cf*Jhe), a degrading enzyme of JH3, was expressed only in one BBB cell cluster, and was highly biased in expression to Major workers (Figs. 2C-E and 2G). Our findings thus illuminate the cell type responsible for differential *cf*Jhe expression and reveal the basis for disparate JH3 levels across castes (*8*).

We reveal that caste-differential expression of *cf*Jhe favoring the Major BBB occurs between 15 to 20 days of pupal development (Figs. 3A and 3B), as determined by scRNA-seq and immunofluorescence. The proportion of *cf*Jhe+ SG2 cells was significantly lower at p18 compared to later d0, raising the question of the origin of *cf*Jhe+ BBB cells, specifically, whether *cf*Jhe+ SG2 differentiate from other glial cells or self-proliferate during the late pupa stage. Deeper analysis of the *cf*Jhe+ SG in p18 revealed strong Major-biased expression of multiple regulatory genes of neuroblast self-renewal and differentiation, including *aurora A* (*aurA*), *brat*, *escargot* (*esg*), and *mushroom body defect* (*mud*) (*100, 150–152*). Thus, at p18, *cf*Jhe+ SG2 are undergoing caste-specific development, with expansion in Majors prior to, or concomitant with upregulation of *cf*Jhe. CoREST has been observed to regulate caste-biased gene expression, including regulation of *cf*Jhe (*8*), consistent with a general model of glia-based caste differentiation, as CoREST has been identified as a major regulator of neuronal and glial differentiation (*153–156*). Our results indicate that rather than mainly undergoing cell enlargement as postulated in *D. melanogaster* (*157*), in ants, SG proliferate over pupal development. Additional developmental analyses and live imaging will fully illuminate this process. Taken together, our results suggest that, after the primary caste differentiation “decision point” in late larval and early pupal development, a second neurodevelopmental component of worker caste differentiation is accomplished in late pupal stage. To our knowledge this is the first observation of such a phenomenon in insects, however, because this is the first reported scRNA-seq from late pupal or newly-eclosed hymenoptera, additional investigation will be needed to validate this finding and hypothesis.

### Novel non-secreted *cf*Jhe, a “gain of function” in social insects over Jhe gene family evolution

We observed two main features of *cf*Jhe that distinguish it from “conventional” Jhe: high specific expression within the perineurial glia, and a striking loss of a secretion signal peptide. These molecular changes are demonstrated by lack of secretion of *cf*Jhe when exogenously expressed in in *D. melanogaster* S2 cell culture (Figs. 4D and 4E), and sufficiency of the *cf*Jhe N-terminus to cause cellular retention of *dm*Jhe (Fig 4D and 4E). Relative to *cf*Jhe, both *dm*Jhe and all other closely-related esterases in *C. floridanus* lack BBB expression and all include a strong signal peptide of secretion; these findings support a model that *cf*Jhe is uniquely expressed by, and retained within the BBB that envelops the brain, thus serving as gatekeeper to restrict JH3 levels entering the brain. We utilized LC-MS to demonstrate higher levels of JH3 in Minor worker brain (Fig. 2H), and, importantly, we show a strong increase in JH3 level in Major worker brain following *cf*Jhe KD (Fig. 5B).

We showed the regulatory consequence of JH3 gatekeeping and caste-determining function via reducing *cf*Jhe in the brain or by injecting JH3 in hemolymph. Both manipulations in Majors resulted in a switch to a Minor-like transcriptome (Figs. 5 and S10). Globally, we observed a strong concordance between DEGs resulting from *cf*Jhe knockdown/JH3 injection in Major workers with Minor-biased DEGs from bulk and single cell RNA-seq profiles (Fig. 5E, 5F). Further, the DEGs provide strong support of a direct link to foraging: the top gene ontology function enriched among genes upregulated by *cf*Jhe KD, that were also biased to Minor workers, was “exploration behavior” (Fig. 5D). Multiple genes we or others (*8, 16*) have linked to regulation of caste, in particular of foraging castes, were up-regulated with *cf*Jhe KD, including *Kr-h1*, *nolo*, Ftz-f1, CoREST and ttk (Fig. S10D). The striking consistency between KD of *cf*Jhe and foraging caste gene expression strongly supports the model that acquisition of unique BBB-localized JH3 gatekeeping underlies the core transcriptional network that promotes foraging in Minor workers. Given the centrality of JH3 to our model, it is likely that other factors such as the JH3 receptor (Met/gce) are involved in this system, however, while expressed in the brain, we did not detect caste- or cell-type bias in Met expression (Fig S10J).

As further crucial evidence*, in vivo* heterologous expression of *cf*Jhe in fly resulted in marked decrease of foraging-related behavior (Fig. 6). Importantly, the efficacy of *cf*Jhe in decreasing foraging was directly related to the glia cell heterologous expression, as surface glial and pan-glial heterologous expression exhibited a reduced foraging behavioral phenotype in both male (Fig. 6D) and female flies (Fig S10D). Though it remains unknown how altered JH levels in the brain regulate foraging status, recent studies of JH signaling in the adult fly brain revealed a role in maturation of mushroom body α’/β’ lobe neurons associated with learning behavior (*158*) and of dopaminergic neurons in a motivational neuronal circuit controlling courtship behaviors in adults (*159*). In addition, we identified these mushroom body Kenyon neurons (*trio*+, *cac*+) and dopaminergic neurons (*ple*+) from motivational circuits in the fly (*158, 159*) within our ant scRNA dataset utilizing the same genetic markers (Fig. 1B, Table S1). Importantly, these cell types were among the responsive neuronal clusters in our *cf*Jhe KD (Fig. S10E) or JH3 injection (Fig. S10F), suggesting conservation in their JH3 responsiveness. These results reveal that the effects of *cf*Jhe result from brain-specific expression, loss of secretion/cytoplasmic-retention, and perineurial glia-localized expression. These features alter downstream JH neuronal signaling within the brain, which is potentially a shared mechanism between species to regulate a spectrum of behaviors, as suggested by our results in fly (Fig 6D).

Furthermore, knockdown of *cf*Jhe in *C. floridanus* Major workers resulted in a striking increase in foraging activity across 10 days following injection, approaching or exceeding levels seen in untreated Minor workers, which typically forage (Fig 6E, S12G). Notably, the changes in Major worker foraging following *cf*Jhe KD were similar to that seen following TSA-injection in our prior work (*8*). This suggests that *cf*Jhe is a central element of Major worker behavioral reprogramming and of natural behavioral programming in the Minor worker, and in both cases this is accomplished through the BBB-specific differential expression of a single, hormone-degrading gene.

### Conclusions and outlook

This study reveals that *cf*Jhe performs a caste-specific function to regulate the amount of JH3 entering the brain. Because JH3 is a systemically-important hormone, alteration of global levels of this hormone would be expected to be problematic during late development or early adulthood. However, our findings indicate that an important component of ant worker caste differentiation involves specific modulation of the levels of this hormone in the brain. *C. floridanus* has achieved an elegant solution by lowering JH3 entering the brain via selective degradation in surface glia. Recent reports have observed BBB-specific hormonal import in fly (*129*), which, together with our results, indicate that the BBB is crucial to brain-specific hormonal regulation. Our findings raise an intriguing possibility that a functionally homologous system exists in mammals—we found that in mouse, several genes with putative hormone-degrading or -altering functions are most highly expressed in brain endothelium relative to other endothelial cells (Fig. S10K) (*160*)). Ark1c14 and H2-Ke6 (aka 17β-HSD), which both degrade testosterone (*161–163*), displayed higher expression in brain vascular endothelium than in endothelial cells from four other tissues, and compared to whole brain (Fig. S10K, left). These data suggest that BBB-based hormonal degradation may be a broader mechanism for regulating brain hormone levels.

Finally, in addition to the focal brain-specific JH3 regulation in this study, JH3 signaling is also the pacemaker of many other cellular activities not directly related to the brain, such as of development and reproduction. Thus, dynamic regulation of JH3 in hemolymph likely occurs over the ant life cycle. The *cf*Jhe+ surface glia must thus adapt to this dynamic change and continue to sustain the caste barrier during fluctuations in global JH3 levels. Future work will reveal the breadth of this mechanism in controlling social insect caste, and of behavior in general across the animal kingdom.

## Methods

### Ant husbandry and sample selection

Mature, queen-right colonies of *C. floridanus* were used in this study, collected as foundresses from the Florida Keys, USA, in 2007, 2011, and 2015. Colonies were maintained in a sealed environmental growth chamber at constant temperature (25°C) and humidity (50%) under a 12:12 light:dark cycle. Colonies were fed twice weekly with excess supplies of water, 20% sugar water (sucrose cane sugar), and Bhatkar-Whitcomb diet (*164*).

Worker caste was determined as in (*7*). Adult workers within one day of eclosion (d0) were identified by their location among brood in the nest (rested close to pupa incubation zone), their general behavior (less offensive), light cuticle coloration (compared to a reference panel of ants aged from pupation through 30 days) and the lack of a pair of “eyebrow”-like bulges near eyes, which usually appears at day 3-4. Pupa workers were isolated at pre-pupae stage (∼1-3 days before pupation, assessed based on size and opaque body color) together with 3 Major and 10 Minor adults from the same colony to simulate a natural colony environment. Only individuals entering pupal stage with complete cocoon the next day were recorded as day-0 pupae (p0) for further age-matched experiments and the rest were returned back to original colonies.

For all assays involving ants, Majors and Minor workers were taken from at least three distinct genetic backgrounds – bar RT-qPCR validation figures S10A and S10B, for which only two genetic backgrounds were used. Colony backgrounds and ratio of each colony between experimental and control groups are the same, and colony-of-origin information was incorporated into analyses when possible. Full details of colony-of-origin and sample numbers taken are given in Table S16.

### cfJhe and dmJhe Cloning

Full length cDNA sequence of *cf*Jhe and *dm*Jhe were reverse transcribed (SuperScript III, Invitrogen) and amplified (Q5 high-fidelity 2x master mix, NEB) using RNA extracted from adult *C. floridanus* brain and *D. melanogaster* whole body tissue (Qiagen RNeasy Mini Kit).

### Generation of UAS-cfJhe Mutant

UAS-*cf*Jhe knock-in fly strain was generated using BestGene Transgenic services based on PhiC31 integrase-mediated transgenesis system by injecting targeting plasmid (pBID-5xUAS-*cf*Jhe), into embryos of a y^1^w^67c23^, P(CaryP)attP40 strain followed by PCR screening and balancing to get heterozygous mutant. Individuals not carrying *yellow* mutant allele were picked to maintain stable heterozygous or generate homozygous and outcross lines.

### Generation of cfJhe/dmJhe antibody

Rabbit anti-*cf*Jhe polyclonal custom antibody was generated as described previously (*8*). Briefly, a GST-tagged antigen was produced corresponding to a.a. 275-461 of *cf*Jhe and a.a. 370-579 of *dm*Jhe, then provided for rabbit injection by Cocalico biologics. An MBP-tagged version of the same antigen was then used for affinity purification of polyclonal antibodies from resulting antisera. The resulting antibody was validated using western blot of the antigen and whole body lysate from the respective species the antigen was derived from.

### Fly Crossing and Rearing

All overexpression crosses (Gal4 driver > UAS-*cf*Jhe/UAS-CD8::GFP) as well as control crosses (Gal4- driver/UAS-*cf*Jhe > WT Iso31/Oregon R) for RT-qPCR, behavioral assay and fluorescent imaging were reared at 25°C and 50% humidity on a 12-hour light/dark cycle using standard Bloomington Drosophila Medium (Nutri-Fly). Day0 males and females were separated and reared in new vials (∼30 flies per vial). 3 old males were put together with females in the same vial to have females mated. Detailed genotype was in Table S13.

### Fly Climbing Assay

Fly climbing assay was performed mainly following previous method(*140*).20 male or female flies of each genotype were transferred to a new vial at least one day before test (no more CO2 anesthesia). At test day, flies were transferred to the test chamber (28.5 * 200mm transparent cylinder with a line drawn at 5cm from bottom, which was assembled using two drosophila vials). 3 taps were given to knock all flies in the chamber to the bottom. After 10s of climbing, % of flies not passing the drawn line was recorded. This assay was repeated for 3 times with 1min interval.

### Fly Foraging Assay

At day7 after eclosion, flies were transferred to a new vial with only wet cotton foam for 24-hour starvation, as a pre-assay treatment maintaining basal food-motivated foraging behavior as reported in previous adult foraging assay (*130–132*). All foraging assays were performed at day8 2-7pm and all data reported were from male flies.

The fly foraging assay arena and procedure were adapted from two previous adult foraging assay designs with minor modification for large-scale experiments (*131, 132*). In brief, a 60mm diameter single-fly foraging arena dish with a food drop holder in the center and an entrance at side wall was 3D printed using polylactic acid material and arrayed as a 2x3 6-mer plate. A universal tunnel at each long side connecting 3 entrances and a loader sliding in the tunnel also printed same as previous design to hold and gently push loaded flies to the entrance. A transparent acrylic board was placed on top to prevent escape and allow video tracking. The detailed design of foraging arena (.stl file) is in [SUP_FILE_Arena].

Foraging assay was performed in a new fly incubator with the same rearing conditions. 24 flies of each genotype were assayed at the same batch using 4 sets of 6-mer arena together. Before each batch of foraging assays, all parts were wiped with 70% ethanol, sterile water and dry tissue in turn, then air-dried for 10 mins. A drop (∼15- 20ul) of fresh fly food (50:50:1 mixture of Fleischmann’s dry yeast, warm sterile water and McCormick Culinary red food color) was pipetted at the center of each arena. Flies of the same genotype were carefully aspirated and loaded into the tunnel but restricted to enter the arena before pushing the loader to the correct position. Flies were first allowed to acclimate for 30 seconds after entering the arena, then a remote control camera (GoPro Hero Black 8/DJI Osmo Action) was used to record fly locomotion from above for 10mins.

### Fly Foraging Data Analysis

24 foraging arenas from one batch of experiment were first cropped as individual videos using VideoProc. Individual fly locomotion and interaction with food were tracked using DeepLabCut (version 2.2.0.3, single-animal mode) (*165*). In brief, demo fly foraging videos (416 x 416 pixels) with “fly” and “food dot” manually labeled in 100 extracted representative frames were provided as a training dataset. We trained a ResNet50-based neural network with default parameters in “pose_config.yaml” for 1030000 of training iterations to identify objects similar with “fly” and “food dot” in new videos. Combined evaluation result output: Shuffle number =1, Train error (px) = 1.07, Test error (px) = 1.87, p-cutoff used = 0.6. The coordinate of fly and food dot at each frame was recorded for further data analysis. Blank and double loaded replicates were manually removed.

Frames for which food or flies prediction likelihood was below 0.8 were removed. Foraging performance was evaluated based on “normalized distance to food” (mean log-transformed distances from food normalized by mean log-transformed locomotion of fly in order to account for locomotion differences between flies), “times approaching food” (number of times fly comes within 20% of total area radius to food after being > 25% of arena radius from food), and “averaged locomotion” (mean pixels traveled per unit time).

### Ant Foraging assay

For assaying foraging we updated our previous assay(*7, 8*) to accommodate automated tracking via DeepLabCut as implemented for fly analysis. 8 treated Major and 8 Minor workers (age day 3-5) were isolated in a two-chamber foraging assay, with one box containing moist plaster and shielded from light (simulated nest) and the foraging arena exposed to light and dry, connected by a food tube. Food was provided as 20% sucrose solution applied to a cotton ball at the center of the foraging box. Video was recorded each dark cycle (8pm-8am) for 10 days using night vision security camera (Teckin TC100), as *C. floridanus* forages primarily at night (*7, 166*). Videos were inspected for artifacts, and videos for which tracking failed were discarded.

### Ant foraging data analysis

Four foraging arenas from one batch of experiment were first cropped as individual videos using VideoProc. Individual ant locomotion and interaction with food were tracked using DeepLabCut (version 2.2rc2, multi-animal mode) (*165*). A demo video with 5 Major and 5 Minor workers moving in the foraging arena chamber was used as a training video dataset. Two separate dlccrnet_ms5-based neural networks were trained to track Major and Minor workers separately. Each Major (or Minor) were labeled as a “individual” with 8 “bodyparts” (3 at head, 2 at thorax, 3 at abdomen) and cotton ball “food” was labeled as a static “object”. 70 (or 50) frames were labeled and 100,000 (or 200,000) iterations were applied in Major (or Minor) training. All parameters in “pose_config.yaml” are default. Combined evaluation result output for Major training: Shuffle number =1, Train error (px) = 1.69, Test error (px) = 2.43, p-cutoff used = 0.6. Combined evaluation result output for Minor training: Shuffle number =1, Train error (px) = 2.05, Test error (px) = 2.1, p-cutoff used = 0.6. The coordinate of head bodypart1 of each individual at each frame was recorded for further data analysis.

Frames for which food or ant prediction likelihood was below 0.8 were removed. Foraging performance was evaluated based on mean number of workers present in foraging arena per night (normalized by number of workers per replicate), and “times approaching food” (number of times an ant comes within 20% of total area radius to food after being > 25% of arena radius from food). “Averaged locomotion” was calculated as mean pixels traveled per frame for all ants when ants were detected, to evaluate contribution of locomotion differences to observed data. Minor workers were analyzed from replicates for which Major workers were injected with GFP dsiRNA controls.

### Brain immunostaining and imaging

Immunostaining protocol was adapted from previous study with minor modification on detergent for gentler permeabilization (*41*). Freshly dissected *C. floridanus* brains in DPBS were fixed in 4% paraformaldehyde in PBST (1xDPBS + 0.1% Tween-20) overnight at 4°C and washed in DPBS for 3 times before embedding in 4% agarose gel. 100μm cross-section were performed using Vibratome (Leica VT1000S) at level-7 vibration and level-3 speed, sectioned samples were then transferred to 48-well plate for further staining. Sections were permeabilized in PBST (1xDPBS + 0.5% Tween-20) for 30 mins and blocked in 5% goat serum in PBST (1xPBS + 0.1% Tween-20) for 1 hour. Then sections were incubated with the following primary antibodies diluted in blocking buffer overnight at 4°C: 1:20 mouse anti-Synapsin (3C11, DSHB); 1:100 custom rabbit anti-*cf*Jhe. Before secondary antibody staining, samples were washed 3 times for 30 min in DPBS. Secondary anti-mouse immunoglobulin G (IgG) (H+L), Alexa Fluor 568 and anti-rabbit IgG (H+L), Alex Fluor 488 (ThermoFisher) was 1:1000 diluted in DPBS and incubated with samples overnight at 4°C. Then all samples were washed in DPBS 3 times for 10min, counterstained in DAPI (2ug/ml in DPBS, Thermofisher), and washed in DPBS 3 times for 10min again. Last, samples were mounted in antifade mounting medium (Vector Laboratories, H-1000). For *melanogaster* GFP overexpression lines, freshly dissected fly brains were fixed in 4% paraformaldehyde in PBST (1xDPBS + 0.5% Tween-20) for 20min. Washing, DAPI counterstaining and loading procedures were the same as *C. floridanus*.

Slides were imaged with a Leica SP8 laser scanning confocal microscope (Penn CDB microscopy core). Default imaging parameters including “laser intensity” and “gain” and “offset” were calibrated using Day0 Major condition (for ants) or “repo-G4 > UAS-CD8::GFP” (for fly). Whole mount images (Fig.2G column 1; Fig. 3G row 1) were generated using ImageJ “image -> Stacks -> Z project -> sum slices”.

### Hemolymph extraction

Day0 Major or Minor workers were anesthetized in ice, and head was cut by dissection scissors to create a hole from both head and thorax side. Forceps were used to gently squeeze the head and thorax parts and drops of clear hemolymph squeezed out from the hole was collected by a 10ul pipette. Overall, 3.5-4ul clear hemolymph (pooled from head and thorax, no visible debris or opaque fat body contamination) can be collected from a Major and 0.2-0.3ul from a Minor. Fly hemolymph collection was done using method described before (*167*). Briefly, 30-35 anesthetized adult male drosophila (day 1-5) were punctured in the thorax using a fine glass needle and placed in a 0.6ml tube with a hole at bottom poked by a 26 gauge needle. The 0.6ml tube was then put in a 1.5ml collection Eppendorf tube and centrifuged at 5500rpm at 4C for 5mins. About 1-1.5ul hemolymph can be collected from each batch.

### Brain and hemolymph Western blot

A single *C. floridanus* brain or 3 *D. melanogaster* brains were dissected and homogenized in 100ul of RIPA lysis buffer (150mM NaCl, 1% NP-40, 0.5% sodium deoxycholate, 0.1% SDS, 25mM pH 7.4 Tris) with 1X cOmplete protease inhibitor cocktail (Roche) and incubated on ice for 30 min. Fly or ant hemolymph was first diluted up to 50ul in PBS and centrifuged for 5min at 500g to pellet cells and debris. Supernatant was 6:1 mixed with 7X RIPA buffer with protease inhibitor and incubated on ice for 30min. Lysate was centrifuged at 15,000*g* for 20 min at 4C and supernatant was transferred to a new tube. Protein concentration was measured using pierce TCA protein assay kit (ThermoFisher) to estimate loading volumes. Antibodies used were as follows: 1:2000 mouse anti-β-tubulin (E7, DSHB); 1:300 custom rabbit anti-*cf*/dmJhe; 1:5000 anti-rabbit/anti-mouse IgG (H+L) (Jackson Immune Research). Band intensity (normalized to beta-Tubulin) was measured using ImageJ. Mann Whitney test was used for statistical comparison.

### In-vitro Jhe Overexpression and secretion ratio analysis

Full-length *cf*Jhe-HA, *dm*Jhe-HA, and HA- mCherry-nls were cloned in an insect-derived cell line overexpression vector pB_hsp70 under control of a constitutive hsp70 promoter for transient transfection. To overexpress chimeric Jhe or mCherry using the same vector, DNA sequence of the non-homologous N-terminus of *cf*Jhe (1-61 a.a.) and d*m*Jhe (1-41 a.a.) were swapped between dm/*cf*Jhe or conjugated to the N-terminus of mCherry-HA. Drosophila S2 cells grown in Schneider’s insect medium (Gibco) with 10% FBS were first 1:1 passaged using serum free sf-900 II SFM medium (Gibco) for adaptation. Cells were passaged again and distributed into 12-well plates (1ml per well) 1-2 days before transfection without changing composition of medium. Transient transfection was performed using TransIT-Insect transfection reagent following the manufacturers recommendations (1ug DNA: 2ul TransIT reagent per ml of cell culture). Each condition was assessed using three replicates each. After 6 hour of transfection, media was changed to 100% serum free sf-900 II SFM medium and cultured for an additional 48 hours. 900ul of cells were harvested and cell pellet and supernatant were separated by centrifugation (300g, 5min).

Cell pellets were washed once with DBPS and lysed in 90ul RIPA buffer with proteinase inhibitor using same method as for WB. 900ul of medium supernatant was filtered (0.45μm syringe filter, Milipore), centrifuged (15,000rpm, 15min), concentrated to 45ul (10K or 30K amicon ultra column, Milipore) to which 45uL of 2x RIPA buffer was added. Equal volumes of cell lysate and concentrated medium were loaded and compared via WB. Antibodies used were as follows: 1:1000 mouse anti-HA.11 epitope (16B12, BioLegend); 1:2000 mouse anti-β-tubulin (E7, DSHB); 1:5000 anti-rabbit/anti-mouse IgG (H+L) (Jackson Immune Research). “proportion secreted” was calculated based on the band intensity of HA-tagged protein in media samples divided by total HA-tagged protein measured in both media and cell lysate samples. Band intensity was measured using ImageJ. Mann-Whitney test was used for statistical comparison.

### RNAi brain Injections

Two individual dsiRNA complexes (IDT) targeting *cf*Jhe and two post KD timepoints (12 hour vs 24 hour) were tested. We determined that two dsiRNA were equally effective and 12 hour was sufficient for efficient KD (Figure S10A). RNAi injection method was as done previously (*7, 8*). Prior to injection, *cf*Jhe and GFP- targeting dsiRNA’s were complexed with Polyethelenimine (in vivo JET-PEI, Polyplus) transfection reagent to improve delivery of dsiRNA to the cytoplasm. PEI: RNA Complexing 10 uL of 20uM stock dsiRNA was mixed with 10uL 10% glucose to yield a 5% glucose+dsiRNA solution. In a separate tube, 1.6 ul of in vivo-jetPEIwas mixed with 10uL 10% glucose and 8.4uL sterile water. The 20uL in vivo jetPEI and 20uL dsiRNA were combined in a new tube and incubated for 15 minutes at room temperature prior to injection. DsiRNA:PEI complexes were made fresh for each batch of injections and never stored.

Ant was anesthetized in ice for less than 5 min before injection. 1uL of dsiRNA:PEI solution was injected per ant, and ants were then returned to a group-housed temporary colony with food and shelter for 12 hours, followed by dissection for RNA-seq.

### JH3 gaster injections

JH III (Sigma) was resuspended in DMSO at 5mg/ml and stored in -80°C. A gradient JH III gaster injection concentration was tested and 200ng/ul appeared to be sufficient at 12 hour via RT-qPCR assessment of induction of Kr-h1 (Fig. S10B). Before injection, JH III was first diluted to 10x concentration in DMSO and then 1: 10 diluted in 5% glucose. JH III at working concentration as well as DMSO control (10% DMSO diluted in 5% glucose) were incubated for 15 min at room temperature. Prior to injection, ants were anesthetized in ice for less than 5 min. 1ul of JH III or DMSO control was injected between the 2^nd^ and 3^rd^ tergite, in order to approximate elevated hemolymph JHII levels. Ants were then returned to a group-housed temporary colony with food and shelter for 12 hours, followed by dissection for RNA-seq. Replicate pairs of cfJhe KD and control (GFP KD) were taken from multiple colony backgrouds, matched between conditions, and this was used in analyses to block for colony-of-origin effect (below).

### JH3 Extraction and Purification

Frozen brains were cryogenically ground using a 5/32” diameter stainless steel ball for two milling cycles, each at 25hz for 30s. Ground brains and hemolymph were vortexed with extraction buffer and incubated on ice for 10min. Extraction buffer consisted of 40:40:20 methanol:acetonitrile:H2O with 0.5% formic acid and 0.2ug/ul valine-d8 internal standard (Toronto Research Chemicals). After incubation on ice, enough 15% (w/v) ammonium bicarbonate (NH4HCO3) in H2O was added to yield a 1.15% (w/v) final concentration in sample extracts(*168*). Extracts were again incubated on ice for 20 minutes and centrifuged twice at max speed (4350g) for 30 min at 4℃ to remove solid tissue fragments.

### Liquid Chromatography Mass Spectrometry Analysis

LC was performed using a Xbridge BEH amide HILIC column (Waters) with a Vanquish UHPLC system (Thermo Fisher). Solvent A was 95:5 water: acetonitrile with 20 mM ammonium acetate and 20 mM ammonium hydroxide at pH 9.4. Solvent B was acetonitrile. The gradient used for metabolite separation was 0 min, 90% B; 2 min, 90% B; 3 min, 75%; 7 min, 75% B; 8 min, 70%, 9 min, 70% B; 10 min, 50% B; 12 min, 50% B; 13 min, 25% B; 14 min, 25% B; 16 min, 0% B, 21 min, 0% B; 21 min, 90% B; 25 min, 90% B. MS analysis was performed using a Q-Exactive Plus mass spectrometer (Thermo Fisher) in positive ionization mode with 10 µL injection volume. An m/z range of 70 to 1000 was scanned. All data were analyzed with El-MAVEN and for experiments involving deuterium labeling, outputs were corrected for natural deuterium abundance using accucor package in R (*169*). Peak areas integrated in El-MAVEN were normalized to valine-d8 internal standard to account for variation in injection volume. The absolute amounts of JH3 and JH3-d3 in each sample were estimated by least squares interpolation of normalized peak areas against standard dilution curves of JH3 and JH3-d3 (Toronto Research Chemicals) which were included in each LC-MS run.

### RT-qPCR

Each single datapoint represented a single ant brain or fly brain. All conditions had ≥5 biological replicates and sampled from 3 distinct colony backgrounds. cDNA was prepared from total RNA using the High-Capacity cDNA Reverse Transcription Kit (Applied Biosystems). Abundance of specific mRNA transcripts was estimated using Power SYBR Green PCR Master Mix (Life Technologies) on a Real Time qPCR machine (Applied Biosystems 7900HT). Relative transcript abundance was estimated using the **ΔΔ**Ct method. Briefly, this method involved the use of the housekeeping gene, RPL32, as an internal normalization control for each sample. Following normalization to this gene, the differences in Ct values between normalized Ct amounts (**Δ**Ct) were taken to obtain **ΔΔ**Ct values, which were converted into fold-change values by the function 2~-**ΔΔ**Ct = F.C. (*170*). Statistical significance was calculated using Mann Whitney test (two group comparison) Kruskal-Wallis test followed by Dunn’test (3 group comparison). Oligo sequences were listed in Table S14.

### Bulk RNA-sequencing

For each sample type, five distinct colony backgrounds were used, in order to subsequently control for inter-colony variation. Individual ants were immobilized on ice for 5 minutes before brains were dissected, rinsed twice in chilled sterile Hank’s balanced salt solution media (HBSS), transferred to 1.5-ml microcentrifuge tubes containing 15 μl of chilled HBSS, and immediately snap-frozen in liquid nitrogen. Total RNA was purified from individual brains by trizol extraction, followed by DNase treatment using TURBO DNase (Invitrogen). DNase-treated RNA was subsequently purified using RNase-free Agencourt AMPure XP beads (Beckman Coulter; 2:1 volume of beads:sample).

For RNA-seq, polyadenylated RNA was purified from total RNA using the NEBNext® Poly(A) mRNA Magnetic Isolation Module (NEB E7490) with on-bead fragmentation as described (*171*). cDNA libraries were prepared the same day using the NEBNext® Ultra™ II Directional RNA Library Prep Kit for Illumina® (NEB E7760). All samples were amplified using 8 cycles of PCR.

### scRNA-sequencing

Single-cell dissociation was adapted from previous study with minor optimization for ambient RNA removal (*41*). In each replicate, 3 Major or Minor ants were pooled, and samples were taken from at distinct colony backgrounds (5x for d0 data, 2x for p18 data) to control for inter-colony variation, with each Major/Minor replicate pair taken from the same background. Brains were dissected in an optimized neuronal culture medium (60% DMEM + 40% neurobasal medium) with 45μM actinomycin D, debris, connective tissues and optic lobes were carefully removed. Dissected brains were enzymatically dissociated by papain and then mechanically dissociated by pipetting. To prepare active papain dissociation buffer, 5ml of dissection buffer was added in one vial of papain (PAP2, Worthington Biochemical, cat# LK003176) and incubated at room temperature for at least 30mins. Adult brains were rotated in 1ml of papain dissociation buffer for 12 min and pupal brains were rotated for 8 min. Brains were then washed 3 times with 1ml cold DPBS + 0.01% BSA and 200ul was left after the last wash. Brains were then slowly pipetted up and down on ice 20-30 times using a 200ul tip, 20-30 times using a 200ul gel loading tip and 20-30 times using a 200ul tip. No visible mass of tissue was expected after dissociation. 1ml of DPBS + 0.01% BSA were added in each vial and dissociated cells were slowly pelleted at 300g for 10min at 4°C. Supernatant was carefully removed without touching the cell pellet. We observed a significant decrease of ambient RNA level after performing one round of washing. Pellet was resuspended in 250ul DPBS + 0.01% BSA. Cell suspension was then filtered on a 40-μm Flowmi strainer 2 times, yielding approximately 180ul. Cell concentration was assessed via trypan blue staining and counting with a hemocytometer. This approach yielded approximately ∼50K cells per Major brain and ∼40K cells per Minor brain.

scRNA-seq was performed using Chromium Single Cell3’Reagent kit v3 (10x Genomics) following user guide rev C. 16,000 cells were loaded on to the microfluidic chip with the goal to recover 10,000 cells for each sample. cDNA and library were both amplified for 10 cycles. Library size distribution and concentration were measured using BioAnalyzer High Sensitivity DNA kit (Agilent) and NEB quantification kit. 2-3 libraries shared one 75-cycle high-yield Illumina flowcell and were sequenced on an Illumina Nextseq 550. 2.8pM library was loaded and the read configuration was 28 (read 1), 8 (index 1) and 56 (read 2).

### Adaptation of reference annotation

Because we observed a large number of genes which possessed truncated 3’UTRs in the *C. floridanus* NCBI annotation relative to aligned brain RNA-seq data, we sought to globally amend annotations to account for this. In order to create this adapted NCBI annotation, we first aligned all reads from this study and two others (*8, 126*) as below, but in the second round of alignment included the option “--outSAMattributes NH HI AS nM XS”. We then ran isoSCM v2.0.12 (*172*) assemble with the options “-merge_radius 150 -s reverse_forward” on all alignments. Following this we extracted only 3’ exons from the resulting draft annotations, which were then overlapped with the current NCBI annotation release. We ten took the longest predicted 3’ exon/UTR that overlapped with the current NCBI version’s 3’ exon, and extended this exon according to the isoSCM 3’ exon. Any predicted 3’ exon extension that overlapped an exon of another gene on the same strand was trimmed to 500 bp upstream of the other gene’s offending exon.

### scRNA-seq analysis

For scRNA-seq counting within cellranger (v3.0.2) we utilized a custom adaptation of the NCBI RefSeq annotations with extended 3’ UTRs as described above, due to the fact that 10X scRNA-seq data is highly biased toward the 3’ end of mRNAs and we observed considerable data recovery with this amended annotation when examining bulk RNA-seq data. We then used Seurat (*38*) to perform clustering and differential testing.

For clustering we only included cells with nFeature_RNA > 500 and < 5,000, and with < 5 % mitochondrial reads. We next used the SCTransform method within Seurat to combine samples, using 3,000 integration features, regressing out % MT reads per-cell, and excluding certain genes (see below). We then selected 2,500 variable features for PCA, but excluded a list of 1,504 genes related to mitochondrial function, tRNA synthesis, cellular respiration, glycolysis and ribosomal functions (Table S15). This was done to obtain consistent clustering across samples, guided by cell type rather than metabolic status.

PCA and UMAP clustering were performed using the top 50 principal components, neighbors were found using FindNeighbors with the parameters k.param=30 and prune.SNN=0.06667. Clusters were determined using a resolution of 0.8 and the Louvain algorithm. For determination of per-cluster caste differential genes, the FindMarkers function of Seurat was run on each cluster, comparing cells from Majors and Minors using the wilcox test. For cluster annotation both the FindAllMarkers Seurat function as well as the soupX package’s (*173*) quickMarkers function were used to determine cluster markers, then top hits from these were examined and compared to prior literature.

In order to ensure specific clusters more reflective of actual cell subtypes Default Seurat clusters at resolution 0.8 were used, but FindClusters was run using resolutions 0.05, and 0.1, 0.25, 0.5 and 0.8, and resulting clusters were used in Seurat’s BuildClusterTree, clustree (*174*), along with visual inspection of top markers and assessment of SoupX markers (comparing focal cluster to ‘second best’ cluster) for each 0.8 resolution cluster. For several clusters which appeared to be artificial splits based upon shared markers, the clustree hierarchy was used along with shared markers to merge clusters prior to naming (Table S2, first column).

For comparing *C. floridanus* d0 scRNA-seq clusters with *H. saltator*, we utilized the data from (*41*) but with updated clustering. Briefly, *H. saltator* 10x data was filtered to only include cells with nFeature_RNA > 500 and < 5,000, and with < 5 % mitochondrial reads. SCTransform method within Seurat to combine samples, using 3,000 integration features excluding certain genes. We then selected 2,500 variable features for PCA, excluding genes given in Table S15. This was done to obtain consistent clustering across samples, guided by cell type rather than metabolic or hormonal status. PCA and UMAP clustering were performed using the top 30 principal components, and clusters were determined using a resolution of 1.0 and the Louvain algorithm. For comparing between species, *H. saltator* data was limited to only genes sharing a 1-to-1 orthologous relationship with *C. floridanus*, and renamed using *C. floridanus* 1-to-1 ortholog gene IDs. The Seurat function FindTransferAnchors was used to establish transfer anchors between *C. floridanus* and *H. saltator* datasets using previously-established variable features and the “cca” method, as we found this approach (vs pcaproject) to more robustly assign cell types across the >100my of divergence between ant species. *C. floridanus* cells possessing a prediction score >0.5 were retained, and assigned to their respective *H. saltator* cluster.

For analysis of pupal scRNA-seq data, pupal scRNA-seq caste-specific datasets (2x replicates per caste) were filtered as with d0 data, and both d0 and pupal data were combined for a d0-centered analysis. SCTransform and integration features were performed and selected as with d0 data (above) as was PCA). Due to observation of batch effects not eliminated by SCTransform, for d0+pupa data we followed this with integration using harmony (*175*) with the Seurat function RunHarmony, controlling for batch and caste on the SCTransform data and the parameters (“kmeans_init_nstart=20, kmeans_init_iter_max=100, max.iter.harmony=20, max.iter.cluster=30”). UMAP clustering and neighbor finding were done using this harmony-generated reduction using the top 50 principal components. Clusters were determined using a resolution of 0.8 and the Louvain algorithm. Markers, caste and stage differing genes were determined as for d0-only data. For inference of d0+pupal cluster identities, d0 cell barcodes were used to lift d0 cluster annotations over to the combined dataset.

For TF clustering (Fig. 1F) average per-cluster expression for all genes possessing an interpro-predicted DNA binding domain. These were further filtered to include only genes determined as cluster markers via SoupX’s quickMarkers function, and only those genes showing >1.5 fold higher expression in the cluster they marked vs the second most strongly marked cluster, as well as expression in >50% of cells in the focal cluster.

For Gene Ontology clustering (Fig. S3), for each cluster the top 200 marker genes via SoupX’s quickMarkers function were selected based upon TF-IDF and functional enrichment (BP) was tested between these 200 markers compared to a list of genes that were seen among the top 400 markers for any other cluster (via the same method). The resulting per-cluster GO test results were then filtered to include only terms showing a p-value > 0.001 in at least one cluster, to produce a list of GO terms significant in at least one cluster. A matrix of P-values for each Term X cluster was generated, and each cluster’s p-values for each term was incorporated (including non-significant p-values) and –log10 transformed. This matrix of Term X cluster p-values was then used to generate a heatmap.

For UMAP significance plotting of gene overlaps between scRNA-seq caste DEGs and bulk RNA-seq DEGs, for each cluster DEGs between Major and Minor cells were established as above, and these were overlapped with lists of bulk RNA-seq DEGs, followed by significance testing via fisher’s exact test using only genes testable (with sufficient data) in both datasets as a background. Per-cluster p-values were –log10 transformed and each cell in the given cluster was given this value. UMAP plots were then generated using these values. Clusters for which no significance could be established (no overlap) were given a value of 0.

For gene regulatory network analysis (Figure S5), the full cell X gene expression matrix was exported from Seurat and used with pySCENIC (*57*) to generate gene adjacencies via GRNboost2 (*176*), and these adjacencies used to calculate expression modules (modules_from_adjacencies) for all genes annotated with a significant hit to a DNA binding domain via (*177*) . This was followed by the determination of ‘regulons’ (regulatory unis of a given TF) using a custom database of ranked gene X motif signatures, determined using Cluster-Buster (*178*). Briefly, motif PWMs were used from fLyFactorSurvey (*179*), idmmpmm (*180*), and On-The-Fly (*181*), converted to Cluster-Buster format, and used to scan the promoters (-4kb and +0.5kb from start of gene model) of all *C. floridanus* genes. The given motif PWMs were annotated with the reciprocal best hit ortholog from ants (NCBI LOC ID), and in the case of fly gnees lacking a RBH hit in *C. floridanus*, *C. floridanus* genes were blasted against those of fly, and hits with E-value < E-10 were taken and used to link these remaining genes to their PWMs, and other PWMs lacking an ant ortholog was discarded. These were then used to prune regulatory modules (pySCENIC prune2df, nes_threshold=2) into ‘regulons’.

Both modules and regulons were then compared to marker lists for each celltype by overlapping the regulated genes for each module and regulon with markers for each cluser (top five for each given in Table S3). Due to the limitations of using fly motifs to annotate ant genes, we found very few filter-passing regulons and that downstream regulon-focused clustering to be uninformative. Thus we only chose to present and include top modules and regulons.

For assessment of cluster representation between sample types (Fig S7B and S7E), individual replicate cell numbers for each cluster were determined using Seurat, and for each cluster, a binomial generalized linear model to test differential representation between caste (Fig S7B), or caste + stage (Fig S7E), modeling cluster cell number (vs total cell number) by noted categorical variables. Resulting per-cluster p-values were then corrected for false discovery using the Benjamini-Hochberg method.

### Bulk RNA-seq analysis

Reads were demultiplexed using bcl2fastq2 (Illumina) with the options “--mask-short-adapter-reads 20 - -minimum-trimmed-read-length 20 --no-lane-splitting --barcode-mismatches 0”. Reads were trimmed using TRIMMOMATIC (*182*) with the options “ILLUMINACLIP:[adapter.fa]:2:30:10 LEADING:5 TRAILING:5 SLIDINGWINDOW:4:15 MINLEN:18”, and aligned to the *C. floridanus* v7.5 assembly (*183*) using STAR (*184*). STAR alignments were performed in two passes, with the first using the options “--outFilterType BySJout -- outFilterMultimapNmax 20 --alignSJoverhangMin 7 --alignSJDBoverhangMin 1 --outFilterMismatchNmax 999 --outFilterMismatchNoverLmax 0.07 --alignIntronMin 20 --alignIntronMax 100000 –-alignMatesGapMax 250000”, and the second using the options “--outFilterType BySJout --outFilterMultimapNmax 20 – alignSJoverhangMin 7 --alignSJDBoverhangMin 1 --outFilterMismatchNmax 999-- outFilterMismatchNoverLmax 0.04 --alignIntronMin 20 --alignIntronMax 100000 --alignMatesGapMax 250000 --sjdbFileChrStartEnd [SJ_files]” where “[SJ_files]” corresponds to the splice junctions produced from all first pass runs. Gene-level counts were generated using the featureCounts program (*185*) with the parameters “-- fraction -O -M - -s 2 –p”and fractional counts were rounded to the nearest integer value, using the extended isoSCM annotation noted above.

Differential gene expression tests were performed with DESeq2 (*186*). For all pairwise comparisons the Wald negative binomial test (test=”Wald”) was used for determining DEGs, using colony background as a blocking factor. Unless otherwise stated, an adjusted p-value cutoff of 0.1 was used in differentiating differentially expressed from non-differing genes in order to maximize the sensitivity of our RNA-seq results. For RNA-seq libraries at least 15M mapped reads were sequenced for each replicate.

For mouse endothelial RNA-seq raw reads were downloaded from GEO (GSE95401) and aligned as above using STAR (mm10), and differential gene expression was assessed using DESeq2, comparing brain endothelial cells to whole brain, as well as to all other endothelial cells’ RNA-seq data. Genes annotated with the functional term “hormone metabolic process” (GO:0042445) were examined for brain endothelial-specific expression, and hits were further examined for prior support for this function.

### Assignment of gene orthology and functional terms

Genes (NCBI Camponotus floridanus Annotation Release 102) were assigned orthology using the reciprocal best hit method (*187*) to both *D. melanogaster* (r6.16) and *H. sapiens* (GRCh38) protein coding genes. Gene ontology function was assigned to genes using the blast2go tool (*188*) using the nr database, as well as interpro domain predictions. Gene Ontology enrichment tests were performed with the R package topGO (*189*), utilizing the fishers elim method. For presented GO results (eg Fig. S1B), GO lists were reduced using ReviGO (*190*), to eliminate redundant terms.

### Esterase phylogenetic tree

For the phylogeny presented in Figure 4A, all fly and mouse genes in the orthoDB (*191*) ortholog group *cf*Jhe was placed within (Carboxylesterase, type B; 206516at33208) were aligned using PRANK (*192*). Poorly aligned regions were filtered using Gblocks (*193*), and the resulting filtered alignments were used to generate a phylogenetic tree using PRANK.

### Prediction of secretion probability

For Jhe family genes presented in Figure 4C, full a.a. sequence of each genes were uploaded to Signal5.0 server (*194*) to locate signal peptide and calculate secretion probability.

### Statistical testing

For gene overlap calculations (such as in Fig. 5B) the GeneOverlap (*195*) R package was used. All other statistical tests given in figures and tables were performed in R and GraphPad Prism unless otherwise stated. For all experiments using ants, colony background was recorded and balanced between compared castes or conditions. For bulk RNA-seq, behavior analyses, as well as JH3 quantification, colony background was entered as a blocking factor for analyses in order to statistically control for colony-of-origin effects in our data.

### Data availability

All sequencing data related to this project have been deposited in the NCBI Sequencing Read Archive under the BioProject PRJNA894104.

## Author Contributions

K.M.G., L.J., J.G. and S.L.B. designed the experiments. K.M.G. and L.J. collected the data. K.M.G. and L.J. analyzed the data. LC-MS was run and analyzed by C.S.K., S.M.D., and S.D.K. L.S., J.G. and B.Z.K. provided technical support. K.M.G. L.J. and S.L.B. wrote the manuscript with input from all co-authors.

## Acknowledgements

We thank members of the Berger Lab for help in editing the manuscript; Sehgal Lab and Bonasio Lab for providing fly lines; Geoffrey Dann (Garcia Lab) for providing help on antigen purification; and Duo Zhang for designing and printing fly foraging arena; and Balint Kacsoh for guidance in initial optimization of DLC video analysis model fitting. This work was supported by NIH training grant F32GM120933 (K.M.G.) and NIA R01 5R01AG055570 (S.L.B.), as well as NSF 1DP2GM137424-01 for S.D.K.

## Supplementary Table legends

**S1.** All cluster markers for d0 scRNA-seq data as identified by both Seurat’s FindAllMarkers, as well as SoupX’s QuickMarkers function. For each along with statistics of cluster-enrichment, reciprocal-best orthologs for fly, human, and *H. saltator* are given, as is interpro scan-determined DNA binding domain status and type.

**S2.** Table of cluster names, including initial Seurat cluster number, annotation long-name, overall celltype class, and the simplified UMAP names given to each cluster in figures and tables.

**S3.** Results from SCENIC GRN analysis. Top ≤ 5 SCENIC modules/regulons significantly overlapping scRNA-seq clusters, ranked by odds ratio of overlap (for significantly overlapping Regulons/modules only). SCENIC was run on all d0 scRNA-seq data, producing co-expression modules, linked to a given TF (column “Analysis_type” = “SCENIC_MODULES”). The resulting regulated genes were then overlapped with cluster markers for each cluster (as well as major cell groups’ markers: Mushroom body neurons, non-MB neurons, and Glia), and module/regulon TFs were included if they showed significant overlap between their reulated genes and the cluster markers for a given cluster, and were also a significant marker for the given cluster themselves. SCENIC modules were also pruned to ’regulons’ using a custom motif ranking database, generated using fly motifs mapped to *C. floridanus* genes (“SCENIC_REGULONS”). Because this led to overly-aggressive pruning (likely due to insufficiency of fly motifs for annotating ant genes), we chose to include both Regulons and Modules. For SCENIC_REGULONS the contributing genes (those overlapping cluster markers regulated by a given TF) are given in the column “Regulon_intersecting_gene_IDs”. p-values, FDR values, and odds ratios represent the results from a fishers exact test comparing genes regulated by a given TF and cluster markers for the given cluster.

**S4.** Genes showing significant caste bias per-cluster when comparing between Major and Minor cells using Seurat’s FindMarkers, with the Wilcoxon test.

**S5.** Gene ontology germs significantly enriched among marker genes for a given cluster, as compared to a background set of all genes identified as a marker in any cluster, in order to control for brain-specific functions associated with brain-specific gene expression.

**S6.** Gene ontology terms associated with genes showing significant caste bias between Major and Minor workers in either d0 or p18 data. Genes were tested relative to all marker genes seen across all clusters.

**S7.** Statistics for all clusters identified among d0 scRNA-seq data for each library used in this study, including median UMI counts (nCount_med), median number of expressed features (nFeat_med), mitochondrial percent (mtPerc_med) and number of cells.

**S8.** Marker genes identified via QuickMarkers for clusters identified when analyzing d0+p18 data.

**S9.** Genes showing significant caste bias per-cluster when comparing between Major and Minor cells using Seurat’s FindMarkers, with the Wilcoxon test using clusters from p18+d0 combined dataset, and evaluating caste bias between both p18 castes as well as d0 castes in the context of the combined-data clusters.

**S10.** Genes showing significant bias per-cluster when comparing between p18 and d0 cells using Seurat’s FindMarkers, with the Wilcoxon test.

**S11.** All genes identified as significantly differing in bulk RNA-seq data analyzed here, for Jhe KD (*cf*Jhe_KD_DEG_Class) JH3 injection DEGs (JH3_INJ_DEG_Class) and d0 Major vs Minor DEGs (d0_Caste_bias), as well as average FPKM values for each sample type averaged between replicates.

**S12.** Results of gene overlap calculations comparing bulk RNA-seq DEGs to per-cluster scRNA-seq Major-vs- Minor d0 DEGs, providing direction of significant scRNA-seq caste bias, bulk RNA-seq DEG type, and numbers of overlapping genes for each cluster.

**S13.** Table of fly genotypes used here.

**S14**. Table of oligo sequences

**S15.** Genes excluded during scRNA-seq data analysis and clustering (but not differential testing or cluster marker finding) for both *C. floridanus* and *H. saltator*.

**S16.** Table of genetic backgrounds and sample origins for experiments in this study.

## Supplementary Figures

**Figure S1:**
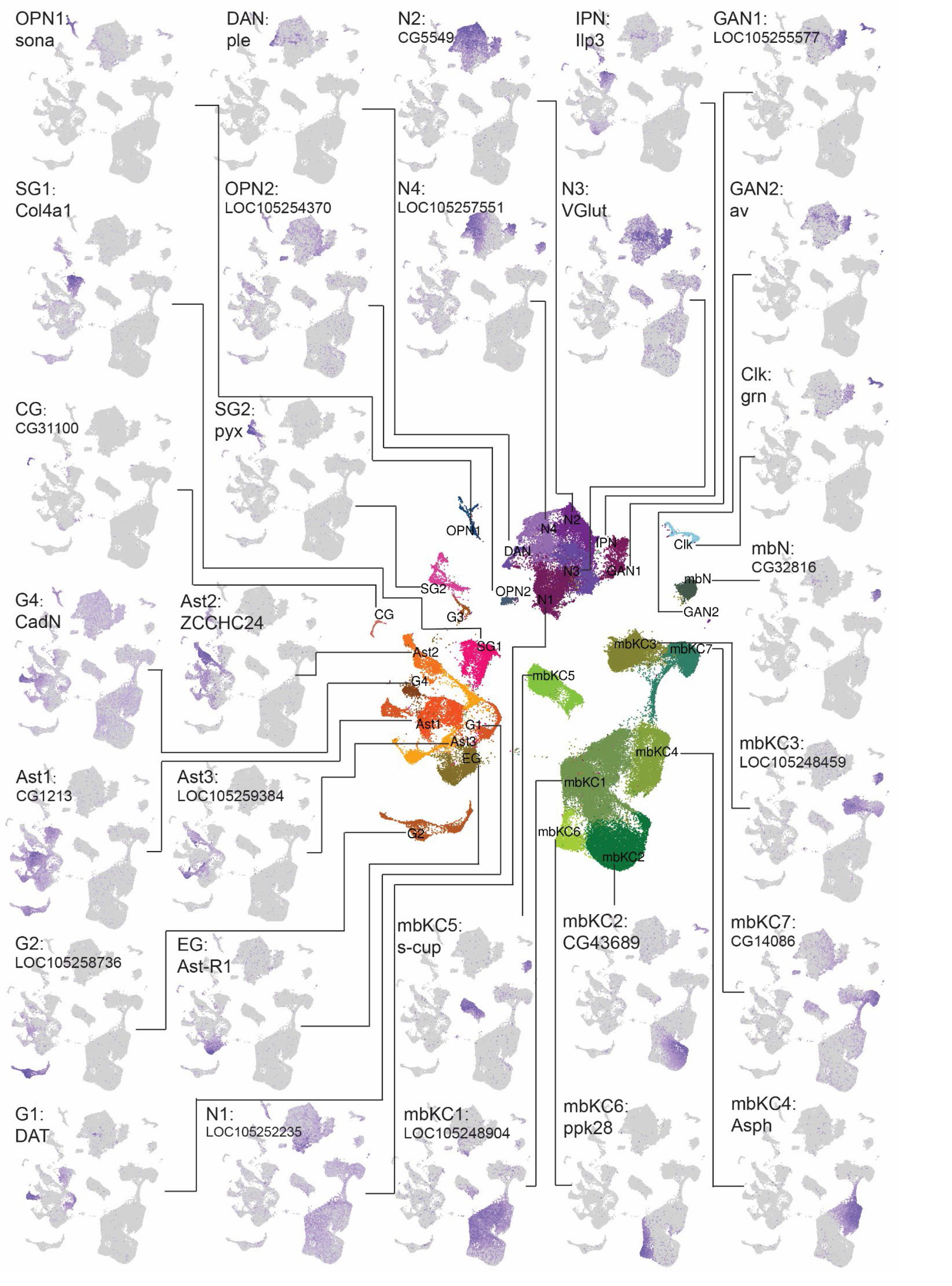
top markers for each cluster, illustrated on UMAP plots. For each cluster the top gene marking the given cluster (by TF-IDF in Table S1) was selected and plotted. Lines connect marker UMAPs to overall colored UMAP (center).

**Figure S2:**
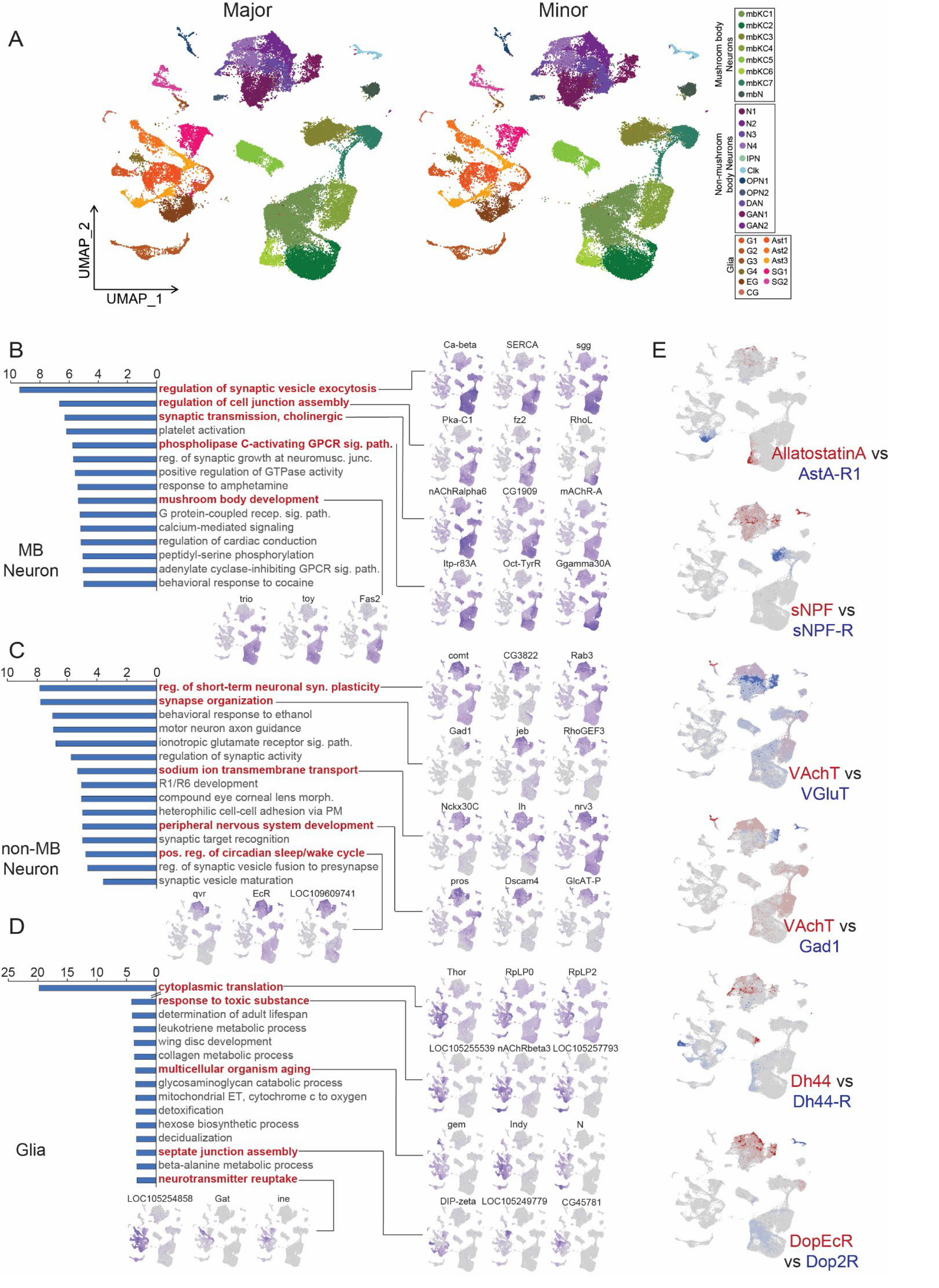
A) UMAP plots of d0 scRNA-seq clustering, showing each caste separately to illustrate generally similar representation in both castes of all major clusters (see Table S7 for full per-sample per-cluster representation). B-D) GO term comparison between MB neuron, non-MB neuron, and glial population marker genes, with informative terms highlighted in red. GO terms represent enriched terms associated with a given cluster’s marker genes as compared to a background of the other two clusters’ marker genes. X-axis on plots represents log10-transformed p-value of the given term’s enrichment among a focal group’s marker genes. E) Co-expression of neuropeptide vs receptor (left columns) and X-ergic markers (right, top two), as well as two classes of dopamine receptors (right, bottom panel).

**Fig S3.**
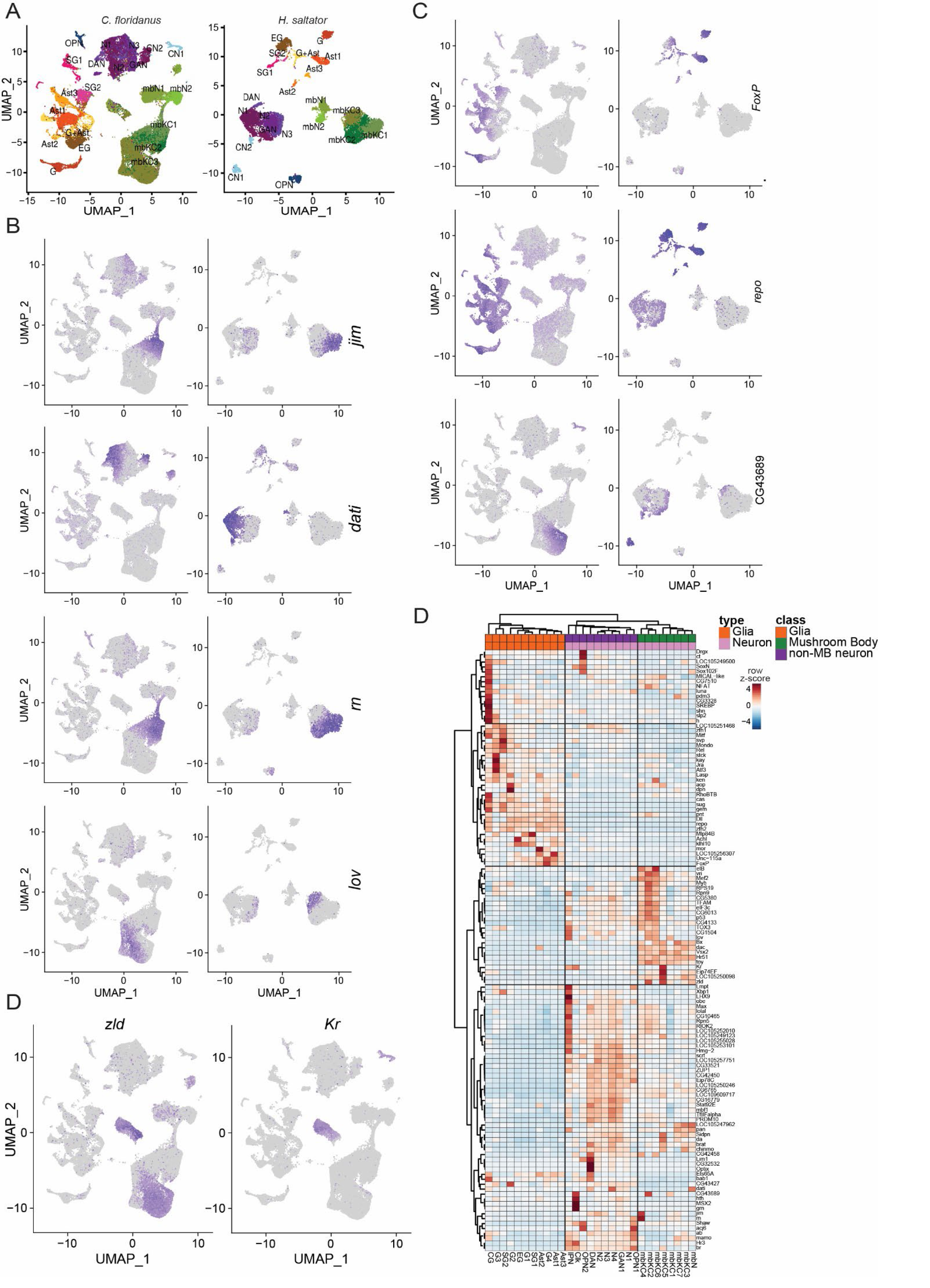
A) Cross-species integration between *C. floridanus* and *H. saltator* scRNA datasets illustrating conservation of major cell types between the two species. Clusters in *H. saltator* were annotated independently based upon cluster-specific markers but named based upon orthologous *C. floridanus* clusters, and *C. floridanus* scRNA-seq data was integrated using only orthologous genes between the two species. B) Genes defining distinct neuronal subtypes in *C. floridanus* and *H. saltator* that were also used as integration features between species. C) Additional genes showing strong homologous cluster-specificity between *C. floridanus* and *H. saltator* for major glial subtypes. D) *zld* and *Kr* expression in d0 scRNA-seq showing co-localization to a single cluster (mbKC5). D) More inclusive TF heatmap as from Figure 1F (TF-IDF > 0.5, >1 fold over 2^nd^ best cluster, >2 fold over global levels, >30% of cells in focal cluster expressing TF).

**Fig S4.**
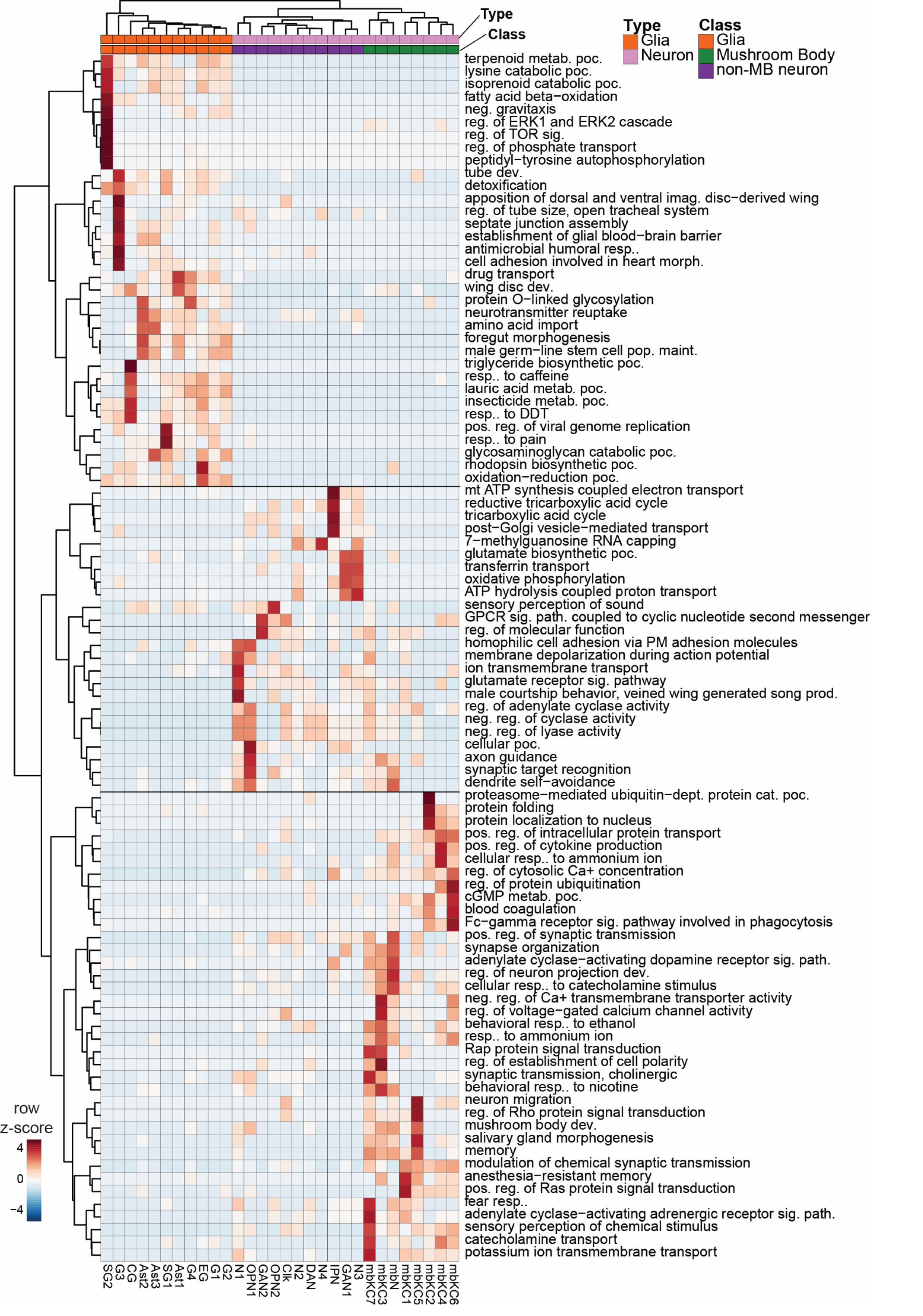
Heatmap of cluster-enriched GO term-based clustering showing segregation of basic main celltypes based entirely on comparison of GO terms, identified by comparing a given cluster’s top marker genes to a background set of all genes seen as markers in any cluster. Top marker genes were defined as thte top 200 markers associated with a given cell cluster, or in the case of clusters with less than 200 markers, all.

**Figure S5:**
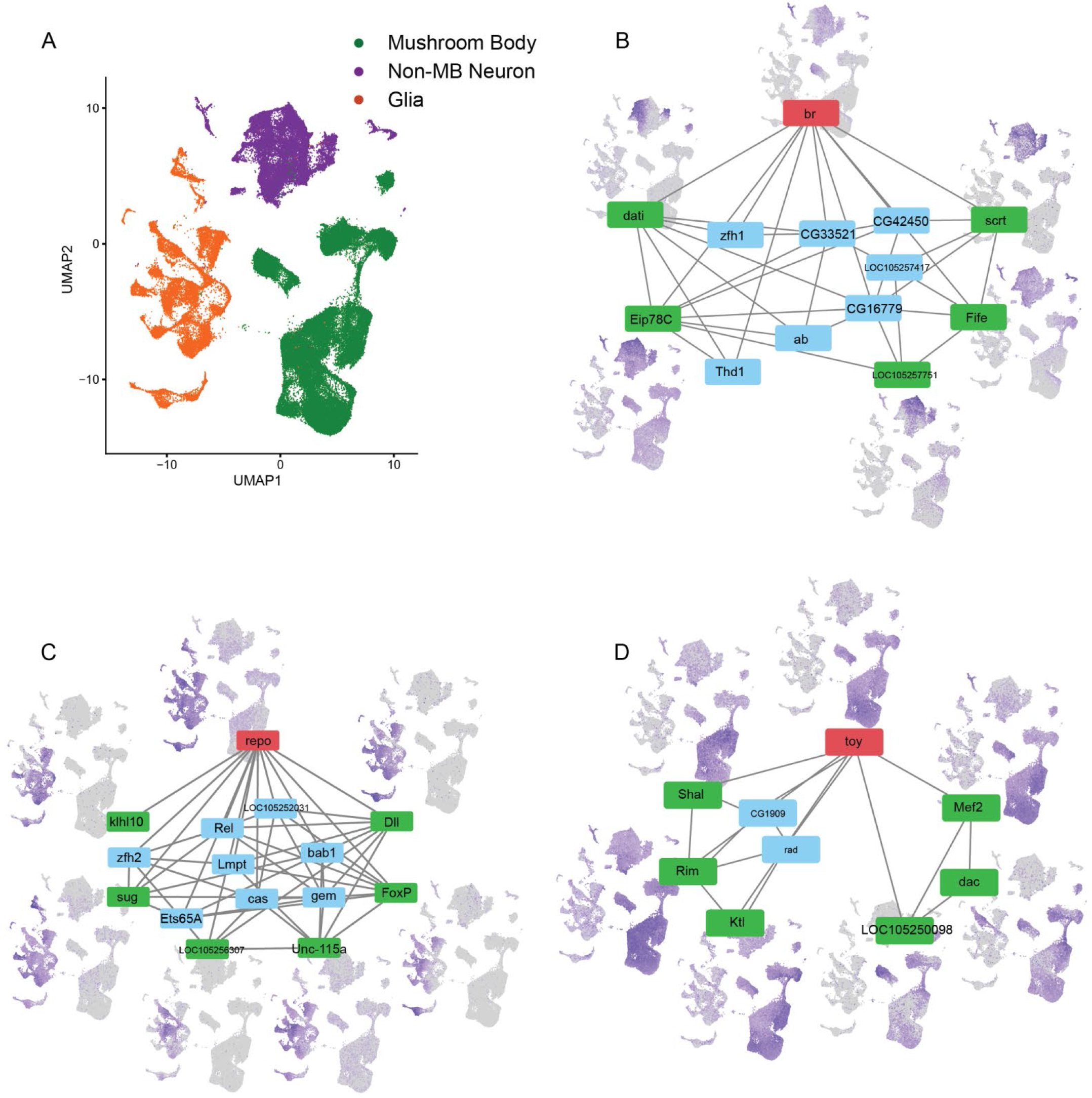
Representative result of gene regulatory analysis on scRNA-seq from *C. floridanus* d0 brain samples. For each major cell group (illustrated in panel A) the top transcription factor regulon is shown as well as its associated TF regulatory network (only showing other TFs downstream of focal TF) are shown, along with UMAP plots of representative downstream TFs showing more sub-cluster specific expression, for B) Mushroom body neurons, C) Glia, and D) Non-mushroom body neurons. The focal parent TF (regulating each network) is shown in red and UMAPs of representative TFs are associated with the green-labeled child nodes. Top regulons were selected by significance of overlap (fishers exact test) between the given major cell group’s marker genes (as determined by Seurat’s FindMarkers function) and the regulated genes of each regulon.

**Figure S6:**
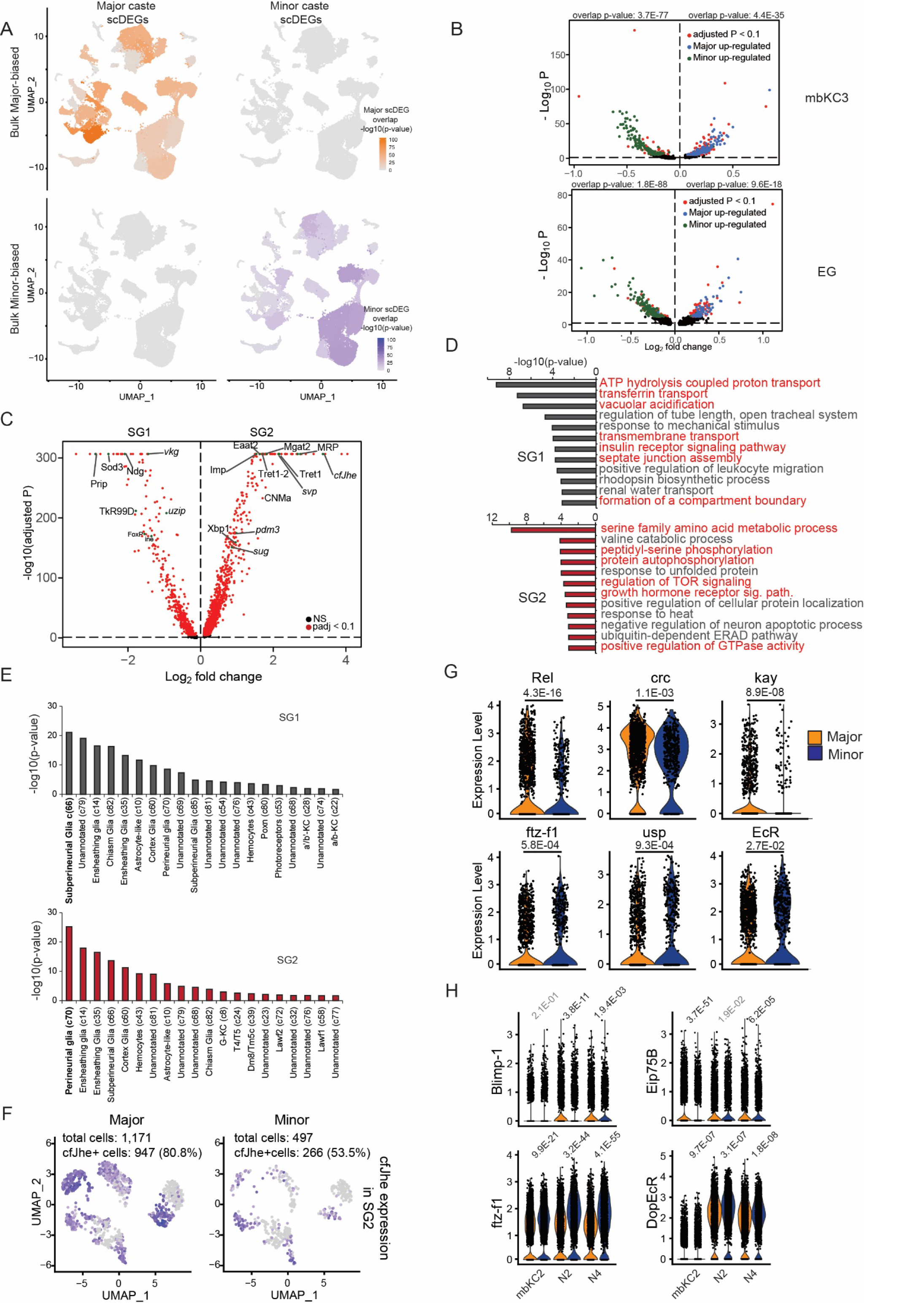
A) Featureplot showing the overlap of d0 bulk RNA-seq significantly caste biased genes and between-caste scDEGs in each cluster based on the –log10 P-value of Fisher’s exact test comparison between bulk DEGs and caste differing scDEGs for each cluster. B) Volcano plot (log2 Major/Minor) of example neuron and glia cluster with strong overlap between bulk DEGs and scDEGs, with overlapping bulk RNA-seq Major-biased DEGs highlighted in blue and Minor-biased DEGs highlighted in green. Caste-biased DEGs for both bulk and scRNA-seq were determined as those with an adjusted p-value < 0.1. C) Volcano plot of DEGs between the two surface glia clusters presented here (log2(SG2 / SG1)), illustrating strong differentiation in gene expression between the two SG clusters. Differentially expressed genes were defined as those showing an adjusted p-value < 0.1 when comparing cells from SG1 to those from SG2, irrespective of caste. D) Top GO-terms associated with genes biased to SG1 (top) or SG2 (bottom) comparing markers of one to a background set of the other, demonstrating functional distinction between the two celltypes markers. Marker genes were defined by comparing SG1 and SG2 cell types, taking those with an adjusted p-value of < 0.01. E) Significance of overlap between ant Surface glia clusters (top: SG2, bottom: SG1) marker genes and cluster-specific markers from all clusters from the fly dataset taken from (*40*). For each fly cluster, marker genes were overlapped with SG-subtype markers. Presented values are the –log10(p-values) of fishers exact tests from these overlaps for the top 20 fly clusters overlapping each ant surface glia subcluster. F) Re-clustering of SG2 cells illustrates higher number of SG2 cells in Major workers, but also much higher percentage of SG2 cells expressing cfJhe independent of cell number differences. G) Violin plot of selected hormonal-signaling related TFs differentially expressed between Major and Minor in SG2. P-values from Seurat differential expression testing (wilcoxon method) comparing Major vs Minor SG2 cells via Seurat. H) Violin plot of two hormonal related TFs differentially expressed between Major and Minor in multiple neuronal clusters (two illustrated here). P-values from Seurat differential expression testing (wilconxon method).

**Fig S7.**
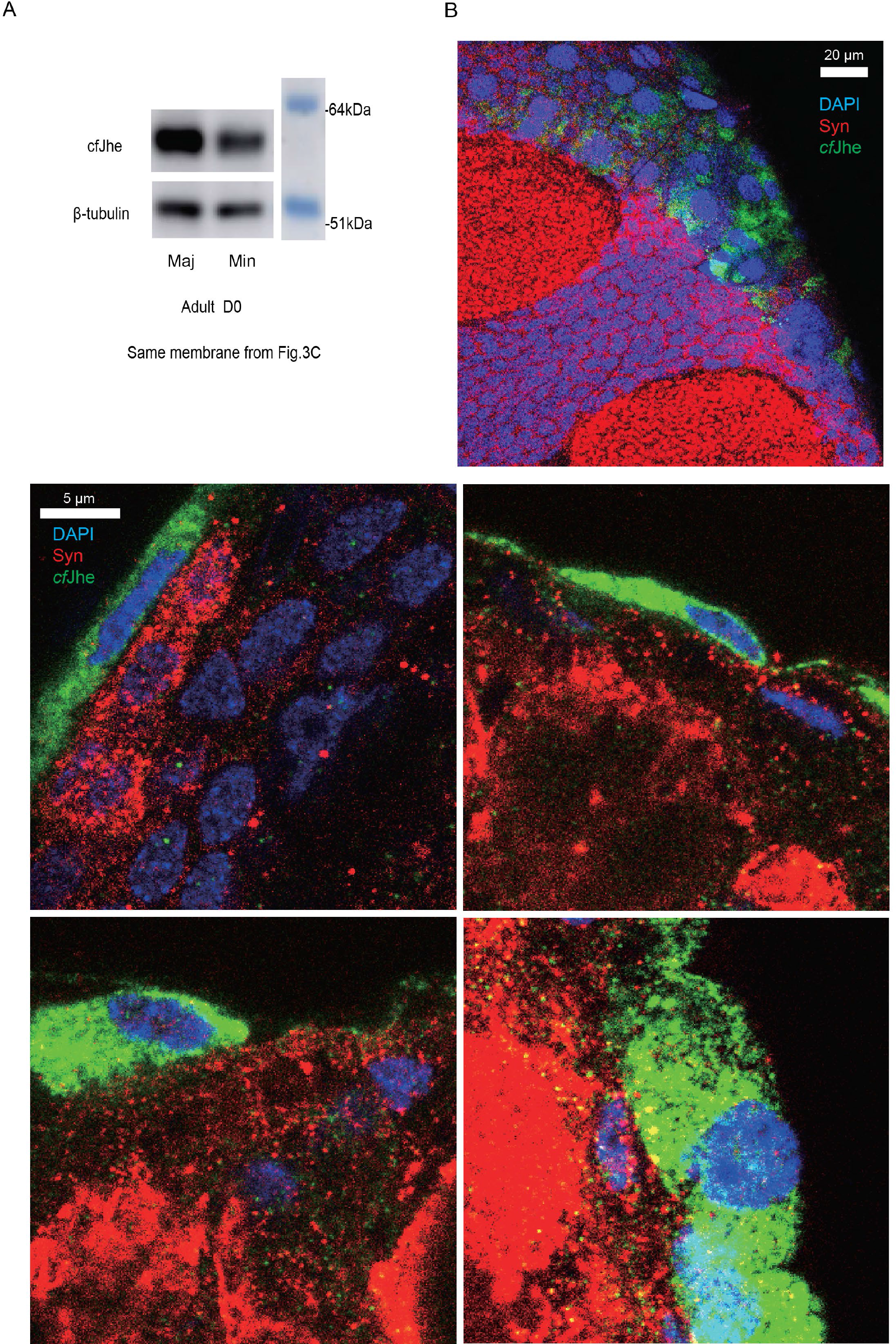
A) Western blot of *cf*Jhe in d0 Major and Minor brain using a custom antibody raised against a *cf*Jhe- derived antigen. B) Upper: Single confocal slice at top of a 100um Major brain section (*cf*Jhe colocalized with the surface glia nuclei with a typical “large and flat” shape). Lower: 100um Major brain sections imaged under 63X objective illustrating single-cell morphology from different sectioning angles.

**Figure S8.**
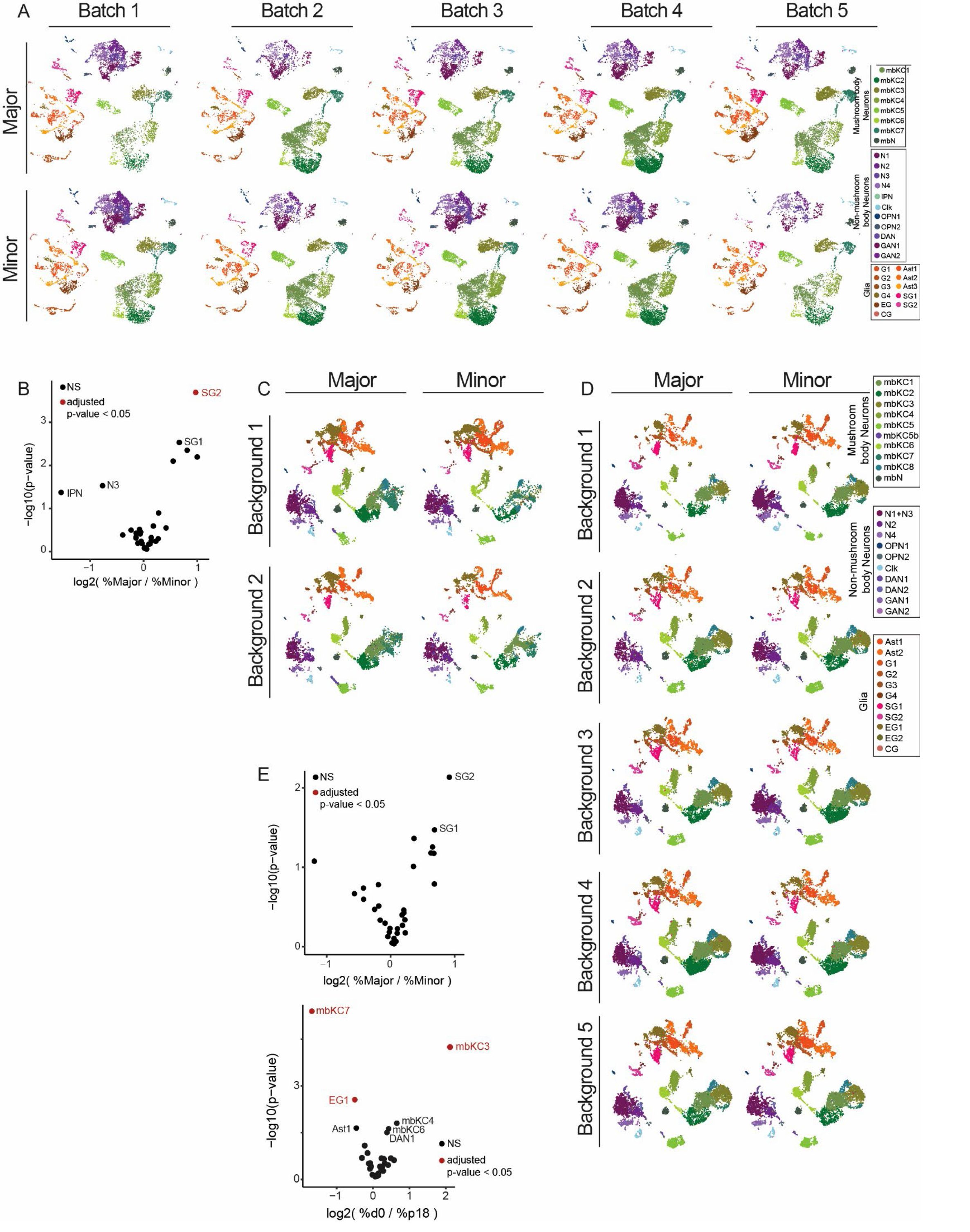
Individual UMAP plots (A, C and D) for d0 (A), and combined (C and D) clustering analyses, showing all replicate representation across celltypes, as well as assessment of differential cluster representation between (B) caste for d0 clustering, as well as (E, upper) caste and (E, lower) stage for combined clustering. Volcano plots were generated using the log2 fold change in average per-replicate cluster percentage (cluster cell numbers / total cell numbers for each replicate) and p-values were generated using a binomial test of cluster representation (see methods), followed by correction for false discovery. Several clusters (mbKC3, mbKC7, N3, EG1) showed significant differential representation, biased to d0 (mbKC3) or p18 (mbKC7 and EG1) samples.

**Fig S9.**
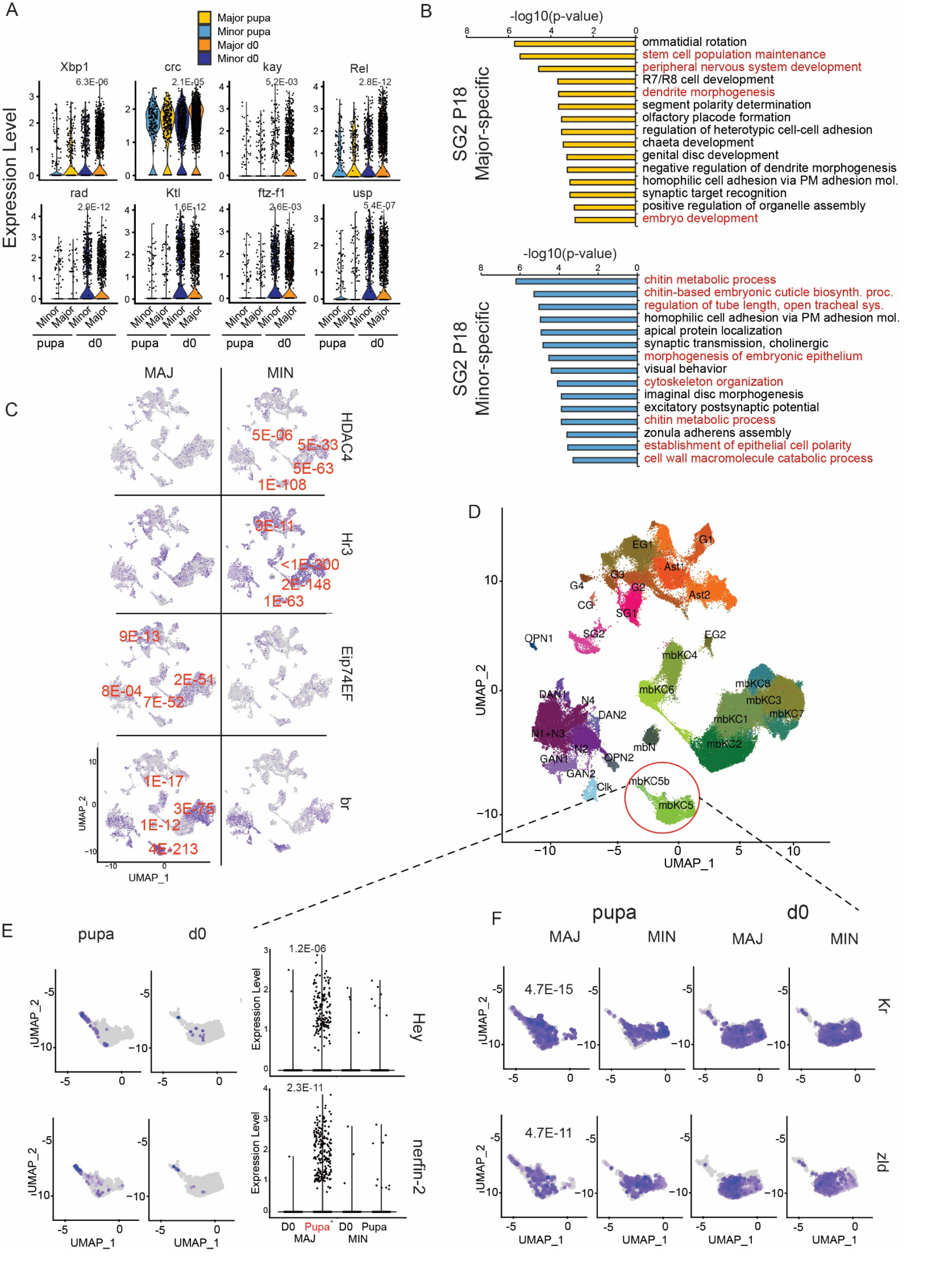
A) Violin plot of d0 caste-biased TFs in SG2 which are very lowly expressed and thus not differing between castes in pupa. P-values represent adjusted p-values from Seurat (Wilcoxon method) comparing Major to Minor for each stage (p18 and d0) and are only presented for significant comparisons (all d0). B) GO term enrichment for DEGs distinguishing Major (top) and Minor (bottom) SG2 cells in pupa. Terms associated with putative SG-specific functions differentiating SG1 and SG2 are highlighted in red. X-axis represents log10- adjusted p-values from GO term enrichment comparing the given SG celltype’s top 200 marker genes to the alternative SG clusters’ markers. C) Three genes showing significant global Minor bias (top row) or Major bias (bottom two rows) in p18 brains plotted on UMAP generated from a combined clustering of all cells from p18+d0 data, but only showing p18 cells. Presented p-values are representative (all given in supplementary Table S10) of the clusters with the largest differenees (by Seurat-generated adjusted-pvalue; wilcoxon method), and presented next to their respective clusters. D) A small cluster of cells in mbKC5 (mbKC5b) showing neuroblast or differentiation-related features and only enriched in pupa Major (E), as well as (F) two TFs implicated in similar roles (*Kr* and *zelda*) seen in both stages. P-values for F given in bottom right corner of pupal Major plots represent significance (adjusted p-value) of caomparing p18 Major to Minor workers using Seurat for cluster SG2.

**Fig S10.**
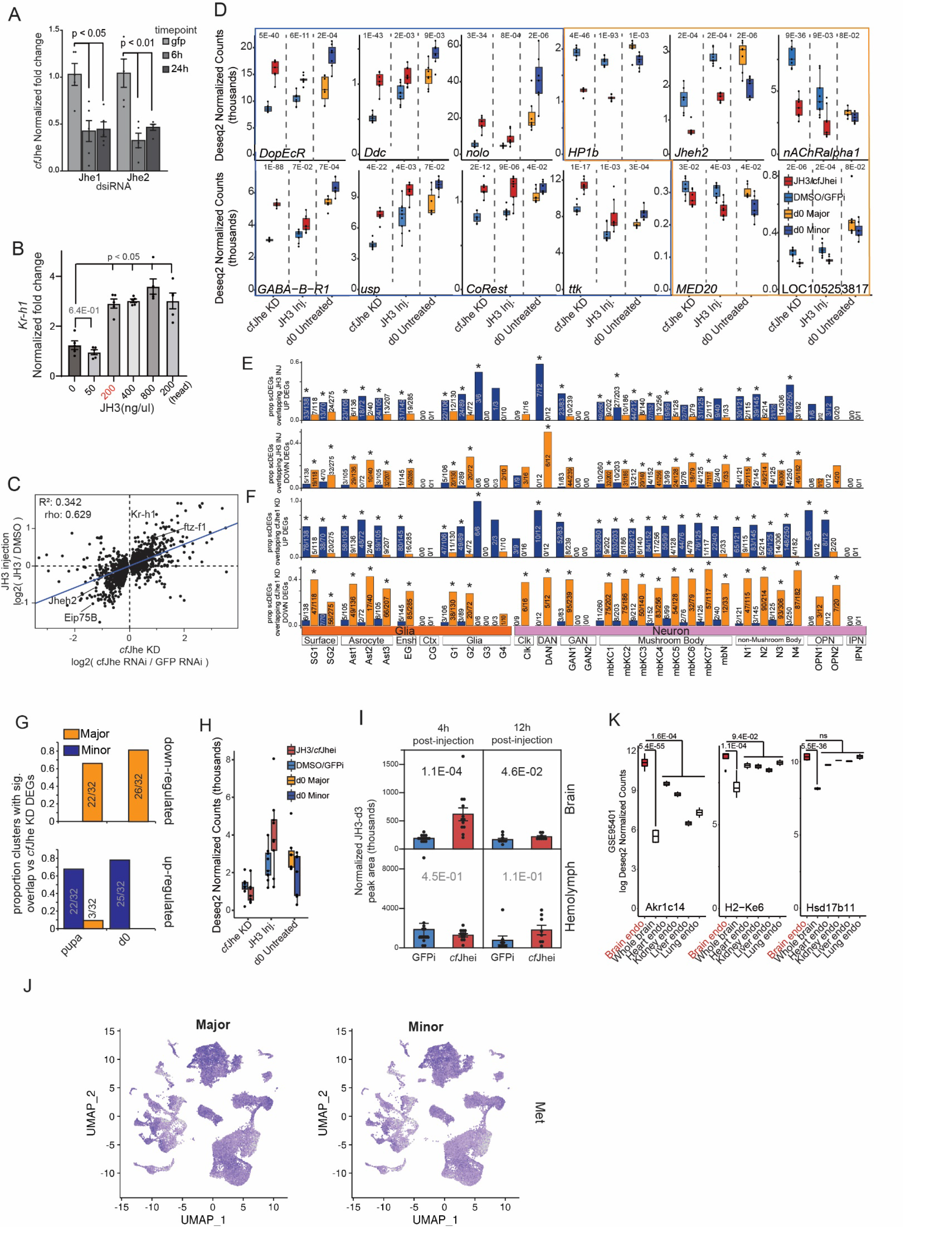
A) RT-qPCR validation of two *cf*Jhe KD dsiRNAs from 6h and 24hs post-injection. KD results in 60% KD for over 24h. such efficient KD (for ants) presumably due to *cf*Jhe’s perineural glia localization (surface). P-values generated via a t-test comparing *cf*Jhe KD to GFP control KD. n =5, P-values generated via a two-sided Kruskal-Wallis tests followed by Dunn’s test. B) RT-qPCR Validation of JH3 injection utilizing Kr-h1 expression (RT-qPCR) as a proxy for JH3 entry into the brain upon gaster injection. 200ng/uL (in 1uL) was chosen as the lowest amount of JH3 injected that still showed a jump in Kr-h1 response. n = 5, P-values generated via Kruskal-Wallis tests followed by Dunn’s test (two-sided). C) Correlation scatterplot between *cf*Jhe KD and JH3 injection transcriptome for all genes differentially expressed upon *cf*Jhe KD (DESeq2 padj < 0.1), illustrating strong concordance between removal of *cf*Jhe barrier and over-loading of brain with JH3. Rho represents test statistic from a spearman’s rank correlation. D) Boxplots of expression (DESeq2 normalized counts) of single-gene expression for bulk RNA-seq related to Fig. 5D from *cf*Jhe KD, JH3 injection, and d0 untreated castes illustrating multiple concordant genes across all three experiments. Plots boxed in blue show significant Minor-baised expression in untreated castes as well as significant bias to JH3 injection and/or *cf*Jhe KD, while those boxed in orange show the opposite (Major-biased expression and repression upon JH3 injection and/or *cf*Jhe KD). P-values represent adjusted p-values from DESeq2. E) Bar graphs showing proportions of significantly caste-biased scDEGs in each cluster overlapping *cf*Jhe KD down-regulated and up- regulated(top and bottom plots, respectively) genes, or (F) JH3 down-regulated and up-regulated (top and bottom plots, respectively) further illustrating preponderance of overlap between Major scDEGs with JH3 down-regulated genes, and Minor scDEGs with JH3 up-regulated genes. Asterisks indicate significant comparisons (fishers exact test p-value < 0.05) of overlap between DEGs from bulk RNA-seq dataset indicated and genes significantly differing between caste within a given cluster (vai Seurat’s Wilcoxon method). G) Bar graphs showing proportions of scDEGs in each cluster overlapping *cf*Jhe repressed (top) and derepressed (bottom) genes utilizing caste-biased scDEGs from d0 and pupal data when clustered together. H) Bulk RNA-seq normalized counts of *cf*Jhe expression level in *cf*Jhe KD experiment (left), JH3 injections (middle) and bulk RNA-seq of untreated d0 Majors and Minors (right) illustrating that injection of JH3 leads to a (nonsignificant due to variance) trend of upregulation of *cf*Jhe while KD does not. I) LC-MS values (normalized peak area) of brain (top) and hemolymph (bottom) JH3-d3 levels, 4h (left) and 12h (right) following injection. n = 12 for 4h, n=8 for 12h, P-values taken from a two-sided Mann-Whitney U test after blocking for colony background. J) UMAP plot with expression of Met plotted for Major and Minor workers separately illustrating brain expression (but lack of bias to specific celltypes) as well as lack of bias between castes in expression. K) Boxplots of expression for three mouse genes implicated-in or known to mediate hormonal degradation that also show mouse brain endothelial bias vs whole brain, as well as (left two) relative to all other endothelial cells examined in (*160*). P-values represent adjusted p-values from DESeq2 comparing the indicated sample types.

**Figure S11.**
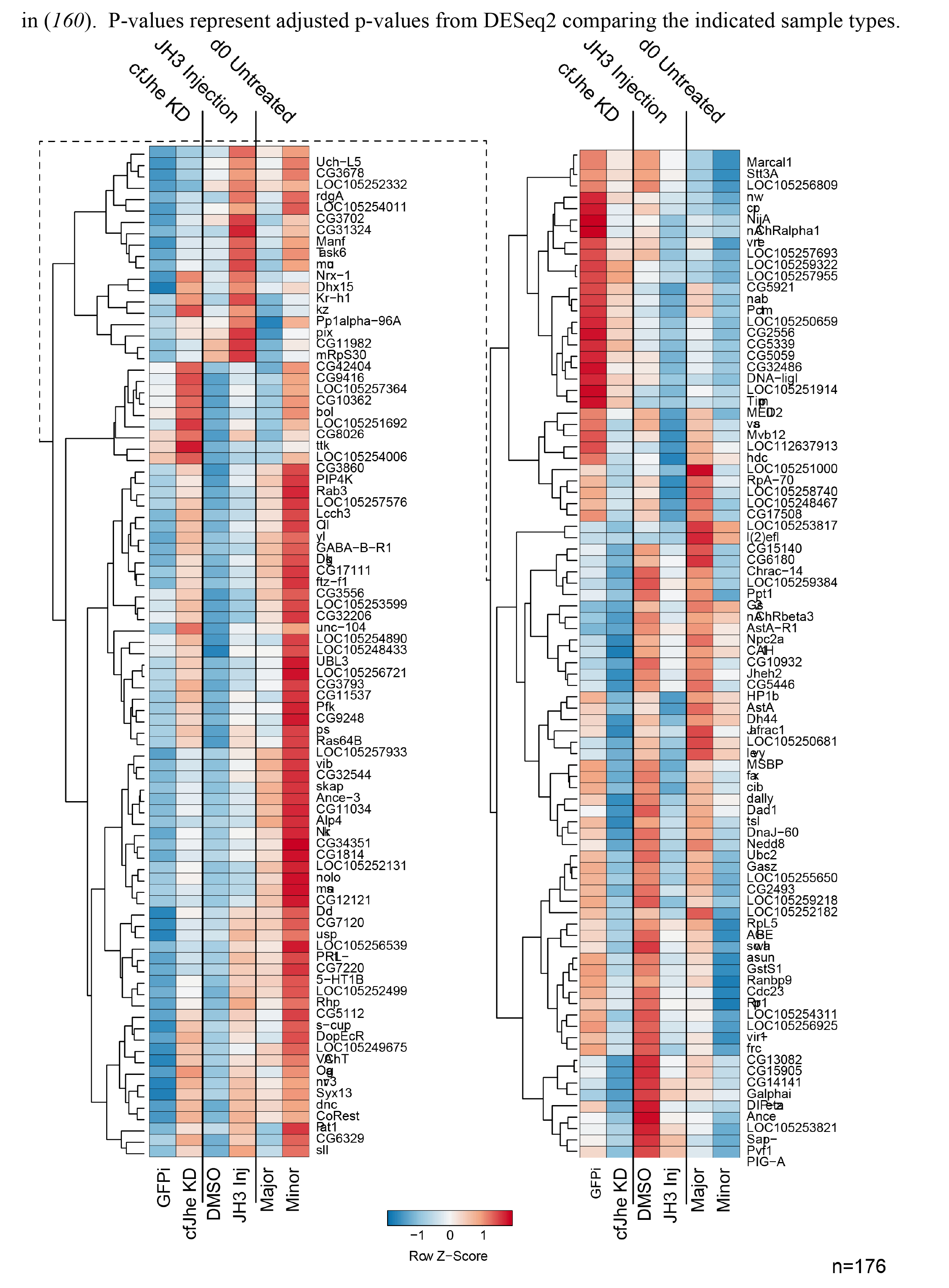
Full set of genes showing consistent directional change upon *cf*Jhe KD, JH3 injection, and untreated Major Minor differences (n=176), related to Figure 5G.

**Fig. S12.**
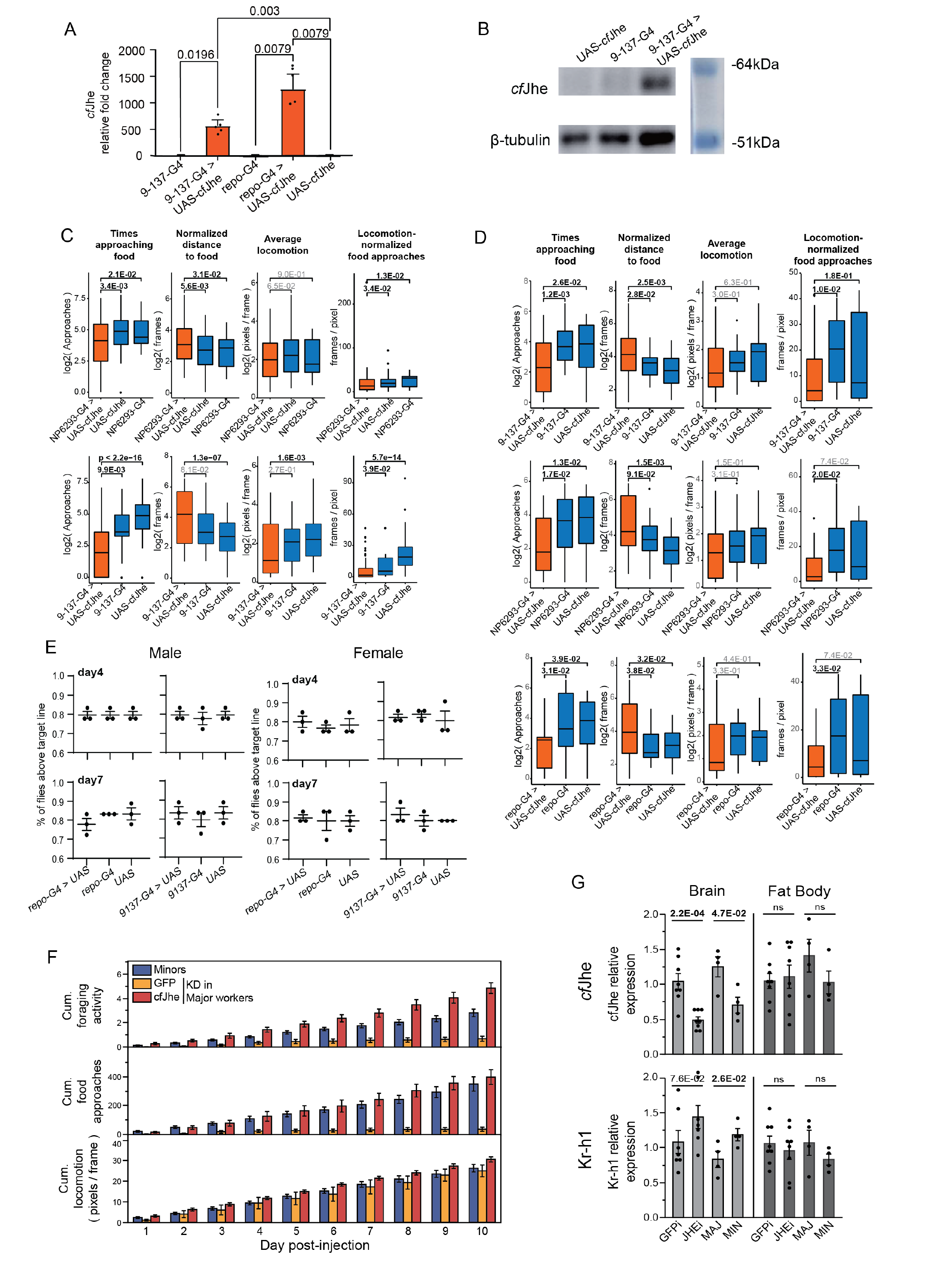
A) *Cf*Jhe RT-qPCR of all genotypes examined in Fig. 6C. n = 5, p-value was generated via Mann-Whitney U test (two-sided) B) Western blot validation of *cf*Jhe overexpression at the protein level, genotypes were the same as Fig. 6C upper. C) Additional fly foraging assay (extension of Fig. 6D) using a secondary surface glia driver (NP6293-Gal4, upper) and a secondary ctrl genetic background (Oregon R, lower), crossing and assaying strategy were identical with Fig.6D. N >20, P-values generated via Kruskal-Wallis tests (two-sided) followed by Dunn’s test. D) Additional fly foraging assay using mated female of genotypes in C) and Fig. 6D. n >20, P-values generated via Kruskal-Wallis tests (two-sided) followed by Dunn’s test . E) Climbing assay using male and female flies of genotypes in Fig. 6D at day4 and day7. Climbing was assessed by counting the percentage of animals climbing above the target line within a fixed amount of time. Each genotype (n=18-20) was repeated for 3 times. F) Individual-day values for automated tracking foraging assay in *C. floridanus* related to Figure 6E showing that multiple days show a significant difference in foraging activity (d5, d6, d7, d9, d10) as well as food interest (d5, d7, d9) between Majors treated with *cf*Jhe-targeting dsiRNA as compared to control injected Majors, as well as similarity of *cf*Jhe KD Majors to untreated Minor workers from the same assay. Majors n=9-10 (*cf*Jhe KD n=10; GFP KD n=9), Minors n=4, Asterisks indicate p<0.05 as determined by a two-sided Mann-Whitney U test comparing *cf*Jhe KD Majors to GFP KD control Majors. G) RT-qPCR results assessing potential systemic impact of *cf*Jhe brain KD via head injection in *C. floridanus* and comparison with untreated Major and Minor workers. *cf*Jhe (KD target) and Kr-h1 (JH3 response gene) mRNA levels showing repeat of KD and Kr-h1 up-regulation in brain (left plots), but no such changes in abdominal fat body (right plots). n = 8 for *cf*Jhe/GFP KD, n = 4 for untreated Major/Minor, P-values from two-sided t-test after assessment of normality using a Shapiro-Wilk test (SW test p-value: 0.3212).

