## Supplemental Figures for "Hormonal gatekeeping via the blood brain barrier governs behavior"

**Supplementary Figures: -**

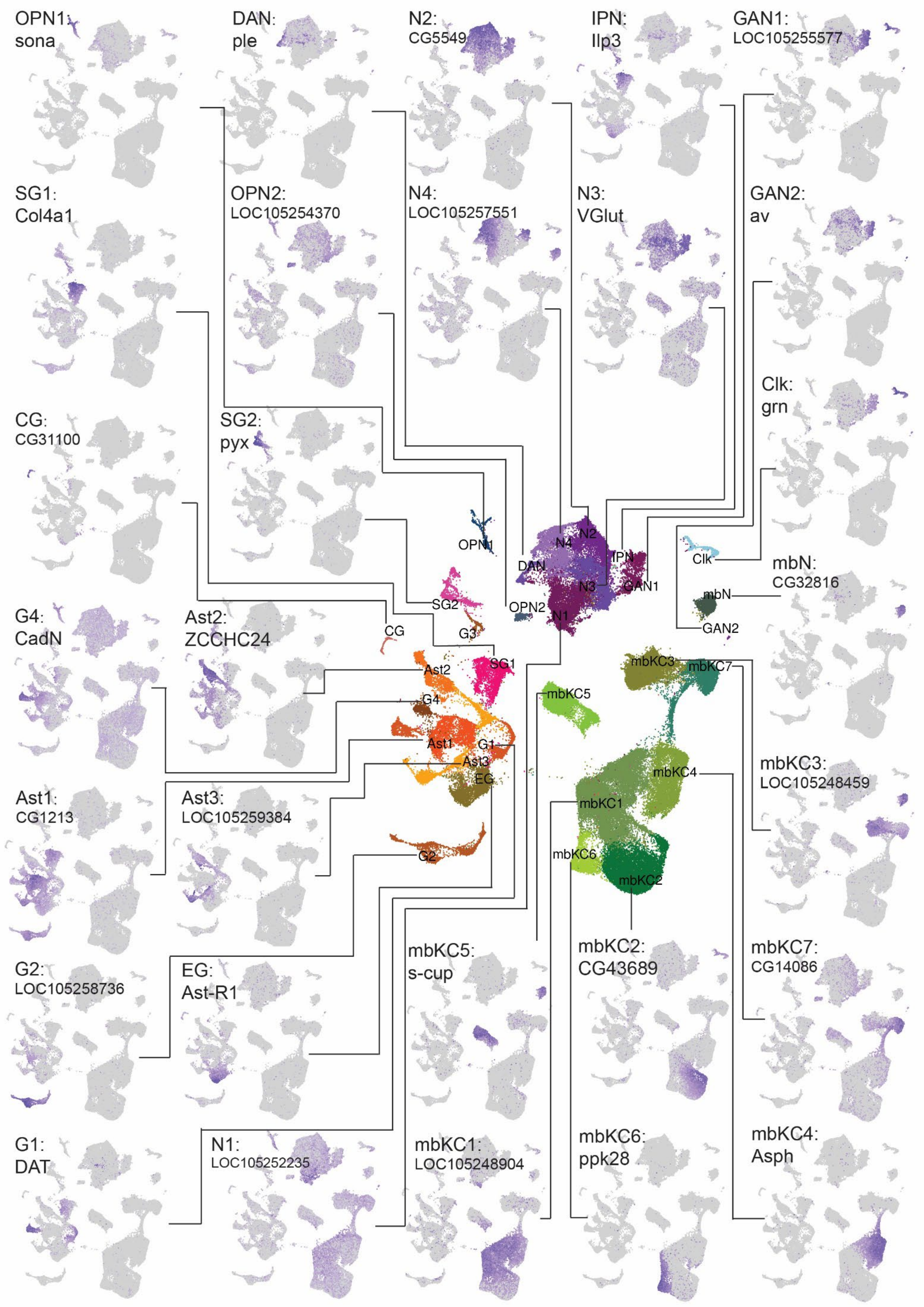

OPN1: sona
DAN: ple
N2: CG5549
IPN: Ilp3
GAN1: LOC105255577
SG1: Col4a1
OPN2: LOC105254370
N4: LOC105257551
N3: VGlut
GAN2: av
Clk: grn
CG: CG31100
SG2: pyx
mbN: CG32816
G4: CadN
Ast2: ZCCHC24
mbKC3: LOC105248459
Ast1: CG1213
Ast3: LOC105259384
G2: LOC105258736
EG: Ast-R1
mbKC5: s-cup
mbKC2: CG43689
mbKC7: CG14086
G1: DAT
N1: LOC105252235
mbKC1: LOC105248904
mbKC6: ppk28
mbKC4: Asph
OPN1
N4
N2
IPN
DAN
Clk
N3
GAN1
SG2
OPN2
N1
mbN
CG
G3
GAN2
SG1
Ast2
mbKC3
mbKC7
G4
mbKC5
Ast1
G1
Ast3
EG
mbKC4
mbKC1
G2
mbKC6
mbKC2

**Figure S1**: top markers for each cluster, illustrated on UMAP plots. For each cluster the top gene marking the given cluster (by TF-IDF in Table S1) was selected and plotted. Lines connect marker UMAPs to overall colored UMAP (center).

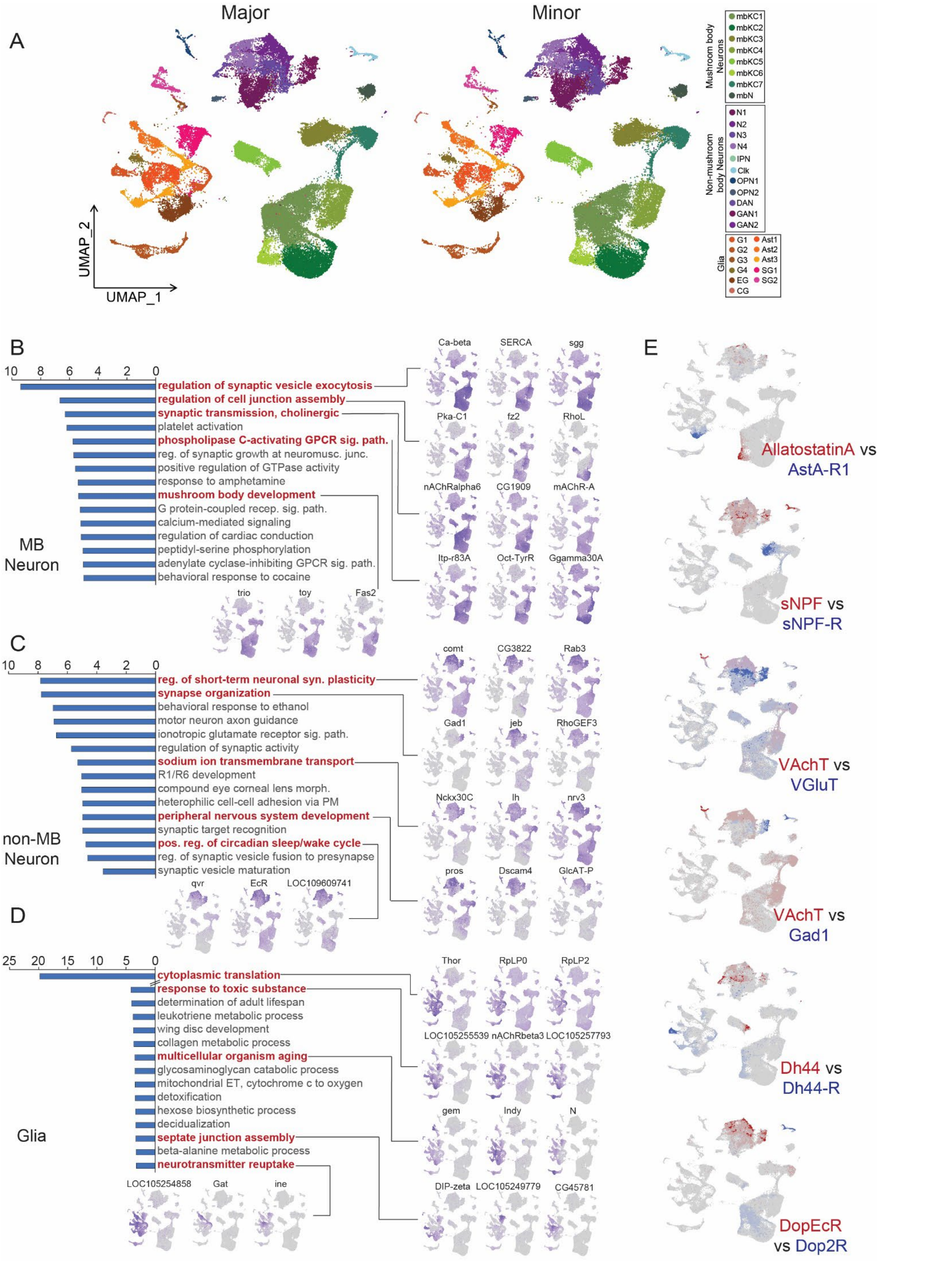

A
Major
Minor
UMAP_2
UMAP_1
Mushroom body Neurons
mbKC1
mbKC2
mbKC3
mbKC4
mbKC5
mbKC6
mbKC7
mbN
Non-mushroom body Neurons
N1
N2
N3
N4
IPN
Clk
OPN1
OPN2
DAN
GAN1
GAN2
Glia
G1
G2
G3
G4
EG
CG
Ast1
Ast2
Ast3
SG1
SG2
B
MB Neuron
regulation of synaptic vesicle exocytosis
regulation of cell junction assembly
synaptic transmission, cholinergic
platelet activation
phospholipase C-activating GPCR sig. path.
reg. of synaptic growth at neuromusc. junc.
positive regulation of GTPase activity
response to amphetamine
mushroom body development
G protein-coupled recep. sig. path.
calcium-mediated signaling
regulation of cardiac conduction
peptidyl-serine phosphorylation
adenylate cyclase-inhibiting GPCR sig. path.
behavioral response to cocaine
Ca-beta
SERCA
sgg
Pka-C1
fz2
RhoL
nAChRalpha6
CG1909
mAChR-A
Itp-r83A
Oct-TyrR
Ggamma30A
trio
toy
Fas2
C
non-MB Neuron
reg. of short-term neuronal syn. plasticity
synapse organization
behavioral response to ethanol
motor neuron axon guidance
ionotropic glutamate receptor sig. path.
regulation of synaptic activity
sodium ion transmembrane transport
R1/R6 development
compound eye corneal lens morph.
heterophilic cell-cell adhesion via PM
peripheral nervous system development
synaptic target recognition
pos. reg. of circadian sleep/wake cycle
reg. of synaptic vesicle fusion to presynapse
synaptic vesicle maturation
comt
CG3822
Rab3
Gad1
jeb
RhoGEF3
Nckx30C
Ih
nrv3
pros
Dscam4
GlcAT-P
qvr
EcR
LOC109609741
D
Glia
cytoplasmic translation
response to toxic substance
determination of adult lifespan
leukotriene metabolic process
wing disc development
collagen metabolic process
multicellular organism aging
glycosaminoglycan catabolic process
mitochondrial ET, cytochrome c to oxygen
detoxification
hexose biosynthetic process
decidualization
septate junction assembly
beta-alanine metabolic process
neurotransmitter reuptake
Thor
RpLP0
RpLP2
LOC105255539
nAChRbeta3
LOC105257793
gem
Indy
N
DIP-zeta
LOC105249779
CG45781
LOC105254858
Gat
ine
E
AllatostatinA vs AstA-R1
sNPF vs sNPF-R
VAchT vs VGluT
VAchT vs Gad1
Dh44 vs Dh44-R
DopEcR vs Dop2R

**Figure S2:** A) UMAP plots of d0 scRNA-seq clustering, showing each caste separately to illustrate generally similar representation in both castes of all major clusters (see Table S7 for full per-sample per-cluster representation). B-D) GO term comparison between MB neuron, non-MB neuron, and glial population marker genes, with informative terms highlighted in red. GO terms represent enriched terms associated with a given cluster's marker genes as compared to a background of the other two clusters' marker genes. X-axis on plots represents log10-transformed p-value of the given term's enrichment among a focal group's marker genes. E) Co-expression of neuropeptide vs receptor (left columns) and X-ergic markers (right, top two), as well as two classes of dopamine receptors (right, bottom panel).

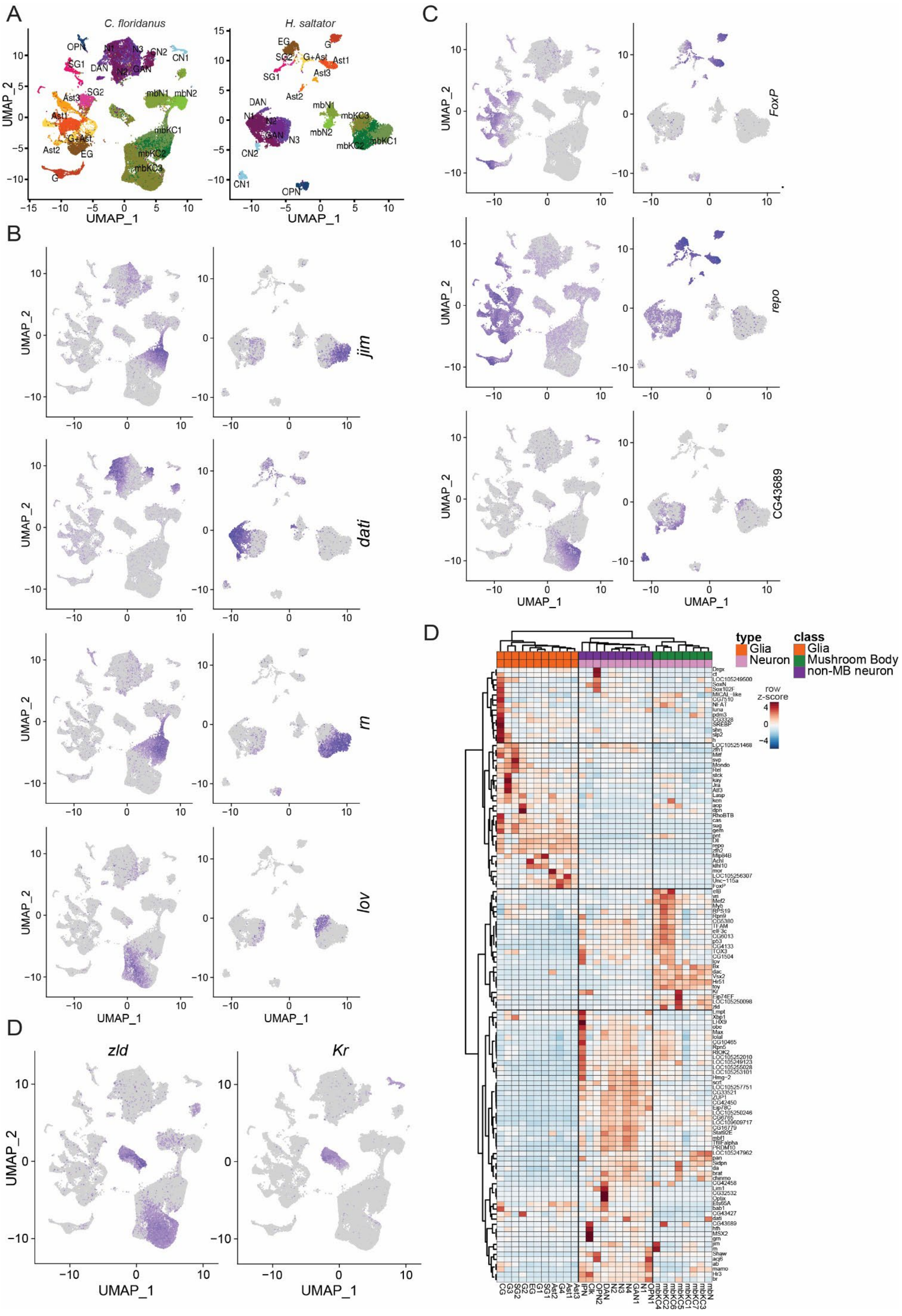
A
C. floridanus
H. saltator
OPN
N1
N3
CN2
CN1
SG1
DAN
N2
GAN
SG2
mbN1
mbN2
Ast3
Ast1
G+Ast
mbKC1
mbKC2
mbKC3
Ast2
EG
G
SG2
G+Ast
Ast3
Ast2
CN1
CN2
OPN
UMAP_1
UMAP_2
B
jim
dati
rn
lov
C
FoxP
repo
CG43689
D
zld
Kr
type
Glia
Neuron
class
Glia
Mushroom Body
non-MB neuron
row z-score

**Fig S3.** A) Cross-species integration between *C. floridanus* and *H. saltator* scRNA datasets illustrating conservation of major cell types between the two species. Clusters in *H. saltator* were annotated independently based upon cluster-specific markers but named based upon orthologous *C. floridanus* clusters, and *C. floridanus* scRNA-seq data was integrated using only orthologous genes between the two species. B) Genes defining distinct neuronal subtypes in *C. floridanus* and *H. saltator* that were also used as integration features between species. C) Additional genes showing strong homologous cluster-specificity between *C. floridanus* and *H. saltator* for major glial subtypes. D) *zld* and *Kr* expression in d0 scRNA-seq showing co-localization to a single cluster (mbKC5). D) More inclusive TF heatmap as from Figure 1F (TF-IDF > 0.5, >1 fold over 2$^{nd}$ best cluster, >2 fold over global levels, >30% of cells in focal cluster expressing TF).

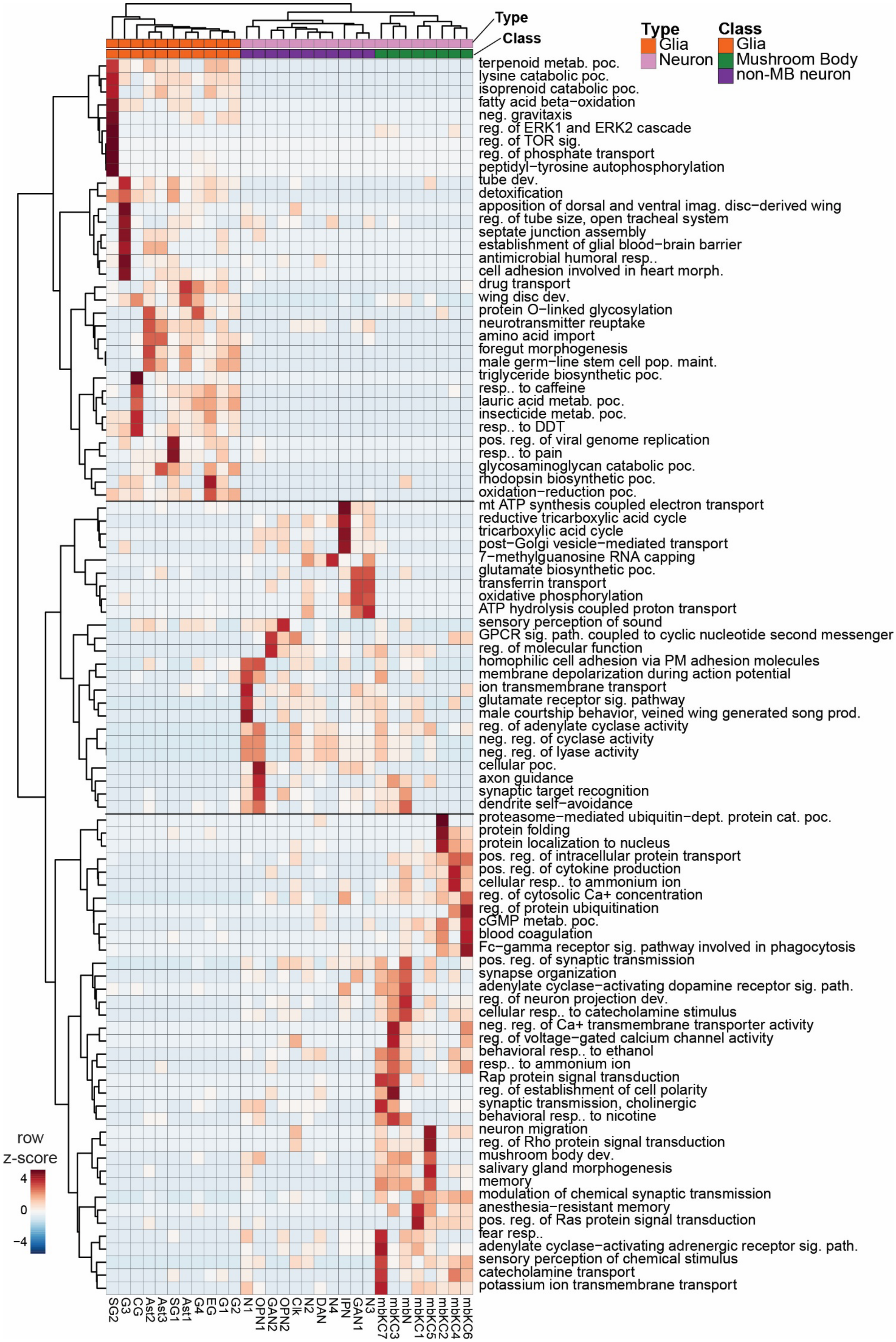
Type
Class
Type
Glia
Neuron
Class
Glia
Mushroom Body
non-MB neuron
row z-score
4
0
−4
terpenoid metab. poc.
lysine catabolic poc.
isoprenoid catabolic poc.
fatty acid beta−oxidation
neg. gravitaxis
reg. of ERK1 and ERK2 cascade
reg. of TOR sig.
reg. of phosphate transport
peptidyl−tyrosine autophosphorylation
tube dev.
detoxification
apposition of dorsal and ventral imag. disc−derived wing
reg. of tube size, open tracheal system
septate junction assembly
establishment of glial blood−brain barrier
antimicrobial humoral resp..
cell adhesion involved in heart morph.
drug transport
wing disc dev.
protein O−linked glycosylation
neurotransmitter reuptake
amino acid import
foregut morphogenesis
male germ−line stem cell pop. maint.
triglyceride biosynthetic poc.
resp.. to caffeine
lauric acid metab. poc.
insecticide metab. poc.
resp.. to DDT
pos. reg. of viral genome replication
resp.. to pain
glycosaminoglycan catabolic poc.
rhodopsin biosynthetic poc.
oxidation−reduction poc.
mt ATP synthesis coupled electron transport
reductive tricarboxylic acid cycle
tricarboxylic acid cycle
post−Golgi vesicle−mediated transport
7−methylguanosine RNA capping
glutamate biosynthetic poc.
transferrin transport
oxidative phosphorylation
ATP hydrolysis coupled proton transport
sensory perception of sound
GPCR sig. path. coupled to cyclic nucleotide second messenger
reg. of molecular function
homophilic cell adhesion via PM adhesion molecules
membrane depolarization during action potential
ion transmembrane transport
glutamate receptor sig. pathway
male courtship behavior, veined wing generated song prod.
reg. of adenylate cyclase activity
neg. reg. of cyclase activity
neg. reg. of lyase activity
cellular poc.
axon guidance
synaptic target recognition
dendrite self−avoidance
proteasome−mediated ubiquitin−dept. protein cat. poc.
protein folding
protein localization to nucleus
pos. reg. of intracellular protein transport
pos. reg. of cytokine production
cellular resp.. to ammonium ion
reg. of cytosolic Ca+ concentration
reg. of protein ubiquitination
cGMP metab. poc.
blood coagulation
Fc−gamma receptor sig. pathway involved in phagocytosis
pos. reg. of synaptic transmission
synapse organization
adenylate cyclase−activating dopamine receptor sig. path.
reg. of neuron projection dev.
cellular resp.. to catecholamine stimulus
neg. reg. of Ca+ transmembrane transporter activity
reg. of voltage−gated calcium channel activity
behavioral resp.. to ethanol
resp.. to ammonium ion
Rap protein signal transduction
reg. of establishment of cell polarity
synaptic transmission, cholinergic
behavioral resp.. to nicotine
neuron migration
reg. of Rho protein signal transduction
mushroom body dev.
salivary gland morphogenesis
memory
modulation of chemical synaptic transmission
anesthesia−resistant memory
pos. reg. of Ras protein signal transduction
fear resp..
adenylate cyclase−activating adrenergic receptor sig. path.
sensory perception of chemical stimulus
catecholamine transport
potassium ion transmembrane transport
SG2
G3
CG
Ast2
Ast3
SG1
Ast1
G4
EG
G1
G2
N1
OPN1
GAN2
OPN2
Clk
N2
DAN
N4
IPN
GAN1
N3
mbKC7
mbKC3
mbN
mbKC1
mbKC5
mbKC2
mbKC4
mbKC6

**Fig S4**. Heatmap of cluster-enriched GO term-based clustering showing segregation of basic main celltypes based entirely on comparison of GO terms, identified by comparing a given cluster's top marker genes to a background set of all genes seen as markers in any cluster. Top marker genes were defined as thte top 200 markers associated with a given cell cluster, or in the case of clusters with less than 200 markers, all.

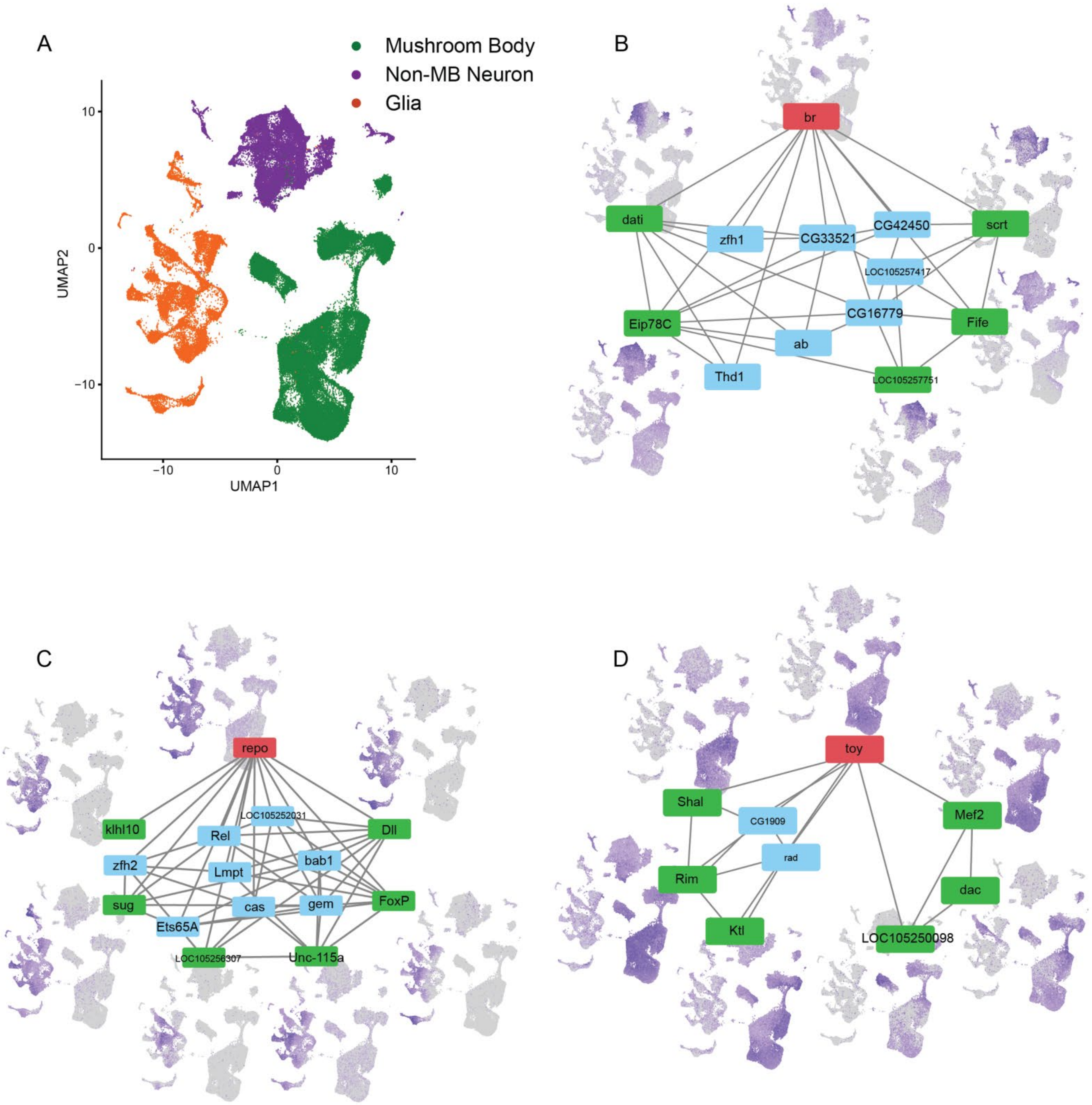
A
Mushroom Body
Non-MB Neuron
Glia
UMAP2
UMAP1
B
br
dati
zfh1
CG33521
CG42450
scrt
LOC105257417
Eip78C
CG16779
Fife
ab
Thd1
LOC105257751
C
repo
LOC105252031
klhl10
Rel
Dll
zfh2
bab1
Lmpt
sug
cas
gem
FoxP
Ets65A
LOC105256307
Unc-115a
D
toy
Shal
CG1909
Mef2
rad
Rim
dac
Ktl
LOC105250098

**Figure S5:** Representative result of gene regulatory analysis on scRNA-seq from *C. floridanus* d0 brain samples. For each major cell group (illustrated in panel A) the top transcription factor regulon is shown as well as its associated TF regulatory network (only showing other TFs downstream of focal TF) are shown, along with UMAP plots of representative downstream TFs showing more sub-cluster specific expression, for B) Mushroom body neurons, C) Glia, and D) Non-mushroom body neurons. The focal parent TF (regulating each network) is shown in red and UMAPs of representative TFs are associated with the green-labeled child nodes. Top regulons were selected by significance of overlap (fishers exact test) between the given major cell group's marker genes (as determined by Seurat's FindMarkers function) and the regulated genes of each regulon.

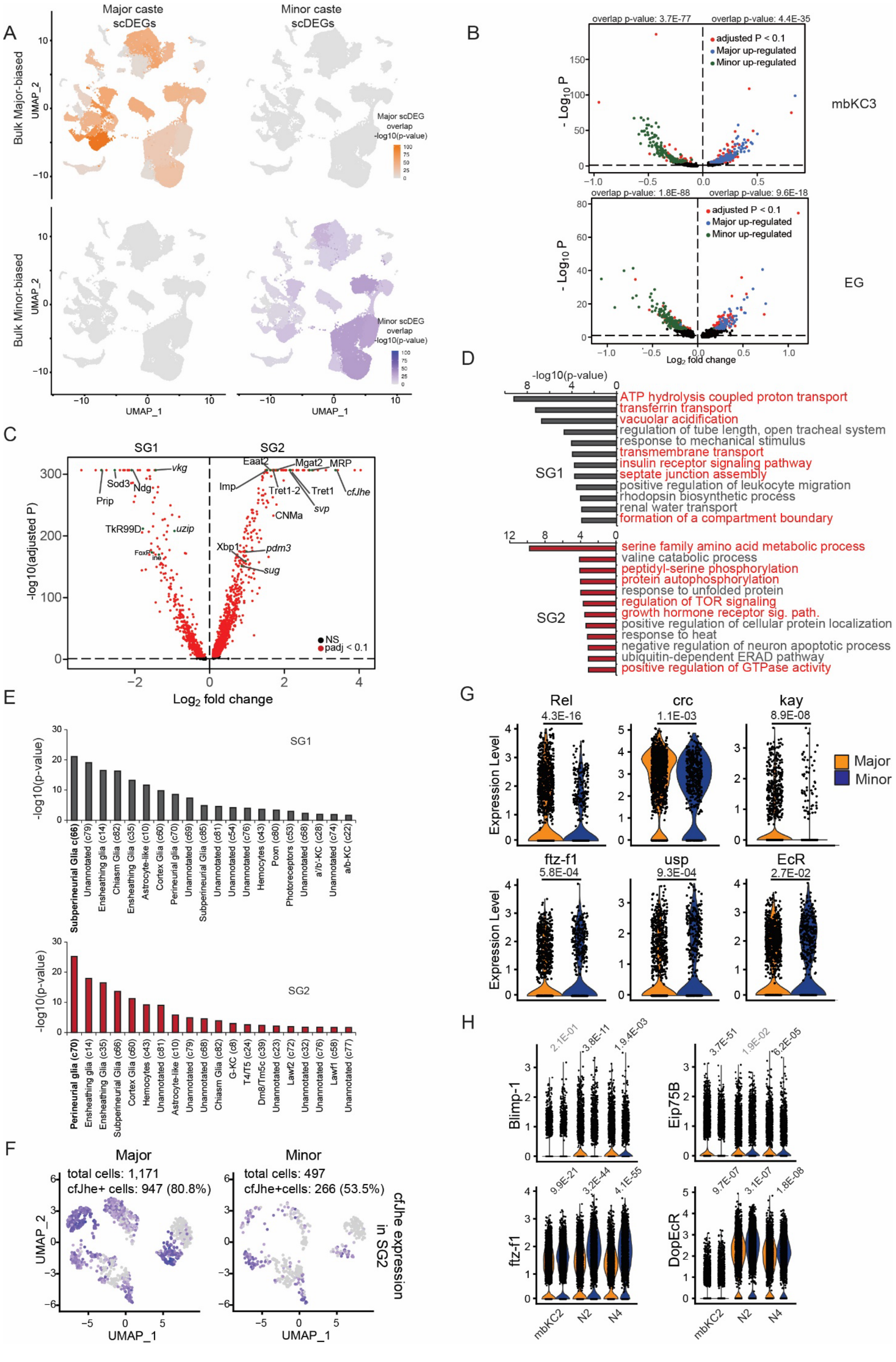
A
Major caste scDEGs
Minor caste scDEGs
Bulk Major-biased
Bulk Minor-biased
UMAP_2
UMAP_1
Major scDEG overlap -log10(p-value)
Minor scDEG overlap -log10(p-value)
B
overlap p-value: 3.7E-77
overlap p-value: 4.4E-35
adjusted P < 0.1
Major up-regulated
Minor up-regulated
mbKC3
overlap p-value: 1.8E-88
overlap p-value: 9.6E-18
EG
- Log10 P
Log2 fold change
C
SG1
SG2
vkg
Sod3
Ndg
Prip
TkR99D
uzip
Eaat2
Mgat2
MRP
Imp
Tret1-2
Tret1
cfJhe
svp
CNMa
Xbp1
pdm3
sug
-log10(adjusted P)
NS
padj < 0.1
Log2 fold change
D
-log10(p-value)
SG1
ATP hydrolysis coupled proton transport
transferrin transport
vacuolar acidification
regulation of tube length, open tracheal system
response to mechanical stimulus
transmembrane transport
insulin receptor signaling pathway
septate junction assembly
positive regulation of leukocyte migration
rhodopsin biosynthetic process
renal water transport
formation of a compartment boundary
SG2
serine family amino acid metabolic process
valine catabolic process
peptidyl-serine phosphorylation
protein autophosphorylation
response to unfolded protein
regulation of TOR signaling
growth hormone receptor sig. path.
positive regulation of cellular protein localization
response to heat
negative regulation of neuron apoptotic process
ubiquitin-dependent ERAD pathway
positive regulation of GTPase activity
E
SG1
-log10(p-value)
Subperineurial Glia c(66)
Unannotated (c79)
Ensheathing glia (c14)
Chiasm Glia (c82)
Ensheathing Glia (c35)
Astrocyte-like (c10)
Cortex Glia (c60)
Perineurial glia (c70)
Unannotated (c69)
Subperineurial Glia (c85)
Unannotated (c81)
Unannotated (c54)
Unannotated (c76)
Hemocytes (c43)
Poxn (c80)
Photoreceptors (c53)
Unannotated (c68)
a'/b'-KC (c28)
Unannotated (c74)
a/b-KC (c22)
SG2
Perineurial glia (c70)
Ensheathing glia (c14)
Ensheathing Glia (c35)
Subperineurial Glia (c66)
Cortex Glia (c60)
Hemocytes (c43)
Unannotated (c81)
Astrocyte-like (c10)
Unannotated (c79)
Unannotated (c68)
Chiasm Glia (c82)
G-KC (c8)
T4/T5 (c24)
Dm8/Tm5c (c39)
Unannotated (c23)
Lawf2 (c72)
Unannotated (c32)
Unannotated (c76)
Lawf1 (c58)
Unannotated (c77)
F
Major
total cells: 1,171
cfJhe+ cells: 947 (80.8%)
Minor
total cells: 497
cfJhe+cells: 266 (53.5%)
cfJhe expression in SG2
UMAP_2
UMAP_1
G
Rel
4.3E-16
crc
1.1E-03
kay
8.9E-08
Major
Minor
ftz-f1
5.8E-04
usp
9.3E-04
EcR
2.7E-02
Expression Level
H
Blimp-1
2.1E-01
3.8E-11
1.9E-03
Eip75B
3.7E-51
1.9E-02
6.2E-05
ftz-f1
9.9E-21
3.2E-44
4.1E-55
DopEcR
9.7E-07
3.1E-07
1.8E-08
mbKC2
N2
N4

**Figure S6:** A) Featureplot showing the overlap of d0 bulk RNA-seq significantly caste biased genes and between-caste scDEGs in each cluster based on the –log10 P-value of Fisher's exact test comparison between bulk DEGs and caste differing scDEGs for each cluster. B) Volcano plot (log2 Major/Minor) of example neuron and glia cluster with strong overlap between bulk DEGs and scDEGs, with overlapping bulk RNA-seq Major-biased DEGs highlighted in blue and Minor-biased DEGs highlighted in green. Caste-biased DEGs for both bulk and scRNA-seq were determined as those with an adjusted p-value < 0.1. C) Volcano plot of DEGs between the two surface glia clusters presented here (log2(SG2 / SG1)), illustrating strong differentiation in gene expression between the two SG clusters. Differentially expressed genes were defined as those showing an adjusted p-value < 0.1 when comparing cells from SG1 to those from SG2, irrespective of caste. D) Top GO-terms associated with genes biased to SG1 (top) or SG2 (bottom) comparing markers of one to a background set of the other, demonstrating functional distinction between the two celltypes markers. Marker genes were defined by comparing SG1 and SG2 cell types, taking those with an adjusted p-value of < 0.01. E) Significance of overlap between ant Surface glia clusters (top: SG2, bottom: SG1) marker genes and cluster-specific markers from all clusters from the fly dataset taken from (*40*). For each fly cluster, marker genes were overlapped with SG-subtype markers. Presented values are the –log10(p-values) of fishers exact tests from these overlaps for the top 20 fly clusters overlapping each ant surface glia subcluster. F) Re-clustering of SG2 cells illustrates higher number of SG2 cells in Major workers, but also much higher percentage of SG2 cells expressing cfJhe independent of cell number differences. G) Violin plot of selected hormonal-signaling related TFs differentially expressed between Major and Minor in SG2. P-values from Seurat differential expression testing (wilcoxon method) comparing Major vs Minor SG2 cells via Seurat. H) Violin plot of two hormonal related TFs differentially expressed between Major and Minor in multiple neuronal clusters (two illustrated here). P-values from Seurat differential expression testing (wilconxon method).

A

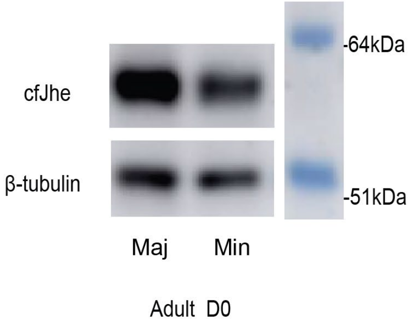


Same membrane from Fig.3C

B

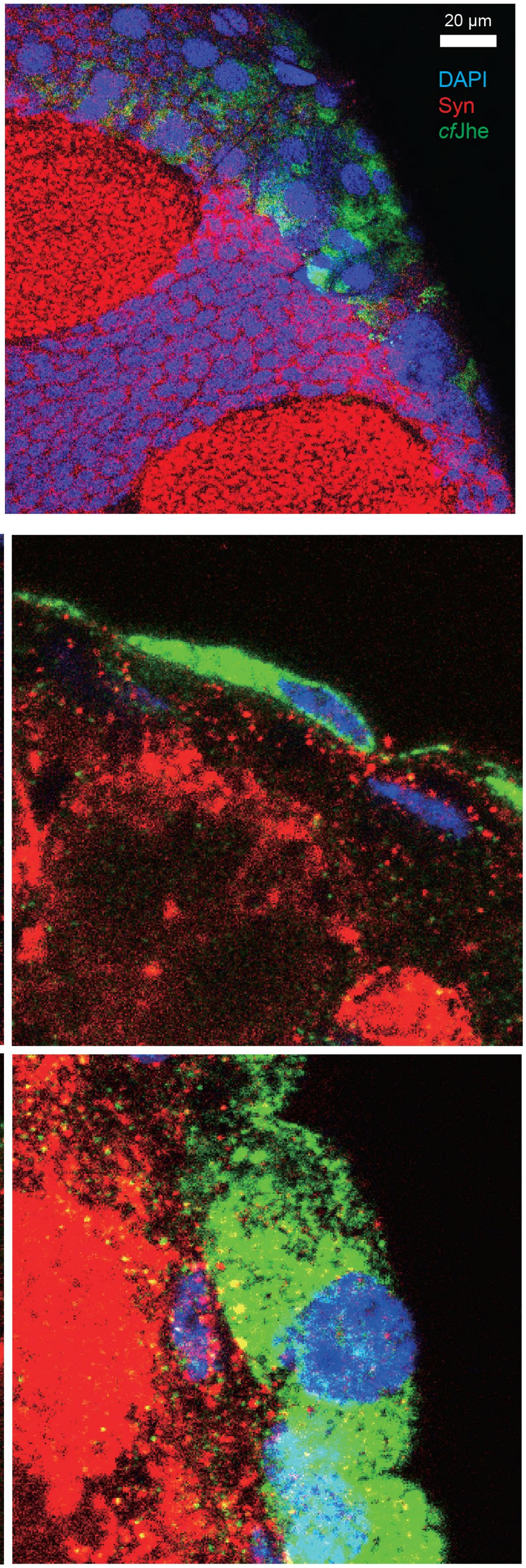

**Fig S7** A) Western blot of *cf*Jhe in d0 Major and Minor brain using a custom antibody raised against a *cf*Jhe-derived antigen. B) Upper: Single confocal slice at top of a 100um Major brain section (*cf*Jhe colocalized with the surface glia nuclei with a typical "large and flat" shape). Lower: 100um Major brain sections imaged under 63X objective illustrating single-cell morphology from different sectioning angles.

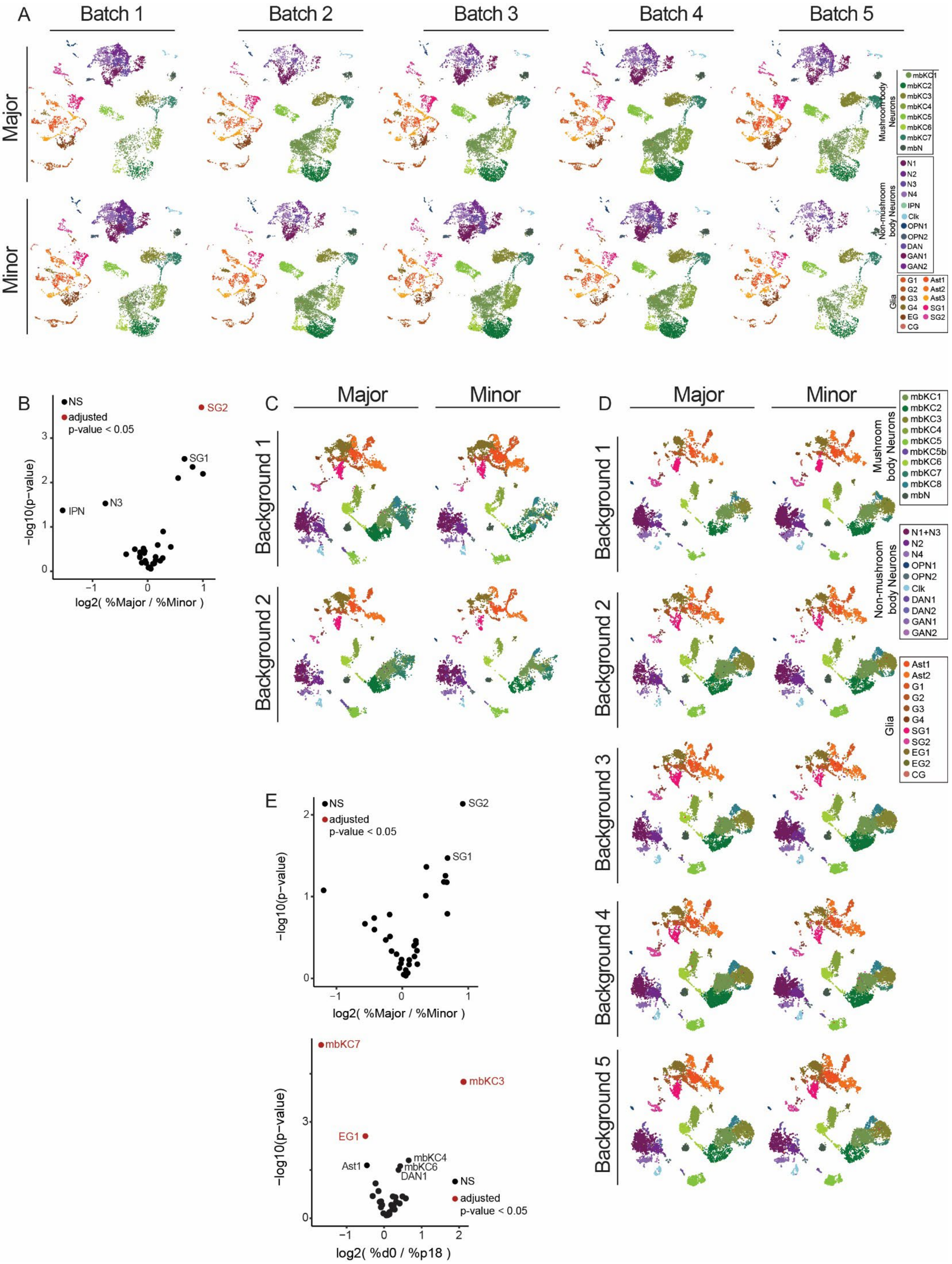
A
Batch 1
Batch 2
Batch 3
Batch 4
Batch 5
Major
Minor
Mushroom body Neurons
mbKC1
mbKC2
mbKC3
mbKC4
mbKC5
mbKC6
mbKC7
mbN
Non-mushroom body Neurons
N1
N2
N3
N4
IPN
Clk
OPN1
OPN2
DAN
GAN1
GAN2
Glia
G1
G2
G3
G4
EG
CG
Ast1
Ast2
Ast3
SG1
SG2
B
NS
adjusted p-value < 0.05
SG2
SG1
N3
IPN
−log10(p−value)
log2( %Major / %Minor )
C
Major
Minor
Background 1
Background 2
D
Major
Minor
Background 1
Background 2
Background 3
Background 4
Background 5
Mushroom body Neurons
mbKC1
mbKC2
mbKC3
mbKC4
mbKC5
mbKC5b
mbKC6
mbKC7
mbKC8
mbN
Non-mushroom body Neurons
N1+N3
N2
N4
OPN1
OPN2
Clk
DAN1
DAN2
GAN1
GAN2
Glia
Ast1
Ast2
G1
G2
G3
G4
SG1
SG2
EG1
EG2
CG
E
NS
adjusted p-value < 0.05
SG2
SG1
−log10(p−value)
log2( %Major / %Minor )
mbKC7
mbKC3
EG1
Ast1
mbKC4
mbKC6
DAN1
NS
adjusted p-value < 0.05
−log10(p−value)
log2( %d0 / %p18 )

**Figure S8.** Individual UMAP plots (A, C and D) for d0 (A), and combined (C and D) clustering analyses, showing all replicate representation across celltypes, as well as assessment of differential cluster representation between (B) caste for d0 clustering, as well as (E, upper) caste and (E, lower) stage for combined clustering. Volcano plots were generated using the log2 fold change in average per-replicate cluster percentage (cluster cell numbers / total cell numbers for each replicate) and p-values were generated using a binomial test of cluster representation (see methods), followed by correction for false discovery. Several clusters (mbKC3, mbKC7, N3, EG1) showed significant differential representation, biased to d0 (mbKC3) or p18 (mbKC7 and EG1) samples.

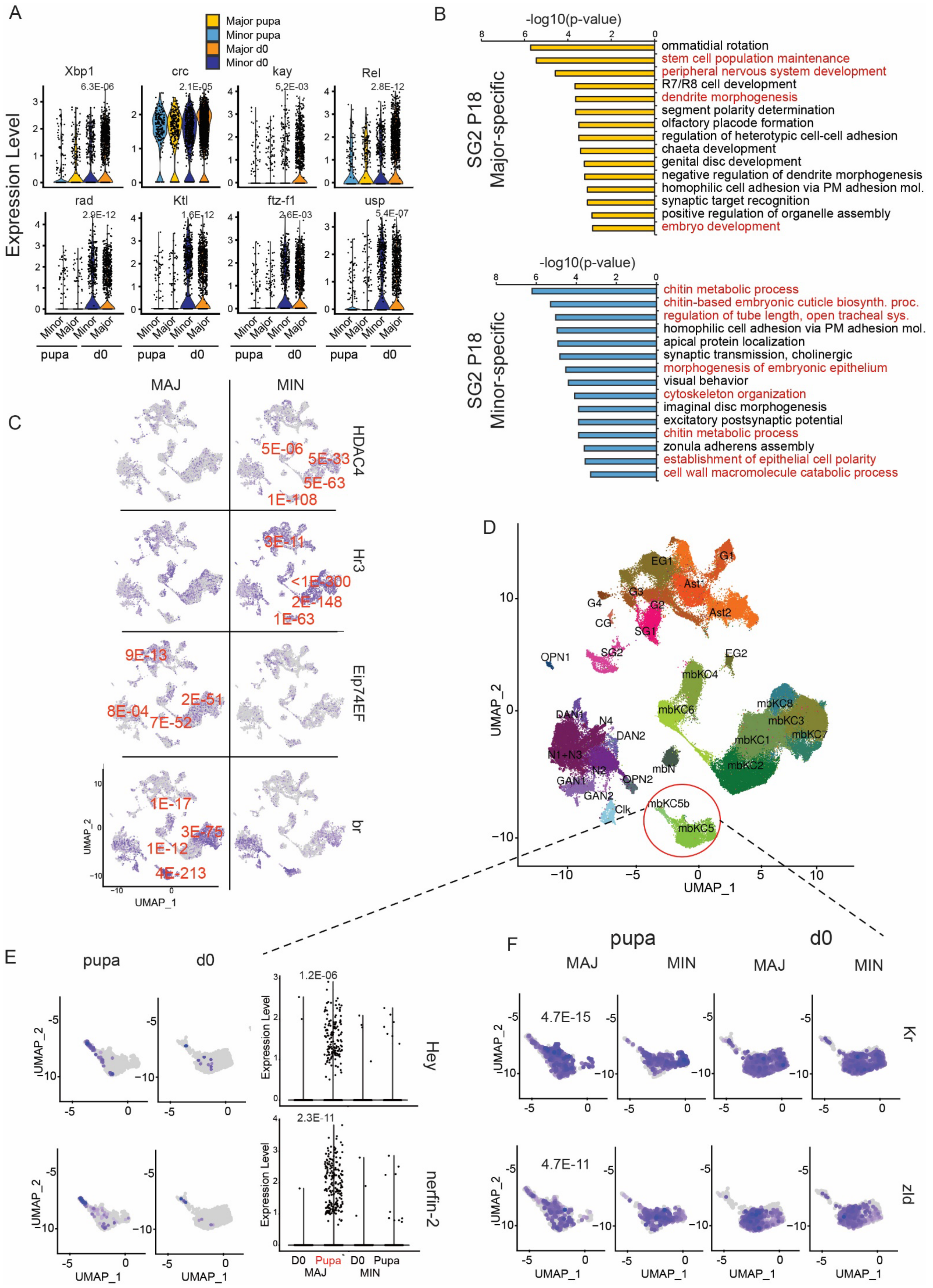
A
Major pupa
Minor pupa
Major d0
Minor d0
Expression Level
Xbp1
crc
kay
Rel
rad
Ktl
ftz-f1
usp
6.3E-06
2.1E-05
5.2E-03
2.8E-12
2.9E-12
1.6E-12
2.6E-03
5.4E-07
Minor
Major
pupa
d0
B
-log10(p-value)
SG2 P18 Major-specific
ommatidial rotation
stem cell population maintenance
peripheral nervous system development
R7/R8 cell development
dendrite morphogenesis
segment polarity determination
olfactory placode formation
regulation of heterotypic cell-cell adhesion
chaeta development
genital disc development
negative regulation of dendrite morphogenesis
homophilic cell adhesion via PM adhesion mol.
synaptic target recognition
positive regulation of organelle assembly
embryo development
SG2 P18 Minor-specific
chitin metabolic process
chitin-based embryonic cuticle biosynth. proc.
regulation of tube length, open tracheal sys.
homophilic cell adhesion via PM adhesion mol.
apical protein localization
synaptic transmission, cholinergic
morphogenesis of embryonic epithelium
visual behavior
cytoskeleton organization
imaginal disc morphogenesis
excitatory postsynaptic potential
chitin metabolic process
zonula adherens assembly
establishment of epithelial cell polarity
cell wall macromolecule catabolic process
C
MAJ
MIN
HDAC4
5E-06
5E-33
5E-63
1E-108
Hr3
3E-11
<1E-300
2E-148
1E-63
Eip74EF
9E-13
2E-51
8E-04
7E-52
br
1E-17
3E-75
1E-12
4E-213
UMAP_1
UMAP_2
D
EG1
G1
Ast1
G4
G3
G2
Ast2
CG
SG1
OPN1
SG2
EG2
mbKC4
mbKC6
mbKC8
mbKC3
mbKC7
mbKC1
DAN1
N4
DAN2
N1+N3
N2
mbKC2
mbN
GAN1
OPN2
GAN2
mbKC5b
Clk
mbKC5
UMAP_1
UMAP_2
E
pupa
d0
Hey
1.2E-06
nerfin-2
2.3E-11
Expression Level
D0
Pupa
MAJ
MIN
UMAP_1
UMAP_2
F
pupa
d0
MAJ
MIN
Kr
4.7E-15
zld
4.7E-11
UMAP_1
UMAP_2

**Fig S9.** A) Violin plot of d0 caste-biased TFs in SG2 which are very lowly expressed and thus not differing between castes in pupa. P-values represent adjusted p-values from Seurat (Wilcoxon method) comparing Major to Minor for each stage (p18 and d0) and are only presented for significant comparisons (all d0). B) GO term enrichment for DEGs distinguishing Major (top) and Minor (bottom) SG2 cells in pupa. Terms associated with putative SG-specific functions differentiating SG1 and SG2 are highlighted in red. X-axis represents log10-adjusted p-values from GO term enrichment comparing the given SG celltype's top 200 marker genes to the alternative SG clusters' markers. C) Three genes showing significant global Minor bias (top row) or Major bias (bottom two rows) in p18 brains plotted on UMAP generated from a combined clustering of all cells from p18+d0 data, but only showing p18 cells. Presented p-values are representative (all given in supplementary Table S10) of the clusters with the largest differenees (by Seurat-generated adjusted-pvalue; wilcoxon method), and presented next to their respective clusters. D) A small cluster of cells in mbKC5 (mbKC5b) showing neuroblast or differentiation-related features and only enriched in pupa Major (E), as well as (F) two TFs implicated in similar roles (*Kr* and *zelda*) seen in both stages. P-values for F given in bottom right corner of pupal Major plots represent significance (adjusted p-value) of caomparing p18 Major to Minor workers using Seurat for cluster SG2.

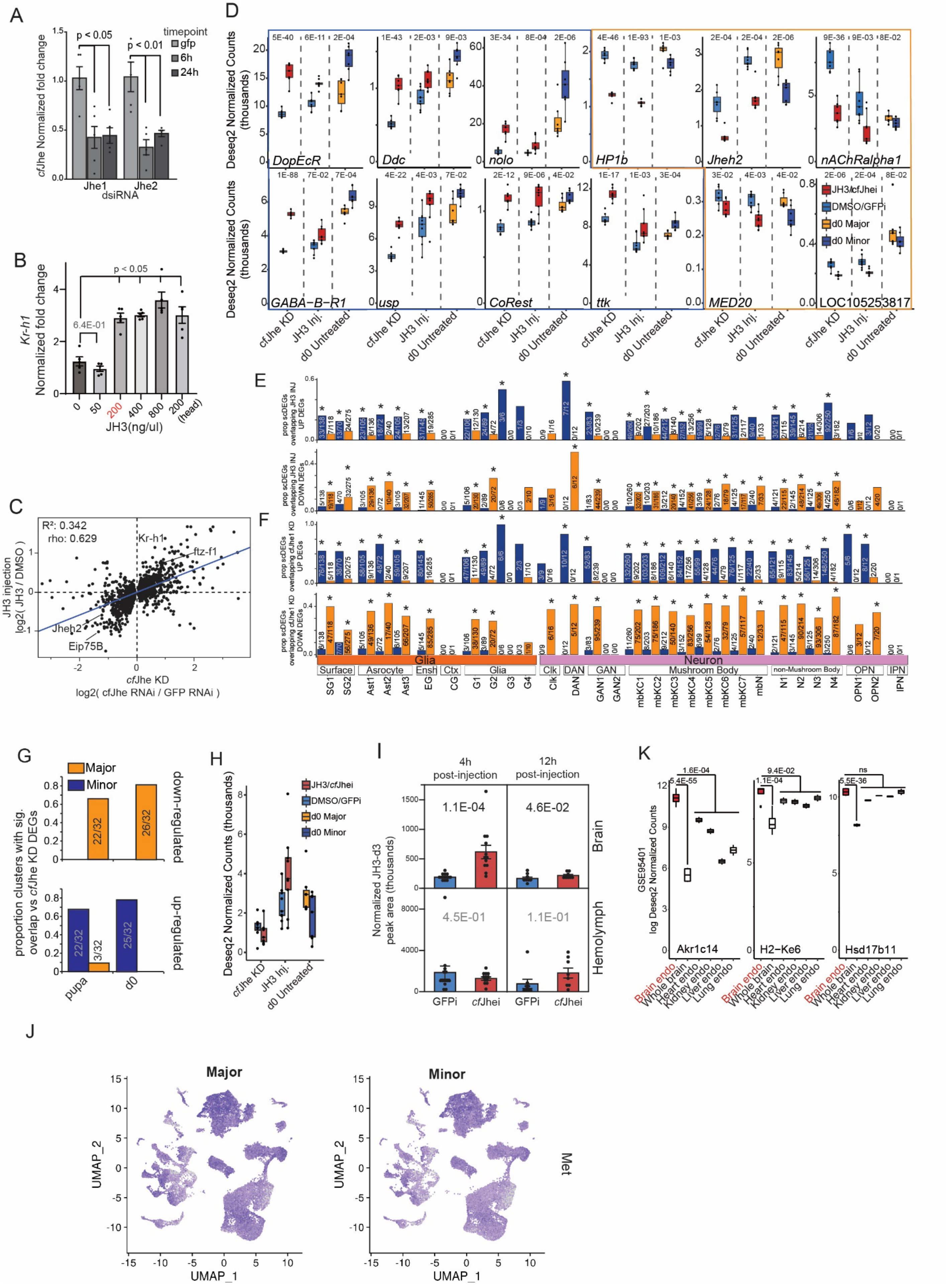

A
B
C
D
E
F
G
H
I
J
K
cfJhe Normalized fold change
p < 0.05
p < 0.01
timepoint
gfp
6h
24h
Jhe1
Jhe2
dsiRNA
Kr-h1
Normalized fold change
6.4E-01
JH3(ng/ul)
(head)
R²: 0.342
rho: 0.629
Kr-h1
ftz-f1
Jheh2
Eip75B
JH3 injection
log2( JH3 / DMSO )
cfJhe KD
log2( cfJhe RNAi / GFP RNAi )
Deseq2 Normalized Counts (thousands)
DopEcR
Ddc
nolo
HP1b
Jheh2
nAChRalpha1
GABA−B−R1
usp
CoRest
ttk
MED20
LOC105253817
JH3/cfJhei
DMSO/GFPi
d0 Major
d0 Minor
cfJhe KD
JH3 Inj.
d0 Untreated
prop scDEGs overlapping JH3 INJ UP DEGs
prop scDEGs overlapping JH3 INJ DOWN DEGs
prop scDEGs overlapping cfJhe1 KD UP DEGs
prop scDEGs overlapping cfJhe1 KD DOWN DEGs
Glia
Neuron
Surface
Asrocyte
Ensh
Ctx
Glia
Clk
DAN
GAN
Mushroom Body
non-Mushroom Body
OPN
IPN
SG1
SG2
Ast1
Ast2
Ast3
EG
CG
G1
G2
G3
G4
Clk
DAN
GAN1
GAN2
mbKC1
mbKC2
mbKC3
mbKC4
mbKC5
mbKC6
mbKC7
mbN
N1
N2
N3
N4
OPN1
OPN2
IPN
Major
Minor
proportion clusters with sig. overlap vs cfJhe KD DEGs
down-regulated
up-regulated
22/32
26/32
22/32
3/32
25/32
pupa
d0
4h post-injection
12h post-injection
1.1E-04
4.6E-02
4.5E-01
1.1E-01
Brain
Hemolymph
Normalized JH3-d3 peak area (thousands)
GFPi
cfJhei
GSE95401
log Deseq2 Normalized Counts
1.6E-04
5.4E-55
9.4E-02
1.1E-04
ns
5.5E-36
Akr1c14
H2−Ke6
Hsd17b11
Brain endo
Whole brain
Heart endo
Kidney endo
Liver endo
Lung endo
UMAP_1
UMAP_2
Met

**Fig S10**. A) RT-qPCR validation of two *cf*Jhe KD dsiRNAs from 6h and 24hs post-injection. KD results in 60% KD for over 24h. such efficient KD (for ants) presumably due to *cf*Jhe's perineural glia localization (surface). P-values generated via a t-test comparing *cf*Jhe KD to GFP control KD. n =5, P-values generated via a two-sided Kruskal-Wallis tests followed by Dunn's test. B) RT-qPCR Validation of JH3 injection utilizing Kr-h1 expression (RT-qPCR) as a proxy for JH3 entry into the brain upon gaster injection. 200ng/uL (in 1uL) was chosen as the lowest amount of JH3 injected that still showed a jump in Kr-h1 response. n = 5, P-values generated via Kruskal-Wallis tests followed by Dunn's test (two-sided). C) Correlation scatterplot between *cf*Jhe KD and JH3 injection transcriptome for all genes differentially expressed upon *cf*Jhe KD (DESeq2 padj < 0.1), illustrating strong concordance between removal of *cf*Jhe barrier and over-loading of brain with JH3. Rho represents test statistic from a spearman's rank correlation. D) Boxplots of expression (DESeq2 normalized counts) of single-gene expression for bulk RNA-seq related to Fig. 5D from *cf*Jhe KD, JH3 injection, and d0 untreated castes illustrating multiple concordant genes across all three experiments. Plots boxed in blue show significant Minor-baised expression in untreated castes as well as significant bias to JH3 injection and/or *cf*Jhe KD, while those boxed in orange show the opposite (Major-biased expression and repression upon JH3 injection and/or *cf*Jhe KD). P-values represent adjusted p-values from DESeq2. E) Bar graphs showing proportions of significantly caste-biased scDEGs in each cluster overlapping *cf*Jhe KD down-regulated and up-regulated(top and bottom plots, respectively) genes, or (F) JH3 down-regulated and up-regulated (top and bottom plots, respectively) further illustrating preponderance of overlap between Major scDEGs with JH3 down-regulated genes, and Minor scDEGs with JH3 up-regulated genes. Asterisks indicate significant comparisons (fishers exact test p-value < 0.05) of overlap between DEGs from bulk RNA-seq dataset indicated and genes significantly differing between caste within a given cluster (vai Seurat's Wilcoxon method). G) Bar graphs showing proportions of scDEGs in each cluster overlapping *cf*Jhe repressed (top) and derepressed (bottom) genes utilizing caste-biased scDEGs from d0 and pupal data when clustered together. H) Bulk RNA-seq normalized counts of *cf*Jhe expression level in *cf*Jhe KD experiment (left), JH3 injections (middle) and bulk RNA-seq of untreated d0 Majors and Minors (right) illustrating that injection of JH3 leads to a (nonsignificant due to variance) trend of upregulation of *cf*Jhe while KD does not. I) LC-MS values (normalized peak area) of brain (top) and hemolymph (bottom) JH3-d3 levels, 4h (left) and 12h (right) following injection. n = 12 for 4h, n=8 for 12h, P-values taken from a two-sided Mann-Whitney U test after blocking for colony background. J) UMAP plot with expression of Met plotted for Major and Minor workers separately illustrating brain expression (but lack of bias to specific celltypes) as well as lack of bias between castes in expression. K) Boxplots of expression for three mouse genes implicated-in or known to mediate hormonal degradation that also show mouse brain endothelial bias vs whole brain, as well as (left two) relative to all other endothelial cells examined in (*160*). P-values represent adjusted p-values from DESeq2 comparing the indicated sample types.

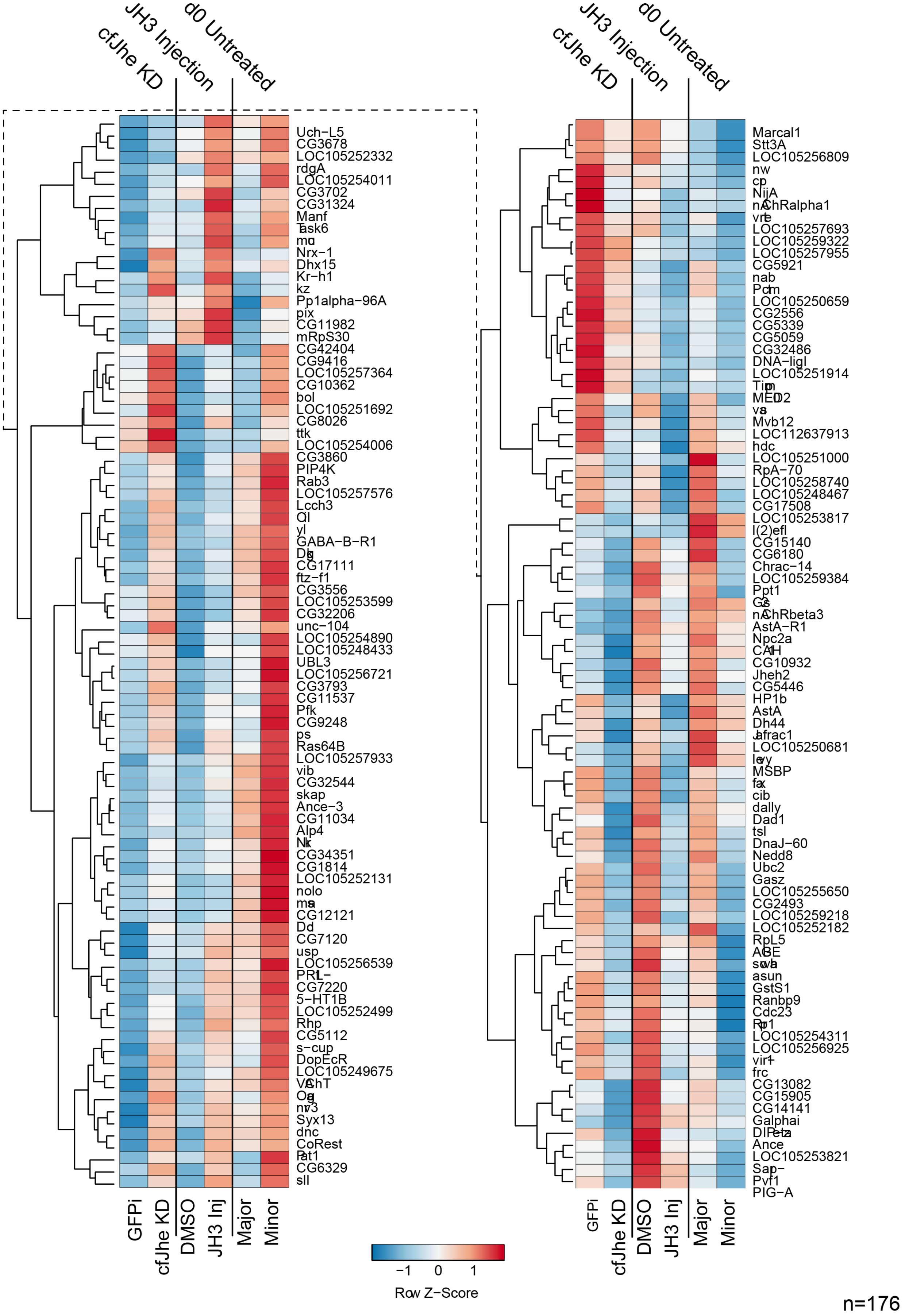

**Figure S11.** Full set of genes showing consistent directional change upon *cf*Jhe KD, JH3 injection, and untreated Major Minor differences (n=176), related to Figure 5G.

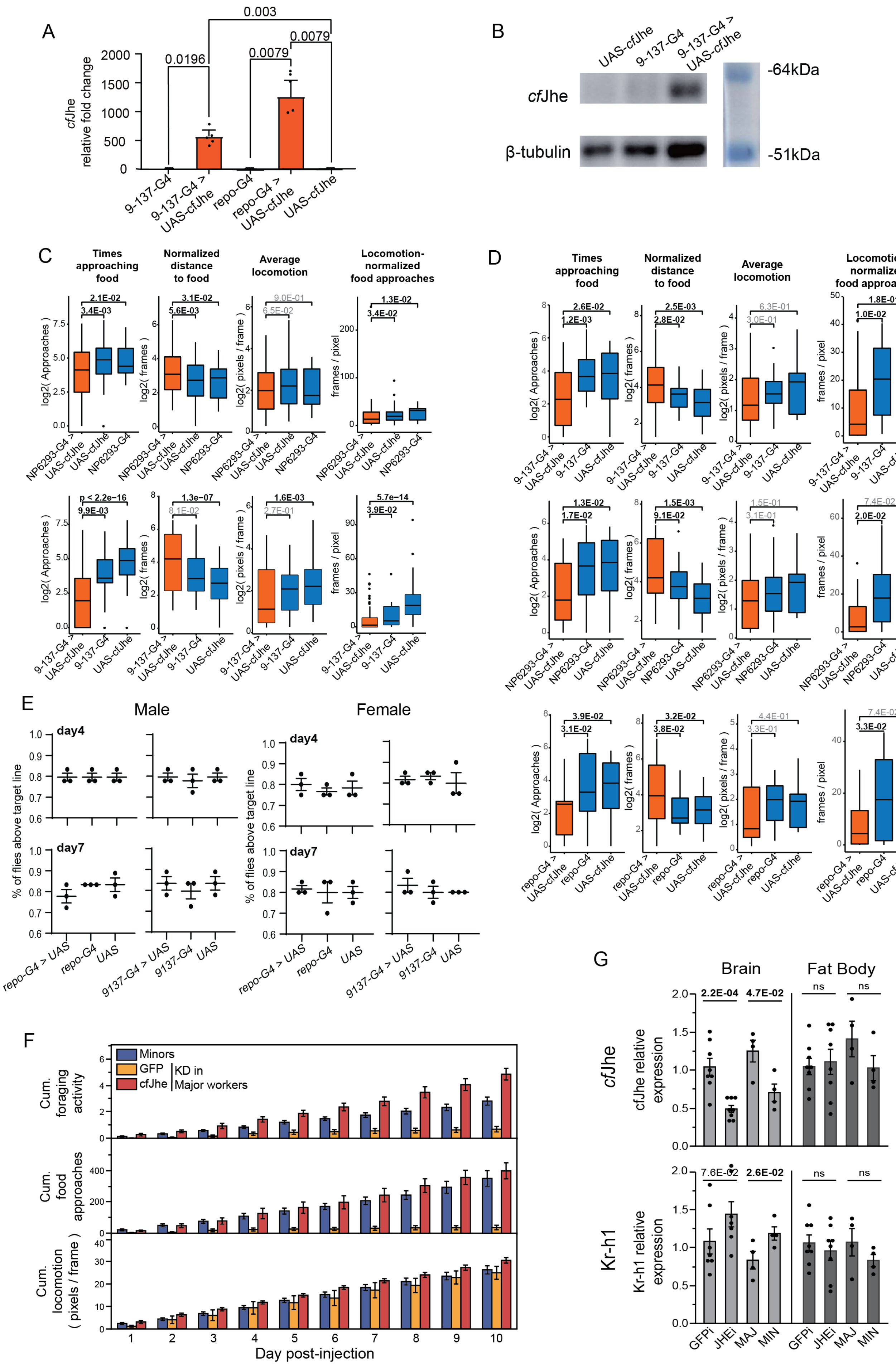

A
0.003
0.0196
0.0079
0.0079
cfJhe relative fold change
2000
1500
1000
500
0
9-137-G4
9-137-G4 > UAS-cfJhe
repo-G4
repo-G4 > UAS-cfJhe
UAS-cfJhe
B
UAS-cfJhe
9-137-G4
9-137-G4 > UAS-cfJhe
cfJhe
β-tubulin
-64kDa
-51kDa
C
Times approaching food
Normalized distance to food
Average locomotion
Locomotion-normalized food approaches
2.1E-02
3.4E-03
3.1E-02
5.6E-03
9.0E-01
6.5E-02
1.3E-02
3.4E-02
log2( Approaches )
log2( frames )
log2( pixels / frame )
frames / pixel
NP6293-G4 > UAS-cfJhe
UAS-cfJhe
NP6293-G4
p < 2.2e-16
9.9E-03
1.3e-07
8.1E-02
1.6E-03
2.7E-01
5.7e-14
3.9E-02
9-137-G4 > UAS-cfJhe
9-137-G4
UAS-cfJhe
D
Times approaching food
Normalized distance to food
Average locomotion
Locomotion-normalized food approaches
2.6E-02
1.2E-03
2.5E-03
2.8E-02
6.3E-01
3.0E-01
1.8E-01
1.0E-02
9-137-G4 > UAS-cfJhe
9-137-G4
UAS-cfJhe
1.3E-02
1.7E-02
1.5E-03
9.1E-02
1.5E-01
3.1E-01
7.4E-02
2.0E-02
NP6293-G4 > UAS-cfJhe
NP6293-G4
UAS-cfJhe
3.9E-02
3.1E-02
3.2E-02
3.8E-02
4.4E-01
3.3E-01
7.4E-02
3.3E-02
repo-G4 > UAS-cfJhe
repo-G4
UAS-cfJhe
E
Male
Female
day4
day7
% of flies above target line
repo-G4 > UAS
repo-G4
UAS
9137-G4 > UAS
9137-G4
UAS
F
Minors
GFP
cfJhe
KD in Major workers
Cum. foraging activity
Cum. food approaches
Cum. locomotion ( pixels / frame )
Day post-injection
G
Brain
Fat Body
2.2E-04
4.7E-02
ns
ns
cfJhe
cfJhe relative expression
7.6E-02
2.6E-02
ns
ns
Kr-h1
Kr-h1 relative expression
GFPi
JHEi
MAJ
MIN
